# Convergent evolution of cluster-wide *Hox* gene regulation in Bilateria

**DOI:** 10.64898/2026.05.18.725834

**Authors:** Billie E. Davies, Francisco M. Martín-Zamora, Tom Frankish, Elise Parey, Nancy Ellis, Noura Maziak, Kero Guynes, Grygoriy Zolotarov, Yi-Jyun Luo, Ferdinand Marletaz, Juan M. Vaquerizas, Arnau Sebé-Pedrós, Nicolae Radu Zabet, Paul J. Hurd, José M. Martín-Durán

**Author notes:** These authors contributed equally to this work (Billie E. Davies, Francisco M. Martín-Zamora). Correspondence: José M. Martín-Durán.

## Abstract

The anteroposterior collinear expression of *Hox* genes is a hallmark of animal development that underpins the diversification of body plans^1^ and life cycles^2^. However, the origin and drivers of this coordinated expression remain elusive: while vertebrates rely on complex cluster-wide *Hox* gene regulation^3–8^, insects define gene-specific, sub-cluster regulatory domains^9–11^. Here, we discover a new mode of *Hox* gene regulation in segmented worms (Annelida). By combining chromatin conformation data with histone modifications profiling in *Owenia fusiformis*, we show that a large distal enhancer forms developmentally regulated, long-range contacts across the Hox cluster, and its activation coincides with the consolidation of a cluster-wide topologically associating domain (TAD), loss of Polycomb-mediated repression, and *Hox* gene transcription. This chromatin structure also occurs in the annelids *Dimorphilus gyrociliatus* and *Capitella teleta*, the latter showing additional subTAD structures that correlate with Hox’s temporal collinearity^12^. Moreover, related phyla with intact, organised Hox clusters and spatial collinearity, such as nemerteans and chitons, show annelid-like chromatin organisations, whereas phyla with disorganised^13^ Hox clusters do not. Coordinated *Hox* gene regulation from a “global control region” is thus ancestral to Lophotrochozoa, indicating that complex regulatory logics based on cluster-wide, long-range chromatin interactions with distal enhancers evolved convergently in vertebrates and spiralians.

## Main

*Hox* genes are a family of homeobox transcription factors (TFs) that play a key role in patterning the anteroposterior axis of bilaterally symmetrical animals and are often physically linked in the genome, forming a gene cluster^1^ (Figure 1a). In many bilaterians, *Hox* genes show coordinated temporal and spatial collinear expression, whereby *Hox* genes near the 3’ end of the cluster are expressed earlier and more anteriorly than those near the 5’ end^13–15^. In vertebrates, this spatial and temporal collinearity is regulated by the dynamic integration, via chromatin extrusion, of the Hox cluster into two flanking topologically associating domains (TADs)^3–5,8^, regions of chromatin with a higher probability of self-interaction^16,17^. This exposes the cluster to distant enhancers that act as global regulators, driving the concerted structure- and tissue-specific expression of *Hox* genes^6,7^ (Figure 1a). Insects, however, rely on fundamentally different principles for *Hox* gene regulation. Insulator regions within the cluster serve as boundary elements, demarcating *cis*-regulatory domains defined by sub-cluster TADs^11,18^ that ensure proper spatial and temporal activation of *Hox* genes during segment patterning and differentiation^9–11^ (Figure 1a). How *Hox* gene activity is controlled in other bilaterian lineages, particularly those that also show coordinated spatial and temporal *Hox* gene expression, remains unknown (Figure 1a), even though understanding this is crucial to elucidating how bilaterian body plans originated and diversified.

**Figure 1.**
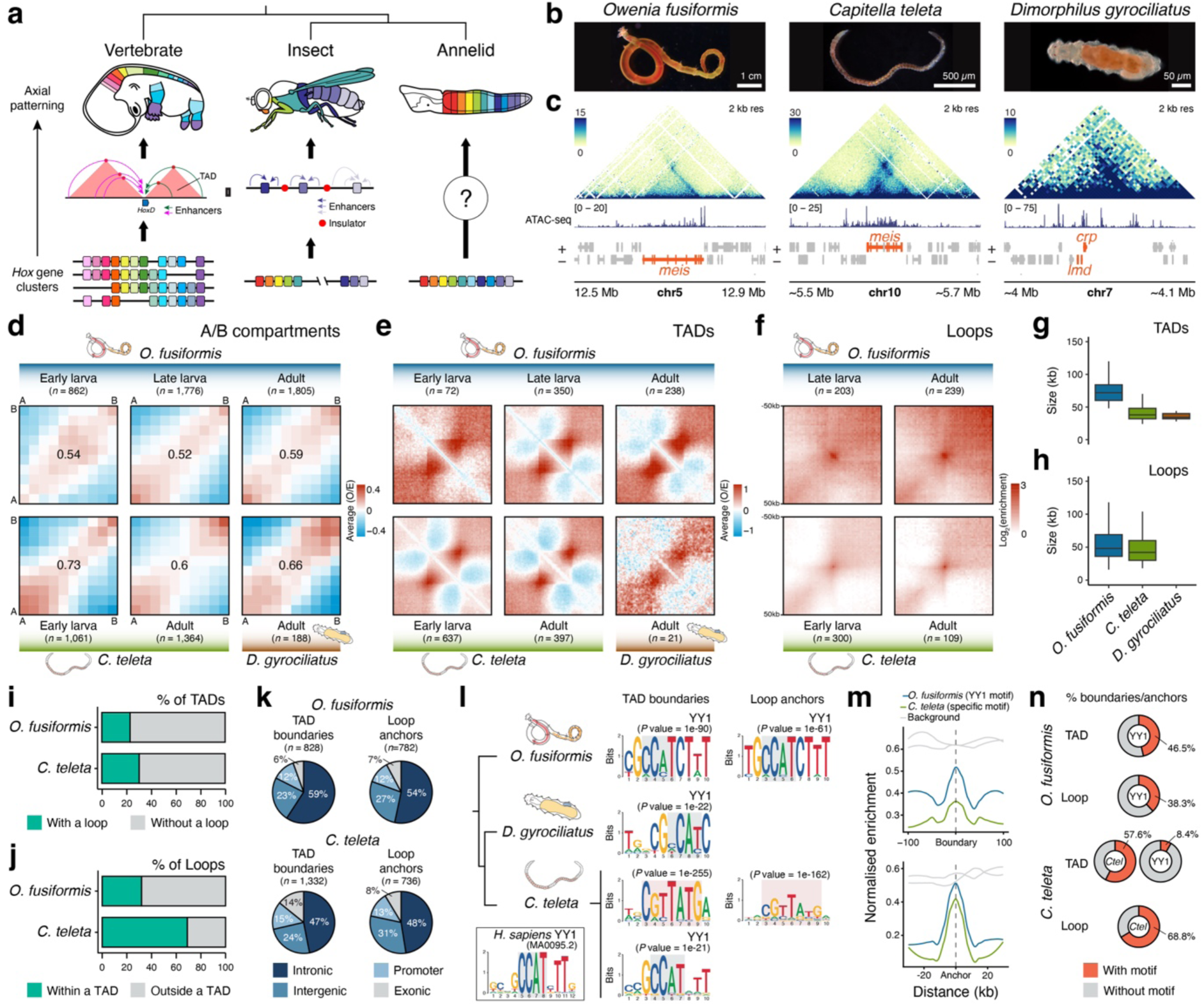
| The annelid 3D chromatin regulatory landscape. (**a**) *Hox* genes are often clustered and pattern the anteroposterior axis of bilaterian animals. However, the genome-regulatory principles controlling *Hox* gene expression differ between vertebrates and insects and are unknown for most other bilaterians, including segmented annelid worms. (**b**) Species studied in this work. Photo credit for *C. teleta*: Prof Elaine Seaver. (**c**) Representative Pico-C contact heatmaps and ATAC-seq tracks around the *meis* locus in *O. fusiformis* and *C. teleta* and *crp* in *D. gyrociliatus*. Heatmaps represent normalised observed counts. (**d**) Saddle heatmaps indicating the strength of A/B compartmentalisation in the three annelids and developmental stages. A compartments are active and gene-rich, while B compartments are more transcriptionally inactive and gene-poor. (**e, f**) Aggregative heatmaps of averaged observed vs expected (O/E) contact frequencies in the called TADs and 50 kb flanking regions (**e**) and contact frequencies around (+– 30 kb) the called loops (**f**). (**g**, **h**) Boxplots indicating the size distribution of consensus TADs (**g**) and loops (**h**) in the three annelid species. (**i**, **j**) Bar plots depicting the proportion of consensus TADs including chromatin loops (**i**) and consensus chromatin loops contained in TADs (**j**) in *O. fusiformis* and *C. teleta*. (**k**) Pie charts indicating the annotation of consensus TAD boundaries and loop anchors in *O. fusiformis* and *C. teleta*. (**l**) In *O. fusiformis* and *D. gyrociliatus*, a motif like that of YY1 in human (bottom left box; JASPAR accession MA0095.2) is enriched in consensus TAD boundaries and loop anchors. A similar motif is also enriched in TAD boundaries in *C. teleta*. However, a species-specific motif without significant similarity to any described motif is highly enriched in TAD boundaries and loop anchors in this species. Grey boxes in YY1 motifs indicate the CCAT core binding site. (**m**) Line plots showing the normalised enrichment of YY1-like motifs and *C. teleta*-specific motifs in consensus TAD boundaries and loop anchors in *O. fusiformis* and *C. teleta*. (**n**) Pie charts depicting the proportion of consensus TAD boundaries and loop anchors containing a YY1-like motif and *C. teleta*-specific motif in *O. fusiformis* and *C. teleta*. Drawings are not to scale.

Differences in the timing of *Hox* gene activation correlate with the evolution of both direct and indirect development, as well as larval types, across phylogenetically distant bilaterians, including annelids, nemerteans, phoronids, hemichordates, and echinoderms^2,19–22^. In these groups, indirectly developing species with larvae that undergo drastic metamorphosis activate *Hox* genes later in their life cycle, after larval hatching and before or during metamorphosis^2,23^. Conversely, species with non-feeding larvae and gradual metamorphosis or direct development activate *Hox* genes during embryogenesis^2^. How these various patterns of *Hox* gene activation are achieved remains unknown, particularly among species within the same phylum that share similar adult body plans but exhibit different life cycles, largely because the genomic principles governing gene expression in these animals are poorly understood^24^. Therefore, the comparative study of such species offers an ideal natural experiment to explore the principles of genome regulation and to determine whether homologous regulatory mechanisms underpin the roles of *Hox* genes in anteroposterior patterning and the diversification of larval forms in bilaterians.

To identify the regulatory mechanisms underlying the different patterns of *Hox* gene activity across species with varying life cycles, we focused on three annelid species — *Owenia fusiformis*, *Capitella teleta*, and *Dimorphilus gyrociliatus* — that share *Hox* gene complements and clusters but differ in the timing of *Hox* gene activation^2,12,25^ (Figure 1b; Extended Data Figure 1a, b). While *O. fusiformis* develops into a feeding larva that delays trunk formation and *Hox* gene expression until late larval stages, *C. teleta* and *D. gyrociliatus* display non-feeding larvae and direct development, respectively, and activate *Hox* genes after gastrulation^2,12,25^. By dissecting their genome-wide landscapes of 3D chromatin architecture and the dynamics of histone post-translational modifications (hPTMs), we demonstrate that activation of a distal global enhancer encapsulating the Hox cluster within a TAD coincides with the timing of *Hox* gene activation in these species, a pattern reminiscent of the regulatory logic controlling *Hox* gene expression in vertebrates. Notably, the presence of a similar chromatin conformation across the Hox clusters of molluscs and nemerteans, groups phylogenetically close to annelids^26,27^, indicates that shared, ancestral regulatory principles control *Hox* gene expression across major bilaterian lineages.

### The chromatin conformation landscape in annelids

To map genome-wide chromatin contacts in these annelids, we initially generated high-resolution Pico-C^28^ datasets for the adult stages (the three species), as well as for the comparable early larval stages showing differences in *Hox* gene activity between *O. fusiformis* and *C. teleta*, and for the late larval stage of *O. fusiformis* that expresses *Hox* genes (Extended Data Figure 1b–d; Supplementary Figure 1). These datasets also helped improve the contiguity of the genome assemblies of *C. teleta* and *D. gyrociliatus* (Supplementary Figure 2). All three species exhibited distinct and dynamic fine-scale 3D chromatin conformations, sometimes involving orthologous genes (Figure 1c; Extended Data Figure 1e, f), suggesting that shared principles of genome regulation operate in these annelids despite differences in genome features such as sizes, transposable element (TE) landscapes, and global 5-methylcytosine levels (Extended Data Figure 1a).

Chromatin segregation into active and inactive domains (A/B compartments) is observed in nearly all animals studied and in their unicellular relatives^29^. Consistently, these three annelids exhibited A/B compartmentalisation (Figure 1d; Extended Data Figure 2a; Supplementary Figures 3–8), with A compartments being gene-rich, TE-depleted, and more transcriptionally active, showing higher coverage of open chromatin regions and gene body methylation levels than B compartments (Extended Data Figure 2b–i; Supplementary Figure 9, 10a). Interestingly, there were fewer, larger, and weaker chromatin compartments in the early larva of *O. fusiformis* (Figure 1d; Extended Data Figure 2a). Accordingly, extensive chromatin reorganisation affecting 41.6% of the genome occurred between this and the late larval stage, compared with more modest changes between late larval and adult stages in this species (26%) and between early larval and adult stages in *C. teleta* (24%) (Extended Data Figure 3a). Most of the compartment changes involved transitions to B compartments, especially during progression to adult stages (Extended Data Figure 3a), and the genes affected were enriched for diverse biological processes (Extended Data Figure 3b, d–f). However, only in *O. fusiformis* did switches to a B compartment result in significantly lower fold changes in expression when the gene was differentially expressed (Extended Data Figure 3c). Therefore, in contrast to previous reports based on low-resolution chromatin conformation data^30^, our findings demonstrate that annelid genomes follow canonical patterns of global chromatin compartmentalisation.

At a finer scale, annelid genomes showed chromatin domains with greater self-insulation, reminiscent of vertebrate and insect TADs (and hence, thereafter referred to as TADs without this implying homology in the underlying molecular mechanisms; Figure 1e), as well as focal points of enriched spatial interactions between distant loci, corresponding to chromatin loops (Figure 1f; Extended Data Figure 1e, f). TADs and loops were larger in *O. fusiformis* than in *C. teleta* and *D. gyrociliatus*, and, therefore, their sizes appeared to positively correlate with total genome size (Figure 1g, h). The identified TADs — predominantly found in A compartments in *O. fusiformis* and *C. teleta* (Extended Data Figure 2k) — covered only a small proportion (8.1% to 14.7%) of the annelid genomes (Extended Data Figure 2j), and accordingly, only a small fraction of genes (5.5% to 13.4%) were contained within these domains that were otherwise gene-rich and TE-depleted (Extended Data Figure 2m–p). About one-third (22.95% and 31.88%) of TADs in *O. fusiformis* and *C. teleta* involved loop formation, yet most loops in *C. teleta* (68.94%), but not in *O. fusiformis* (30.33%), were contained within TADs (Figure 1i, j; Extended Data Figure 2s). As with compartments, the early larva of *O. fusiformis* had fewer, weaker, and overall larger TADs than later stages (Figure 1e, f; Extended Data Figure 2l), and we could not reliably identify any chromatin loop at this stage. This suggests there is a larva- and adult-specific 3D regulatory landscape in *O. fusiformis* that becomes established after larval development (Extended Data Figures 1e, 3g). Collectively, our datasets convincingly demonstrate that annelids exhibit a 3D chromatin organisation comparable to that observed in vertebrates, insects, and other marine bilaterians^31–33^, indicating that an ancestral regulatory logic supports the functioning of bilaterian genomes.

### YY1 motifs are ancestrally enriched in TAD boundaries and loop anchors in annelids

TADs and chromatin loops facilitate long-range regulatory interactions between functional genomic units, such as enhancers, promoters, and insulators^16,34^. In *O. fusiformis* and *C. teleta*, TAD boundaries and loop anchors were mainly found in intronic (47–59%) and intergenic (23–31%) regions (Figure 1k), primarily connecting enhancers with enhancers (intergenic and intronic regions) and enhancers with promoters (Extended Data Figure 4a). To explore the molecular machinery that physically links these regions, we overlapped TAD boundaries and loop anchors with open chromatin data^2^ to identify *cis*-regulatory elements in those regions (Extended Data Figure 4b) and performed *de novo* DNA-binding motif discovery. In *O. fusiformis* and *D. gyrociliatus*, a motif like that of the human YY1 protein, with a conserved CCAT core binding site^35^, was significantly enriched in TAD boundaries and loop anchors (Figure 1l, m). This motif also appeared in *C. teleta* (Figure 1l), but less enriched and only in 8.4% of TAD boundaries, compared with the 46.5% and 38.3% presence of the YY1 motif in *O. fusiformis* boundaries and anchors, respectively (Figure 1n). Conversely, *C. teleta* showed a highly enriched, species-specific motif without any clear similarity to existing profiles in 57.6% and 68.8% of its TAD boundaries and loop anchors (Figure 1l–n). Therefore, YY1, which establishes enhancer-promoter loops in vertebrates^36,37^ and contributes to Polycomb-mediated chromatin looping in *Drosophila melanogaster*^38^, may serve as a core ancestral regulator of TAD and loop formation in annelids. Still, as observed in *D. melanogaster*^39^ (Extended Data Figure 4c), lineage-specific insulators seem to mediate chromatin architecture in the annelid *C. teleta*.

To investigate this divergence in chromatin insulator factors further, we characterised the YY1 proteins in annelids. The three species, like most other invertebrate bilaterians except *D. melanogaster*^40^, have a single YY1 gene (Extended Data Figure 4d). The domain architecture and structure of YY1 in annelids are like their human counterpart, including four DNA-binding C2H2 zinc fingers and a REPO/OPB domain that recruits Polycomb-group (PcG) proteins^41^ (Extended Data Figure 4e–g; Supplementary Figure 11). In *O. fusiformis*, YY1-containing TAD boundaries and loop anchors occurred predominantly between enhancer-to-enhancer interactions (65% and 68.1%, respectively), comparable to the proportion of boundaries and anchors (54.4% and 57.5%, respectively) that mediated enhancer-to-enhancer contacts and contained motifs for the putative *C. teleta*-specific insulator (Extended Data Figure 4h). In *C. teleta*, the TAD boundaries with a YY1-like motif preferentially connected enhancer–enhancer and enhancer–promoter regions (35.9% and 15.8%, respectively) (Extended Data Figure 4i) and were significantly enriched for TFs and developmental genes (Extended Data Figure 4j, k). One such factor was *evx* (*even-skipped* in *D. melanogaster*), which is duplicated in *C. teleta*^42^, with the two loci connected through a pair of YY1-containing downstream insulators in this annelid (Extended Data Figure 4l). Notably, the *even-skipped* locus in *D. melanogaster* is also regulated by a downstream Polycomb- and YY1-dependent insulator^43,44^. Thus, the probable transition to a lineage-specific mechanism of 3D genome regulation in *C. teleta* might have restricted YY1 to a small set of loci with evolutionarily conserved regulatory logics.

### Annelid TADs and loops are enriched in developmental genes

Since the identified TADs and chromatin loops only involve a small fraction of the genome and a limited number of genes, we aimed to investigate the biological roles of genes associated with these higher-order chromatin structures in annelids. Overall, genes within TADs and loops, especially those constitutively included within these structures (Extended Data Figure 3g–k), were enriched for functions related to transcriptional regulation, animal development, cell differentiation, and signalling (Figure 2a). Remarkably, 13.5% and 11.2% of genes in TADs and loop anchors in *O. fusiformis* and *C. teleta*, respectively, possessed a DNA-binding domain and were functionally related to transcriptional regulation and embryogenesis (Extended Data Figure 5b). This represented a significant enrichment compared to the background proportion of such genes in the genome (Figure 2b).

**Figure 2.**
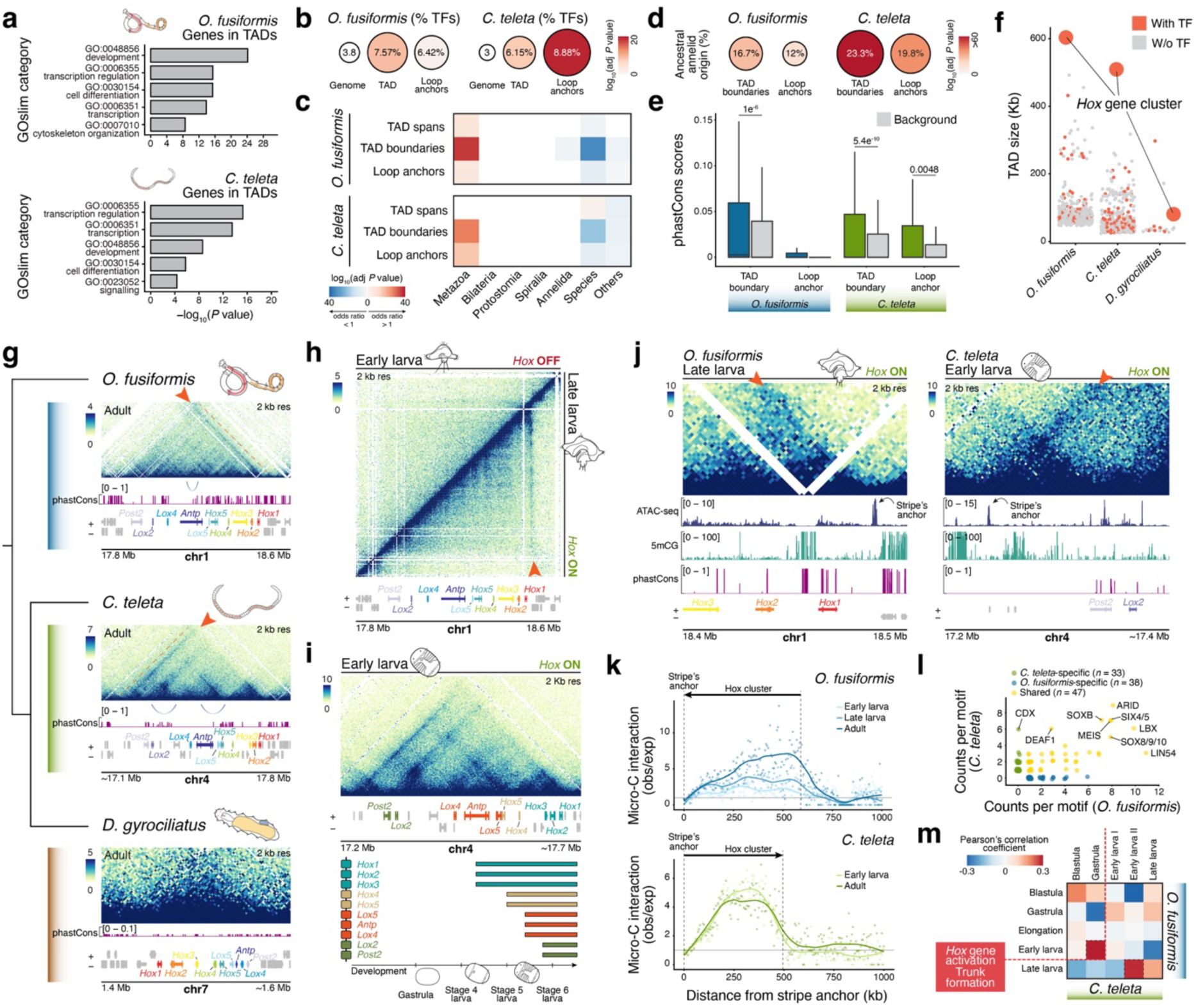
| The 3D chromatin regulation of annelid Hox clusters. (**a**) Gene Ontology slim categories are enriched for biological functions related to development, differentiation and transcriptional regulation in consensus TADs in *O. fusiformis* and *C. teleta*. (**b**) Consensus TADs and loop anchors are enriched for transcription factors in *O. fusiformis* and *C. teleta*. (**c**) Consensus TAD boundaries and loop anchors are enriched for genes of ancestral metazoan origin and depleted of species-specific genes in these two annelids. (**d**) Consensus TADs and loop anchors are enriched for ancestral annelid sequences and exhibit statistically significantly higher sequence conservation than background sequences (**e**). (**f**) The largest TAD in *O. fusiformis* and *C. teleta*, and the third largest TAD in *D. gyrociliatus* encapsulates the Hox cluster. (**g**) Pico-C contact heatmaps of the Hox cluster at the adult stage of *O. fusiformis*, *C. teleta*, and *D. gyrociliatus*. (**h**) Pico-C contact heatmaps of the Hox cluster in the early larva, when *Hox* genes are not expressed, and late larva, when *Hox* genes are expressed, in *O. fusiformis*. Note the establishment of the architectural stripe (red arrowhead) in the late larval stage. (**i**) Pico-C contact heatmap of the Hox cluster in the early larva of *C. teleta* with the *Hox* genes coloured according to their timing of transcriptional activation (bottom panel). (**j**) Pico-C contact heatmaps around the Hox stripe anchor in the late larva of *O. fusiformis* (left) and early larva of *C. teleta* (right). The stripe (red arrowhead) overlaps a peak of accessible chromatin that is low-methylated and shows no overall sequence conservation as indicated by phastCons scores (higher values indicate more sequence conservation). (**k**) Scattered plots of observed vs expected chromatin interactions in 2 Kb bins from the strip anchor along the Hox cluster across stages in *O. fusiformis* and *C. teleta*. Lines are smoothed loess regressions. (**l**) Scattered plots of motif count in the stripe anchors of *O. fusiformis* and *C. teleta*. (**m**) Heatmap depicting the correlation of footprinting binding scores in the Hox stripe anchor during the embryogenesis of *O. fusiformis* and *C. teleta*. The most similar stages are before and after *Hox* gene activation, recapitulating the heterochronic shift in Hox regulation between these two annelids. In (**g**–**j**), heatmaps represent normalised observed counts. In (**a**–**e**), the statistical test is a two-tailed Fisher’s exact test, with Bonferroni correction in (**b**–**e**).

Furthermore, TAD boundaries and loop anchors in both annelids, as well as TAD spans in *O. fusiformis*, were significantly enriched for metazoan-origin genes and depleted for lineage-specific genes (Figure 2c). Indeed, TAD boundaries and loop anchors were also enriched for sequences tracing back to the last common ancestor of annelids, and TAD boundaries in both annelids and loop anchors in *C. teleta* showed significantly higher levels of sequence conservation (Figure 2d, e; Supplementary Figure 12). Collectively, these findings suggest that TADs and chromatin loops involve core metazoan genes and are probably maintained under functional evolutionary constraints in annelids.

In cephalopods, TADs are linked to newly emerged gene microsyntenies, often associated with the morphological innovations found in these molluscs^31^. Phylogenetically distant, *O. fusiformis* and *C. teleta* only share 11 blocks of collinear microsynteny. However, these overlap with TADs in eight (72.7%) and seven (63.6%) cases in *O. fusiformis* and *C. teleta*, respectively (Extended Data Figure 5b, c). Out of all genes located in TADs with a one-to-one orthologous relationship between these two annelids, 168 of them (13.1% in *O. fusiformis* and 5.2% in *C. teleta*) were within a TAD in both species (Extended Data Figure 5d). However, this proportion increased sharply when focusing on TFs and developmental genes, to 31.7% and 16%, respectively (Extended Data Figure 5e). To explore whether pairs of one-to-one orthologs contained in TADs in both *O. fusiformis* and *C. teleta* also shared similar expression domains, we characterised the expression of eight TFs and developmental genes (*meis*, *runx*, *soxB1*, *pax6*, *Notch*, *prospero* [*pros*], *foxA,* and *musashi* [*msi*]) at the larval stages when their TADs are observed (late larvae of *O. fusiformis* and early larvae of *C. teleta*). Seven of these genes (*runx*, *soxB1*, *pax6*, *Notch*, *pros*, *foxA,* and *msi*) had already been characterised during *C. teleta*’s development^42,45–48^, demonstrating a role in establishing the annelid body plan. All these genes exhibited complex, tissue-specific expression domains, often involving multiple larval structures, including the brain (e.g., *soxB1*, *pax6*), nerve cord (*runx*, *soxB1*, *pax6*, *pros*), developing trunk (*meis*, *Notch*, *msi*), and foregut (*foxA*) (Extended Data Figure 5f–m). In all cases, at least one of those domains was conserved between *O. fusiformis* and *C. teleta*. Therefore, although the molecular mechanisms controlling 3D chromatin architecture have diverged between *O. fusiformis* and *C. teleta*, these annelids share a chromatin regulatory logic, particularly around ancestral developmental genes that contribute to the establishment of their body plans.

### Hox gene expression correlates with the formation of a TAD in annelids

Consistent with their association with developmental genes, the largest TAD in *O. fusiformis* and *C. teleta* (604 and 510 kb, respectively), and the third largest in *D. gyrociliatus* (80 kb), encapsulated the Hox cluster (Figure 2f, g). Although the dataset resolution was insufficient to resolve the detailed structure of this TAD in the compact genome of *D. gyrociliatus*, it revealed an architectural stripe defining the TAD in *O. fusiformis* and *C. teleta* (Figure 2g). In vertebrates, stripes in contact maps are regarded as signatures of asymmetric chromatin extrusion resulting from shared regulatory elements “scanning” a genomic region for target genes^49,50^. Strikingly, this stripe formed at the 3’ end of the Hox cluster in *O. fusiformis* (just downstream of *Hox1*) but at the 5’ end (upstream of *Post2*) in *C. teleta* (Figure 2g). Notably, the formation of this stripe correlated with the timing of *Hox* gene activation in *O. fusiformis*: the Hox TAD was weakly insulated, with only a few signs of the stripe, in early larvae when *Hox* genes are not expressed^2^, but became more strongly insulated with an obvious stripe covering the entire TAD in late larvae, when all *Hox* genes are spatially collinearly expressed in the developing trunk before metamorphosis^2^ (Figure 2h; Extended Data Figure 6a). Accordingly, the Hox TAD was strongly self-insulated, and the architectural stripe was apparent already in early larvae of *C. teleta*, which expresses all *Hox* genes in a spatially collinear fashion^12^ (Figure 2i). Furthermore, the Hox TAD in *C. teleta* exhibited subTAD organisation, delineating four areas of higher self-interaction that subdivided the Hox cluster into at least three gene groups (*Hox1*, *Hox2*, and *Hox3*; *Lox5*, *Antp,* and *Lox4*; *Lox2* and *Post2*), aligning with their temporally collinear expression during embryogenesis^12^ (Figure 2i). This subTAD compartmentalisation, however, was not clear in *O. fusiformis* (Figure 2g, h), and, fittingly, this annelid does not show temporal collinear *Hox* gene activation during trunk formation (Extended Data Figure 6b). Therefore, the distinct temporal dynamics of *Hox* gene activity seen in *O. fusiformis* and *C. teleta* correlate with the establishment of a cluster-wide chromatin conformation, albeit with profound differences between these annelids.

As expected from its proposed role in vertebrates^49–52^, the anchor of the architectural stripes in the Hox TADs of *O. fusiformis* and *C. teleta* overlapped with a large (2,310 and 1,719 bp, respectively) region of accessible chromatin (Figure 2j) containing YY1 and the *C. teleta*-specific motifs, respectively. This regulatory element established stronger interactions with the Hox cluster than expected from their physical distance (Figure 2k) and became most accessible in the late larval stage in *O. fusiformis* (when *Hox* genes are active^2^) but during gastrula and early larval stages (the onset of *Hox* gene activation^12^) in *C. teleta* (Extended Data Figure 6c). Although the stripe anchors showed no sequence conservation between the two annelids (Figure 2j), they shared 47 DNA-binding motifs (out of 85 motifs in the stripe, i.e., 55.3%, in *O. fusiformis* and out of 80 motifs in the stripe, i.e., 58.8%, in *C. teleta*) inferred through similarity regression and TF orthology (Figure 2l). Notably, the most abundant shared motifs included TFs expressed in the annelid trunks (e.g., *meis* and *soxB1*; Extended Data Figure 5f, h), *Hox* gene regulators (e.g., DEAF1^53^), segmentation patterning genes (e.g., *lbx*^54^), and components of chromatin remodelling complexes, such as LIN54^55^ and the Trithorax SWI/SNF constituent ARID^56^ (Figure 2l). Many of these were also prevalent in the internal insulators of the Hox TAD in *C. teleta* (Extended Data Figure 6d, e). TF footprinting revealed opposite binding dynamics at the stripe anchor during embryogenesis in these annelids. More TFs, including trunk-related TFs such as *meis*, *runx*, and Notch downstream signalling effectors, were bound, and more strongly, in late larval stages in *O. fusiformis*, coinciding with active *Hox* gene expression (Extended Data Figures 6f–h). Conversely, the peak of binding activity occurred early in development, preceding *Hox* gene activation in *C. teleta* (Extended Data Figures 6f, g). In fact, the stages with the greatest similarity in binding profiles at the stripe anchor were between the early larva of *O. fusiformis* and the gastrula of *C. teleta* (pre-Hox activity stages), and between the late larva of *O. fusiformis* and the early larva of *C. teleta* (the earliest stage of Hox activity) (Figure 2m), recapitulating the heterochronic shift in *Hox* gene expression observed between these two annelids^2^. Thus, the consolidation of a TAD and the transcriptional activation of *Hox* genes coincide with the opening of a distant regulatory element that establishes physical contacts along the entire cluster, reminiscent of the “global control regions” in vertebrate Hox clusters.

### Histone modification dynamics during O. fusiformis development

In vertebrates and insects, *Hox* gene activation involves the removal of Polycomb-mediated H3K27me3, a repressive histone modification^57,58^. To investigate the interplay between the “histone code” and the regulation of *Hox* genes, we focused on *O. fusiformis* because the delay in *Hox* gene activation until late larval stages provides a uniquely tractable system for studying epigenomic dynamics before and after *Hox* gene activity. We generated CUT&Tag libraries to profile, at base resolution, four histone post-translational modifications (hPTMs) — H3K4me3, H3K4me1, H3K27me3, and H3K27ac — across four developmental stages, including the blastula, gastrula, early larva (pre-*Hox* gene activation), and late larva (post-*Hox* gene activation; Figure 3a; Extended Data Figure 1b, 7a; Supplementary Figures 13–17). This enabled us to identify, genome-wide, active promoters (H3K4me3^+^), Polycomb-repressed chromatin (H3K27me3^+^), poised bivalent domains (H3K4me1^+^, H3K27me3^+^), and active (H3K27ac^+^, H3K4me1^+^) and inactive enhancers (H3K4me1^+^) throughout development^59^.

**Figure 3.**
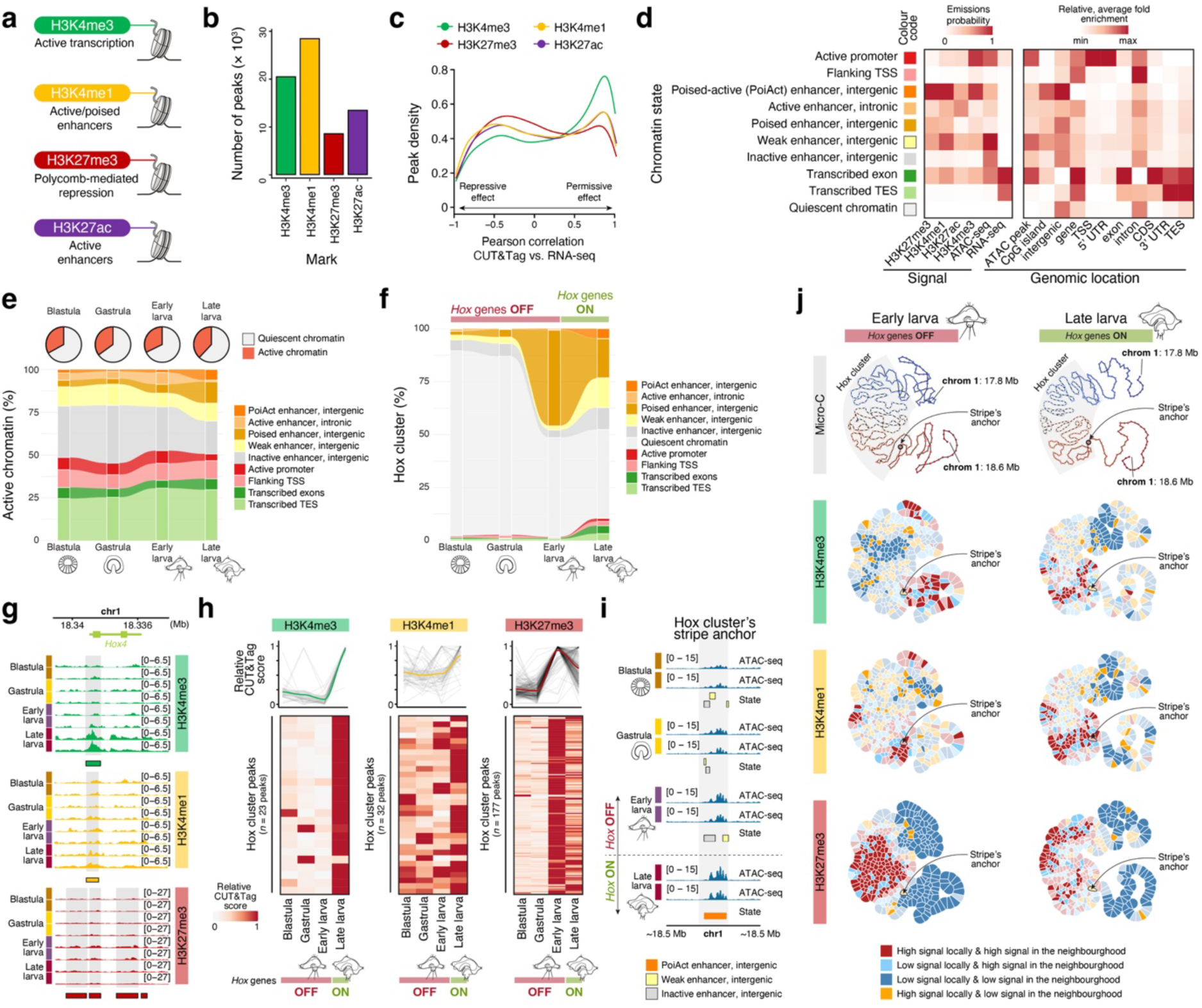
| The histone-based regulation of annelid Hox clusters. (**a**) Histone post-translational modifications profiled during the embryonic and larval development of *O. fusiformis*. (**b**) Bar plots of consensus peak numbers for each histone modification. (**c**) Line plots depicting the correlation between each histone modification and transcription. (**d**) Heatmaps depicting the signal composition (left; based on emissions probabilities) and genomic location (right; based on averaged relative enrichments) of the ten chromatin states inferred from our dataset. (**e**) Chromatin state dynamics during the embryonic and larval development of *O. fusiformis*. On top, the proportion of active and quiescent chromatin. On the bottom, an alluvial plot of the proportion of each regulatory state at the blastula, gastrula, early larval and late larval stages. (**f**) Alluvial plot of the proportion of the Hox cluster of *O. fusiformis* exhibiting each chromatin state during embryogenesis and larval growth. (**g**) Representative tracks of H3K4me3, H3K4me1 and H3K27me3 marks in a Hox gene (*Hox4*) during the life cycle of *O. fusiformis*. (**h**) Dynamics of CUT&Tag scores in the H3K4me3, H3K4me1 and H3K27me3 peaks located in the Hox cluster during embryogenesis and larval development. On top, the coloured line shows the trend after regression. (**i**) Chromatin states in the Hox stripe anchor during *O. fusiformis*’ life cycle. (**j**) Gaudí plots projecting H3K4me3, H3K4me1 and H3K27me3 signals onto a two-dimensional Kamada-Kawai graph layout (top), with solid colours indicating statistically significant (*p*-value < 0.05) 2 Kb bins identified using a one-sided permutation test.

We identified 20,486, 28,471, 8,630, and 13,474 consensus peaks marked by H3K4me3, H3K4me1, H3K27me3, and H3K27ac, respectively, although the number of active peaks fluctuated during development (Figure 3b; Extended Data Figure 7b). As expected, the promoter mark H3K4me3 was highly enriched around transcription start sites (TSSs) (Extended Data Figure 7c; Supplementary Figure 18), and as observed through open chromatin profiling^2^, most peaks, regardless of the hPTM, were located within gene bodies, especially in introns, and one kilobase upstream of TSSs, rather than in intergenic regions (Extended Data Figure 7d, e; Supplementary Figure 19). However, promoter peaks were particularly prevalent for H3K4me3 (23.06%), whereas H3K4me1 (27.33%) and H3K27me3 (30.75%) showed larger proportions of intergenic peaks, consistent with their roles as enhancer marks^59^. In line with its function in other systems^60^, H3K4me3 was strongly correlated with gene transcription, as were H3K4me1 and H3K27ac, albeit to a lesser extent, whereas a transcriptionally repressive trend was evident for H3K27me3 (Figure 3c; Supplementary Figure 20). Despite this, the peak’s genomic location was also a strong, significant predictor of transcription (Extended Data Figure 7g, h). Therefore, *O. fusiformis* exhibits a canonical and dynamic hPTM landscape during embryogenesis and larval growth.

The genome-wide integration of these four hPTM marks with open chromatin and transcriptomic data^2^ allowed us to define 10 distinct chromatin states that, given their genomic locations, represented discrete regulatory elements and active conditions (Figure 3d; Extended Data Figure 8a–e). Specifically, we identified signatures of active promoters, active enhancers in intergenic and intronic regions, poised and poised-activated enhancers^61,62^, inactive enhancers, and transcriptional units. Throughout development, “quiescent” chromatin—areas lacking enrichment of any of the profiled marks and mainly found in intergenic and intronic regions—comprised between 62.2% and 67.7% of the genome, with the highest proportion observed at the early larval stage (Figure 3d, e; Extended Data Figure 8f). As expected, the proportion of actively transcribed chromatin increased with larval development and growth; however, the percentage of chromatin associated with active promoters decreased (Figure 3e). Interestingly, the proportion of poised (H3K4me1^+^, H3K27me3^+^, H3K27ac^−^) and poised activated (H3K4me1^+^, H3K27me3^+^, H3K27ac^+^) intergenic enhancers increased across the genome at the late larval stage, possibly reflecting a primed regulatory state that may enable rapid transcriptional responses to settlement cues during metamorphosis^63^ (Figure 3e). Likewise, active intronic enhancers, which represented the vast majority (68–80%) of the active enhancer landscape during *O. fusiformis* embryogenesis, were only 13% of the chromatin with an active enhancer state in the late larva (Figure 3e; Extended Data Figure 8g). This, together with the loosely defined chromatin conformation landscape at the early larval stage (Figure 1d–f), supports a genome-wide regulatory reorganisation from more local (intronic) to primarily distant (intergenic) regulation between the early and late larval stages in *O. fusiformis*.

### Polycomb-mediated repression of *Hox* genes is ancestral in bilaterians

Chromatin state dynamics in the Hox cluster differed from those observed genome-wide in *O. fusiformis* (Figure 3f; Extended Figure 8h). In the blastula and gastrula, the Hox cluster remained largely silenced, with quiescent chromatin accounting for 85.7% and 83.1% at those stages, respectively, compared to 67% and 65.5% genome-wide (Figure 3e, f). However, the Hox cluster underwent a notable regulatory reconfiguration in the early larva, marked by a sharp rise in the proportion of chromatin in a poised regulatory state (44.8%) (Figure 3f). This pattern again contrasts with genome-wide trends, in which poised and poised-activated enhancers peaked at the late larval stage (Figure 3e). As *Hox* genes become active in the late larvae, the fraction of actively transcribed chromatin, including promoters, as well as poised-activated, weak, and inactive enhancers, increased (Figure 3f; Extended Data Figure 8h). These chromatin state dynamics were mirrored at the level of individual hPTMs (Figure 3g; Extended Data Figure 9a, b). As in vertebrates^5^ and insects^64^, the Hox cluster became broadly marked by the Polycomb-mediated repressive mark H3K27me3 in the early larva of *O. fusiformis* (Extended Data Figure 9a, c), before the onset of *Hox* gene activation^2^. Concomitant with *Hox* transcription in the late larva in this annelid, there was a removal of H3K27me3 and an increase in H3K4me3 signal across the cluster (Extended Data Figure 9a, c), which was also reflected in the opposite dynamics of peak activity between repressive (H3K27me3) and permissive (H3K4me3 and H3K4me1) marks between the early and late larva stages (Figure 3h; Extended Data Figure 8i). Therefore, our findings support a cluster-wide repressive role of the Polycomb group in *Hox* gene regulation in the annelid *O. fusiformis*, indicating that shared regulatory principles govern *Hox* gene activity across bilaterians despite lineage-specific differences in the global regulatory logic controlling *Hox* genes.

To examine how hPTM dynamics relate to the formation of the architectural stripe and TAD along the Hox cluster in *O. fusiformis*, we first characterised the chromatin states in the distal regulatory element overlapping the stripe anchor (Figure 3i; Extended Data Figure 9d). During embryogenesis and in the early larva, this regulatory element exhibited small domains marked as inactive and weakly active enhancers (Figure 3i). However, coinciding with the activation of the *Hox* genes and the formation of the stripe and TAD, this distal regulatory element became a poised-activated enhancer, a class of enhancer typically associated with developmental genes^61,62^ (Figure 3i). This bivalency probably reflects the heterogeneous activation of *Hox* genes in the late larva, with the developing trunk primarily expressing these genes, rather than the rest of the larval tissues^2^. Moreover, the integration of chromatin conformation and hPTM data statistically confirmed the link between these two layers of genome regulation in the Hox cluster in *O. fusiformis* (Figure 3j; Extended Data Figure 8j, k). In the early larva, regions of the cluster enriched for H3K27me3 were in proximity, within a central domain depleted of H3K4me3 and H3K27ac (Figure 3j; Extended Data Figure 8k), suggesting that condensation of Polycomb complexes might be responsible for the chromatin architecture of the Hox cluster during gene repression, as in vertebrates and insects^57^. This association disappeared as *Hox* genes became active (Figure 3j), thereby suggesting that the activation of the global distal enhancer and consolidation of the architectural stripe reconfigure the regulatory state and conformation of the entire Hox cluster.

### A TAD encapsulating the Hox cluster is an ancestral trait

Temporal and spatial collinearity of *Hox* gene expression is observed across Spiralia, one of the three superclades of bilaterally symmetrical animals to which Annelida belongs^26,27^ (Figure 4a). To examine whether the formation of higher-order chromatin conformations correlates with the coordinated expression of *Hox* genes and cluster integrity in phylogenetically related groups, we studied representatives of five phyla (Chaetognatha, Mollusca, Phoronida, Brachiopoda, and Nemertea) with Hox clusters and available chromatin conformation data. The chaetognath *Paraspadella gotoi*^65^ and the brachiopod *Lingula anatina*^66^ have disorganised^13^ Hox clusters, with the brachiopod exhibiting two additional inversions. Neither group shows strict spatial collinearity in *Hox* gene expression^65,67^, and there was no obvious global chromatin conformation across the Hox cluster in these two species (Figure 4a). A similar situation occurred in the phoronid *Phoronis australis*, which has a disorganised Hox cluster^27,68^ and belongs to a clade without temporal or spatial collinearity^21^ (Figure 4a). However, the polyplacophoran *Lepidochitona cinerea*, a member of the only molluscan clade with reported Hox spatial collinearity^69^ and an organised Hox cluster, as well as the nemertean *Lineus longissimus*, which also has an organised Hox cluster and belongs to a group with spatial and temporal Hox expression collinearity^19^, showed signs of a TAD across their Hox clusters (Figure 4a). Notably, an architectural stripe downstream of *Hox1* delimited one of the TAD boundaries in the chiton *L. cinerea*, reminiscent of the condition observed in the annelid *O. fusiformis* (Figure 4a). Therefore, similar patterns of 3D chromatin conformation occur among spiralian clades with organised Hox clusters and at least spatial collinearity, suggesting that ancestral regulatory principles may govern *Hox* gene expression and help explain the conservation and erosion of *Hox* gene clusters in Spiralia.

**Figure 4.**
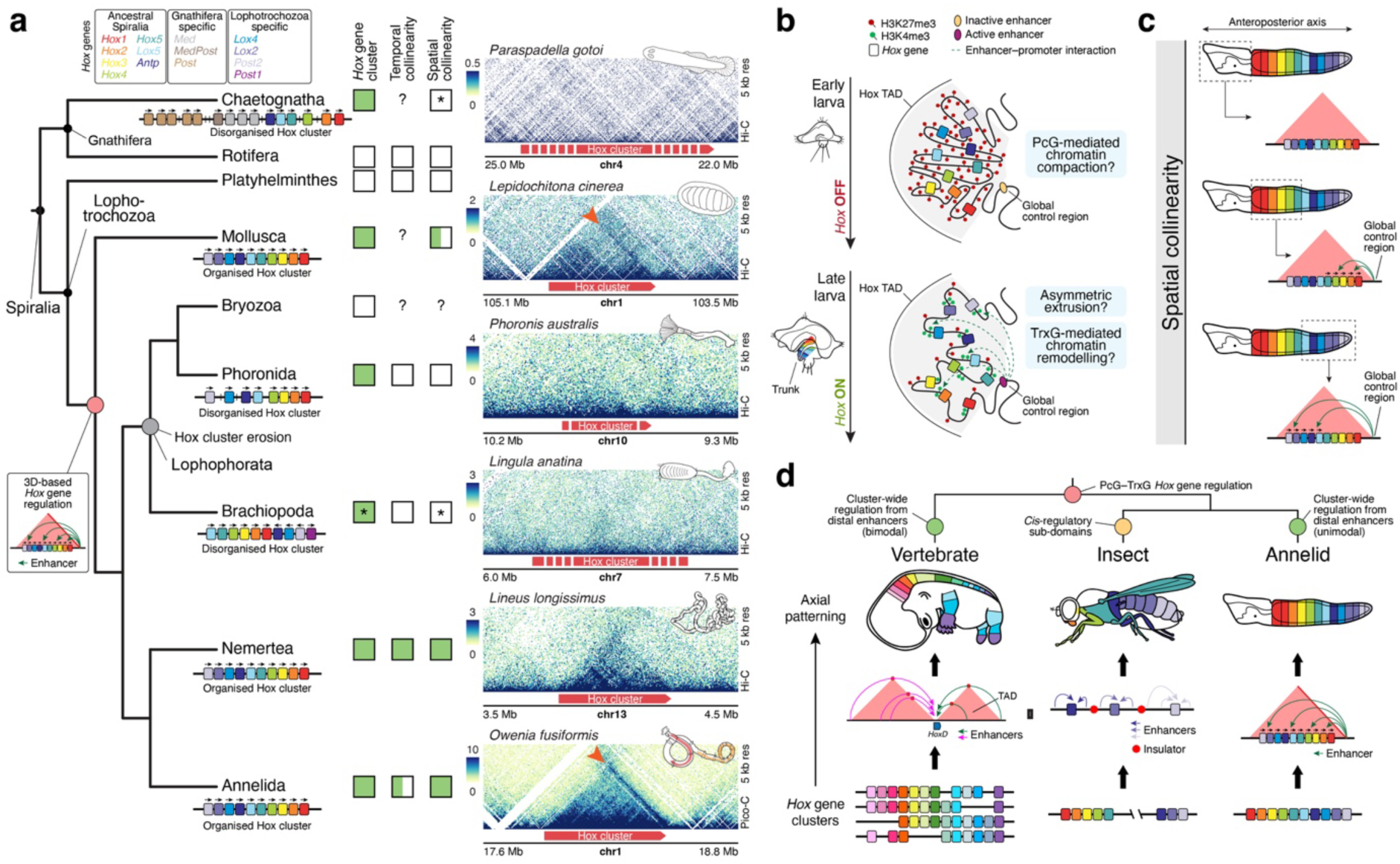
| The evolution of *Hox* gene regulation in Spiralia. (**a**) The chromatin architectural landscape of Hox clusters across Spiralia. On the left, a cladogram of the phylogenetic relationships of selected spiralian taxa, focusing on those with a Hox cluster (Chaetognatha, Mollusca, Phoronida, Brachiopoda, Nemertea, and Annelida). In the middle, presence (green squares) and absence (empty squares) of a Hox cluster and temporal/spatial collinearity across Spiralia. The asterisks in Chaetognatha and Brachiopoda indicate that collinearity involves only a subset of *Hox* genes and that the cluster is rearranged in the brachiopod *Lingula anatina*. Half-green, half-empty squares indicate that the trait is present in some species but absent in others. On the right, Pico-C contact heatmaps of the Hox clusters of the chaetognath *Paraspadella gotoi*, the polyplachophoran mollusc *Lepidochitona cinerea*, the phoronid *Phoronis australis*, the brachiopod *Lingula anatina*, the nemertean *Lineus longissimus*, and the annelid *O. fusiformis*. Spiralians with organised clusters and spatial collinearity exhibit a TAD in their Hox clusters. Moreover, the polyplachophoran *L. cinerea* shows signs of an architectural stripe (red arrowhead) in the same position as in *O. fusiformis*. Heatmaps represent normalised observed counts. (**b**, **c**) Schematic drawings of the proposed model of *Hox* gene regulation in annelids. Polycomb-mediated repression maintains *Hox* genes in an initially silenced state (**b**, top). The activation of *Hox* genes coincides with activation of the distal enhancer, formation of an architectural stripe, and removal of Polycomb-mediated repression (**b**, bottom). The formation of a stripe suggests a heterogeneous, context-dependent exposure of *Hox* genes to the distal enhancer (**c**), which could explain the spatial collinearity in Hox gene expression observed in annelids. (**d**) The proposed evolutionary scenario of *Hox* gene regulation in bilaterians. While the Polycomb-Trithorax interplay to repress and activate Hox genes, respectively, is probably an ancestral feature, major bilaterian groups have independently evolved lineage-specific mechanisms of coordinated *Hox* gene regulation. In particular, the use of distal enhancers acting as “global control regions” evolved convergently in vertebrates and Spiralia. Drawings are not to scale.

## Discussion

Our study demonstrates that a cluster-wide, long-range 3D chromatin conformation underpins the distinct dynamics of *Hox* gene regulation observed across annelids with different life cycles and larval types^2^. Annelid Hox clusters contain a large distal enhancer whose activity correlates with the onset of *Hox* gene expression, and, as suggested by the formation of an architectural stripe from this *cis*-regulatory element, it may act as a shared “global control region” for these genes (Figure 4b). Although the mechanisms generating architectural stripes are still debated and may be locus-specific^49–51,70,71^, these patterns in chromatin contact maps are often linked to asymmetric chromatin extrusion^49,50^, which would allow a common regulatory element, in this case the shared distal enhancer, to scan and regulate different loci within a contiguous chromatin domain, in this case the Hox cluster (Figure 4b). Notably, this distal enhancer is bound by genes involved in annelid trunk development and patterning (e.g., *lbx*^54^, *meis*, *soxB1*; Extended Data Figure 5f–m) and by known *Hox* gene activators, particularly components of the Polycomb-antagonist Trithorax SWI/SNF chromatin remodelling complex^57,72^. This is consistent with the broad deposition of Polycomb-mediated H3K27me3 repressive marks in the Hox cluster of *O. fusiformis* at the early larval stage, when *Hox* genes are not expressed. Therefore, our data suggest a testable model in which regulatory exposure of *Hox* genes to the distal enhancer would release global Polycomb-mediated cluster repression and compaction, thereby promoting *Hox* gene activation (Figure 4b). More importantly, this model would not only explain differences in the timing of *Hox* gene activation between species but also the temporal and spatial collinearity of these genes in annelids, as differences in when and which *Hox* genes are exposed to the distal enhancer could generate coordinated spatiotemporal patterns of gene expression (Figure 4c).

More broadly, our study clarifies the evolution of *Hox* gene regulation in bilaterian animals (Figure 4d). The repressive and activating interplay between the Polycomb and Trithorax protein groups^57^, respectively, is present in vertebrates, insects, and possibly annelids, and thus might represent an ancestral regulatory feature of *Hox* gene expression in Bilateria (Figure 4d). Moreover, the existence of a self-insulated domain across the organised Hox clusters of annelids, nemerteans, and molluscs, and more importantly, the formation of an architectural stripe 3’ of *Hox1* in the chiton *L. cinerea*, as in the annelid *O. fusiformis*, indicates that the principles of *Hox* gene regulation discovered in annelids may represent the ancestral condition for Lophotrochozoa^26^ (Figure 4a). This model of *Hox* gene regulation, which relies on a “global control region”, resembles the regulatory principles governing *Hox* genes in vertebrates^3,5–7^, thereby making the condition in insects a possible specialisation related to the evolution of anteroposterior body patterning based on gap and pair-rule genes^73^ (Figure 4d). However, certain insect lineages, such as lepidopterans (i.e., moths and butterflies), exhibit strong self-insulation around their Hox clusters^74^, although the developmental implications of this chromatin structure remain unknown, and intra-cluster boundaries remain essential for preserving *cis*-regulatory domains in these insects^11,75^.

The use of distal, global enhancers to regulate *Hox* gene expression in vertebrates and annelids is, however, most likely the result of convergent evolution (Figure 4d). Chromatin conformation data in cephalochordates (i.e., amphioxus), the sister group to all remaining chordates^76^, do not suggest the presence of distal global enhancers in the Hox cluster in this taxon^33,77^, indicating that the bimodal regulation of *Hox* genes is a vertebrate innovation^78^. Moreover, *Hox* gene repertoires, particularly median^65,79^ and posterior^65,77,80^ orthology groups, likely expanded independently in deuterostomes (e.g., chordates and echinoderms), ecdysozoans (e.g., insects and nematodes), gnathiferans (e.g., chaetognaths) and lophotrochozoans^81^. Therefore, the consolidation of specific *Hox* gene repertoires, clusters, and expression patterns, as in the last common lophotrochozoan ancestor (Figure 4a), may have co-occurred with the evolution of lineage-specific modes of *Hox* gene regulation. Notably, this scenario highlights the pervasiveness of developmental system drift^82,83^ in evolution, as comparable patterns of *Hox* gene expression along bilaterian anteroposterior axes, even between species of the same phylum (e.g., *O. fusiformis* and *C. teleta*), can emerge from substantially different regulatory principles.

Collectively, our work proposes a novel framework for studying genome regulation in bilaterians. The functional assessment of our proposed model of *Hox* gene regulation will not only reveal the mechanisms controlling these genes in annelids but also shed light on the ancestral roles of YY1 and other chromatin remodellers, as well as the molecular and biophysical principles underpinning higher-order chromatin structures in bilaterians. Likewise, extending high-resolution, genome-wide epigenomic analyses to a broader diversity of bilaterians will reveal the extent to which *Hox* gene regulation varies across lineages and clarify the evolutionary interplay among cluster-wide regulation, cluster preservation, and the spatiotemporal collinear expression of these genes. This, in turn, will transform our understanding of the origin and diversification of bilaterian body plans and life cycles.

## Methods

### Adult culture, spawning, and *in vitro* fertilisation

Sexually mature *O. fusiformis* Delle Chiaje, 1844 adults were obtained from intertidal waters near the Station Biologique de Roscoff (Centre National de la Recherche Scientifique (CNRS)-Sorbonne University, Roscoff, France) and maintained in-house as described before^84^. *Capitella teleta* Blake, Grassle & Eckelbarger, 2009, and *D. gyrociliatus* O. Schmidt, 1857 were cultured, and their embryos, larvae, and adults were collected following previously described protocols^25,85^.

### Pico-C

Pico-C libraries were generated using a previously published protocol^28^. Briefly, snap-frozen samples were mechanically dissociated using a pestle in phosphate buffer saline (PBS) with 0.5% Triton (PBST), or artificial seawater (ASW) in the case of the early larval samples of *C. teleta*, and crosslinked in a 2% formaldehyde solution at room temperature (RT) for 15 minutes in PBST/ASW. To quench the formaldehyde, 1.25 M glycine was added, and samples were then washed twice with PBST/ASW. Second crosslinking was performed with 3 mM of ethylene glycol bis(succinimidyl succinate) (ThermoFisher, 21565) in PBST/ASW at RT for 45 minutes. The reaction was quenched with 2 M Tris, pH 7.5, containing 0.5% Triton for 5 minutes, then all samples were washed twice with PBST and stored at −80 °C until further use.

To optimise the amount of micrococcal nuclease (MNase) required to generate chromatin monomers/dimers, crosslinked samples were resuspended in cold MB1 (50 mM NaCl, 10 mM Tris-HCl, 5 mM MgCl_2_, 1 mM CaCl_2_, 1× NP-40 (PIC, Roche, #5892791001) and crushed using a pestle until the solution was largely clear. Samples were then incubated on ice for 20 minutes and then centrifuged for 5 minutes at 5000 g and 4 °C. Samples were washed in MB1 and spun again and resuspended in 1× binding buffer (200 mM HEPES-KOH pH 8, 100 mM KCl, 10 mM CaCl_2_, 10 mM MnCl_2_, 5 mM spermidine, diluted in MB1 buffer) supplemented with concanavalin-A coated beads (1:100 volume: volume; BioMag BP531) washed and resuspended in 1× binding buffer. After incubation for 10 minutes at RT, the beads were washed and resuspended in MB1 before adding an appropriate amount of MNase. Samples were then incubated at 37 °C for 10 minutes at 950 rpm, quenched with 500 mM EGTA, washed in MB2 (50 mM NaCl, 10 mM Tris-HCl, pH 7.5, 10 mM MgCl2), and resuspended in extraction buffer (50 mM Tris-HCl, 50 mM NaCl, 1 mM EDTA, 1% SDS) with proteinase K. After an overnight incubation at 65 °C, DNA was purified following a standard phenol:chloroform:isoamyl alcohol extraction. The resultant extract was then quantified using a Qubit high-sensitivity assay and run on a 3% agarose gel. The amount of MNase that gave a strong mononucleosomal band (∼150 bp) with minimal undigested DNA was chosen for the full protocol.

For library preparation, following MNase incubation, EGTA quenching, and MB2 washes (as described above), samples were end-chewed with 25 U of T4 PNK (NEB #M0201) and incubated for 15 minutes at 37 °C at 700 rpm. Then, 25 U of Klenow (NEB, #M0210) were added, and samples were incubated for a further 15 minutes at 37 °C. After end labelling with Biotin-dATP (Jena biosciences #NU-835-BIO14) and Biotin-dCTP (Jena biosciences #NU-835-BIOX), ends were then ligated twice with 5000 U of T4 DNA ligase (NEB #M0202) for 2.5 hours at RT with slow rotation. Biotin-dNTPs were removed from unligated ends using Exonuclease III (NEB #M0206) for 15 minutes at 37 °C with interval mixing. Samples were then reverse crosslinked with proteinase K overnight at 65 °C. DNA was purified following standard phenol:chloroform:isoamyl alcohol extraction, then precipitated with 3 M sodium acetate (pH 5.2), cold 100% ethanol, and GlycoBlue (ThermoFisher AM9515) for at least 1 hour at -80 °C. DNA pellets were resuspended in TE buffer (10 mM Tris-HCl, pH 8.0, 1 mM EDTA), RNase-treated, and quantified using Qubit high sensitivity. After streptavidin binding, libraries were end-repaired with the NEBNext Ultra II End Repair module (NEB, E7546S), and Illumina adaptors were ligated with the NEBNext Ultra II Ligation module (NEB, E7595S), following manufacturer recommendations. To identify the sample-specific library amplification conditions, a fraction of the library amplification PCR reaction was collected after 10, 11, 12, and 13 cycles and run on a 2% TAE agarose gel, with the minimum number of PCR cycles required to show a visible band between 370 bp and 620 bp chosen as the optimal setup for final library preparation. Finally, AMPure XP beads (Beckman Coulter, A63881) were used for two-sided size selection. Libraries were sequenced using double-end 150 bp Illumina sequencing.

### Initial Pico-C data processing

Paired reads were aligned with BWA-MEM^86^ (v0.7.17) with parameters -5SP, -T0. Pairtools^87^ (v.0.3.0) was used to parse and identify valid ligation events using parameters --min-mapq 3 --walks-policy 5unique --max-inter-align-gap 30. Parsed pairs were sorted using pairtools sort, and duplicates were removed with pairtools dedup using parameters --max-mismatch 1 (Supplementary Figure 1; Supplementary Table 1).

### Genome scaffolding and reannotation

YaHS^88^ (v.1.2) was used to improve assembly contiguity of the genomes of *C. teleta* and *D. gyrociliatus* using the mapped Pico-C reads (Supplementary Figure 2). Juicer^89^ pre (v1.2) and Juicebox^90^ (v1.11.08) were used to generate and view a .hic file for manual correction of large inversions and misassemblies. Previous gene annotations^25,91^ were lifted over with Liftoff^92^ (v1.6.3), and repeat elements were predicted and annotated *de novo* with RepeatModeler2^93^ (v.2.0.5) and RepeatMasker (v.4.1.5) using RepBase as the reference. These assemblies, along with the previously published genome assembly and annotation for *O. fusiformis*^2^, were used in all downstream analyses.

### Compartment, TAD, and chromatin loop calling

Cooler^94^ cload pairix (v.0.8.6) was used to create 1 kb cooler files for all replicates, and cooler coarsen (v.0.8.6) was used to generate multiple resolutions, which were then merged into a single cooler file with cooler merge (v.0.8.6). Contact matrices were balanced with Cooler^94^ (v.0.8.6) using iterative correction and eigenvalue decomposition (ICE)^95^. HiC files were generated using the Juicer^89^ tools (v.1.22.01), and FAN-C cooler files were converted to FAN-C^96^ format and corrected using ICE.

For quality check, mapping stats were taken from the initial processing pipeline (see above). Principal component analysis was performed on cooler files at 10 kb resolution using FAN-C^96^ PCA, and hicCorrelate^97^ (v.3.7.3) was used to produce correlation matrices from 10 kb cooler files. For resolution plots, replicate cooler files were merged, and resolutions of 1, 2, 3, 5, 10, and 50 kb were generated to calculate the percentage of bins with more than 1,000 contacts. Contact frequency plots were generated using Cool Tools^94^ (v.0.8.6) at a resolution of 20 kb.

A/B compartments were called using the FAN-C pipeline^96^, with FAN-C hic (--low-coverage-auto -n -m ICE --deepcopy) and resolutions of 100 kb for *O. fusiformis* and 50 kb for *C. teleta* and *D. gyrociliatus*. Observed over expected (O/E) contact maps were generated, and eigenvectors 1 to 3 were plotted for each chromosome and sample and individually examined to identify the eigenvector that correlated with the pattern seen on the contact map. The chosen eigenvectors were then oriented using RNA-seq data previously generated in the lab for each time point^2,25^. Those with higher RNA-seq expression were deemed A compartments, and those with lower RNA-seq expression were deemed B compartments. Saddle plots were generated using FAN-C compartments^96^, classifying bins into 10 percentiles based on eigenvector values. The average O/E values for contact pairs were calculated and plotted in the saddle plot.

TADs were called with Arrowhead^98^ from juicer tools^89^ (v.1.22.01) using .hic files with the parameter -m 2000 at resolutions of 4 kb for *O. fusiformis* and *D. gyrociliatus,* and 2 kb for *C. teleta* (except for the *Hox* gene cluster, for which 5 kb resolution was used). TADs that correlated to minor misassemblies when visualised in Juicebox were manually removed. Chromatin loops were called with SIP^99^ (v.1.6.4) in ICE normalised 2 kb cooler contact matrices for all samples with parameters (-t 2000 -fdr 0.1 -d 6 -mat 1000 -nbZero 6 -cpu 4 -g 1.5 -min 2 -max 2). Species-specific consensus TADs and chromatin loops were generated with BEDTools^100^ (v2.31.1) merge and AQuA tools^101^, respectively. TAD and chromatin loop aggregate plots were generated with the FAN-C pipeline^96^. Contact interaction heatmaps were created with plotardener^102^ (v.1.15.1).

### Compartment, TAD, and chromatin loop characterisation

BEDTools^100^ (v2.31.1) coverage was used to generate TE, gene, and open chromatin coverage databases (based on previously published ATAC-seq datasets^2^) for the A/B compartments (Supplementary Table 2). Gene body cytosine methylation levels were calculated as previously described^103^. BEDTools^100^ (v2.31.1) nucBed was used to calculate GC content per compartment, and BEDTools^100^ (v2.31.1) intersect with a -F 0.5 was used to classify genes by compartment so that any genes falling between compartments were assigned to the compartment they lay mostly in. Compartment coefficients were estimated as previously described^104^. A similar approach was used to assign genes and repeats to TADs and loop anchors, and estimate repeat, gene, and open chromatin coverages for these structures (Supplementary Tables 3–9).

To explore the changes in A/B compartments during the life cycle of *O. fusiformis* and *C. teleta*, oriented eigenvectors for developmental stages were merged in an R dataset, and any area where the eigenvector sign did not match was classed as a ‘flip’ (Supplementary Tables 10 and 11). Previously published RNA-seq datasets for *O. fusiformis* and *C. teleta*^2^ were used to assess the log2-fold change in expression of differentially expressed genes (*P* value < 0.05) that switched compartment, as calculated with DESeq2^105^. BEDTools^100^ (v2.31.1) intersect with a reciprocal overlap of 80% (-f 0.8 -F 0.8) for TADs and 90% (-f 0.9 -F 0.9) for loops was used to identify dynamic TADs and chromatin loops across stages. To identify enriched Gene Ontology terms, the annotated GO terms for *O. fusiformis* and *C. teleta* (Supplementary Tables 14 and 15) were expanded to include all ancestral biological process terms using GOBPANCESTOR from GO.db R package (v.3.23.1). These terms were then mapped to the generic GO Slim ontology using the GSEABase^106^ (v.1.72.0) R package by intersecting ancestral path with Slim IDs. To maintain one-to-one gene-category output, terms mapping to the same Slim category were collapsed. GO enrichment was performed on gene groups of interest using the topGO^107^ R package (v.2.62.0) against the GO Slim gene universe assigned above. A classic Fisher’s exact test was used to identify gene categories significantly enriched within these groups. Results were then filtered to only retain terms defined within the GO Slim collections.

Transcription factor and developmental gene content in consensus TADs, TAD borders and loop anchors in *O. fusiformis*, *C. teleta* and *D. gyrociliatus* was estimated using previously annotated transcription factor genes^2^ and a list of Gene Ontology terms related to development and transcription regulation (Supplementary Tables 12–16). One-to-one orthologs between *O. fusiformis* and *C. teleta* and gene ages (phylostrata) were identified based on previously computed gene families^91^ (Supplementary Tables 17–19).

Consensus TAD boundaries and chromatin loop anchors were annotated to specific genomic features using HOMER2^108^ (v.4.11) and the annotatePeaks.pl script (Supplementary Tables 20–23).

### Sequence conservation analyses

Progressive Cactus^109^ (v.2.1.1) was used to build a whole-genome alignment of 12 annelids and two mollusc outgroups (Supplementary Table 24). All genomes were soft-masked with RepeatMasker v4.1.2, using a *de novo* repeat library built for each genome with RepeatModeler2^93^ (v.2.0.2). To build a guide species tree for Cactus, we ran a homology-based annotation with MetaEuk^110^ (v.6) using the AnnotateSnakemake pipeline^111^ with parameter --config metaeuk_only=’y’ and the proteomes of *O. fusiformis, C. teleta,* and *D. gyrociliatus* as reference for the five genomes (*Magelona johnstoni, Hermodice carunculata, Alitta virens, Sabella spallanzanii* and *Arenicola marina*) without an available gene annotation. 962 one-to-one ortholog groups were identified with OrthoFinder^112^. Multiple protein sequence alignments for each orthogroup were produced with MAFFT^113^ (v.7.508) and used to build a tree from the concatenated alignment with RAxML-NG^114^ (v.1.1.0), using the LG+G4+F model, 10 starting parsimony trees, and 100 bootstrap replicates. After alignment with Progressive Cactus^109^ (v.2.1.1), coverage statistics were obtained with the halCoverage tool, and sequence conservation scores for *O. fusiformis*, *C. teleta* and *D. gyrociliatus* were computed with phastCons from the PHAST^115^ package (v.1.4). To achieve this, the Cactus alignment was converted to .maf format using the halToMaf utility from Cactus, with parameters --noAncestors and --onlyOrthologs. We next filtered the resulting .maf alignments to remove columns consisting solely of gaps and estimated a neutral model from fourfold degenerate sites (4d) using PhyloFit (from PHAST^115^ v.1.4). We first used phastCons to estimate parameters for a conserved and a non-conserved model from the .maf alignment and the previously estimated neutral model, and then used phastCons a second time for prediction, that is, to compute conservation scores (Supplementary Table 25).

### Transcription factor motif transfer

To annotate the transcription factor complements of *O. fusiformis*, *C. teleta*, and *D. gyrociliatus*, we used hmmsearch^116^ using a set of 89 HMM models from the Pfam-A database that correspond to DNA-binding domains specific to different transcription factor families. The domain hits were then extracted from each protein, along with up to 30 additional flanking amino acids on each side to retain the sequence context around the DNA-binding domain. Each transcription factor family was subsequently clustered using MCL^117^ with an inflation parameter of 1.1 to define putative homology groups within each family. Each homology group was then aligned using MAFFT^113^ in L-INS-i mode, and the resulting alignments were trimmed using ClipKIT^118^ and used as input for phylogenetic inference with IQ-TREE2^119^. The resulting trees were parsed to obtain orthology groups using *Possvm*^120^ with *Mus musculus* as the reference species for orthogroup naming.

Binding motifs represented as position weight matrices (PWMs) for the annotated transcription factors were predicted using two complementary strategies. First, using the JASPAR inference tool with default parameters, which is based on similarity regression^121^. Second, 4,558 experimentally determined PWMs from the CIS-BP database^122^ were transferred from the phylogenetically closest reference sequences within each orthogroup, based on the previously inferred gene trees (see above). For each query protein, a single motif was selected from sister sequences. Candidate PWMs were ranked by their information content score and average similarity to all other motifs, ensuring that the selected motif was both high-quality and representative. This approach resulted in 385, 566, and 389 transferred motifs for *O. fusiformis*, *C. teleta*, and *D. gyrociliatus*, respectively. Final motif predictions for each species were obtained by combining both approaches. When both gene tree-based and JASPAR-based predictions were available for a protein, the gene tree-based prediction was preferred. When only a JASPAR-based prediction was available, that prediction was retained (Supplementary Tables 26–28).

### Motif enrichment, annotation, and footprinting

To identify significantly enriched DNA binding motifs in TAD boundaries and chromatin loop anchors, we first used BEDTools^100^ (v2.31.1) to overlap these regions (+\- 1 bin) with consensus chromatin accessible regions as defined by previously published ATAC-seq datasets^2^. For TAD boundaries, this resulted in 4,002 ATAC-seq peaks in *O. fusiformis* and 3,446 peaks in *C. teleta*. For chromatin loop anchors, this yielded 1,450 ATAC-seq peaks in *O. fusiformis* and 1,675 peaks in *C. teleta*. Enriched DNA-binding motifs were then identified using HOMER^108^ (v.5.1) FindMotifsGenome.pl on these extracted ATAC-seq peaks in *de novo* motif discovery mode with equivalent-sized and GC-content genomic regions used as background sequences. HOMER^108^ (v.5.1) annotatePeaks.pl was then used to map the most enriched motifs to border elements and to calculate motif enrichment around TAD boundaries (Supplementary Tables 29–34). HOMER^108^ (v.5.1) annotatePeaks.pl was also used to annotate the transferred transcription factor DNA-binding motifs (see above) in the Hox cluster distal enhancers (stripe anchors) and internal cluster boundaries in *O. fusiformis* and *C. teleta* (Supplementary Tables 35–37).

To predict transcription factor binding, we conducted footprinting analysis using TOBIAS^123^ (v.0.13.3). First, replicate-specific mapped ATAC-seq .bam files were merged and indexed using SAMtools^124^ (v.1.19.2). TOBIAS ATACorrect was then used to account for Tn5 transposase bias in the consensus ATAC-seq peak set. Corrected signals were then utilised to calculate footprinting scores across the consensus peak set, thereby identifying regions of localised depletion. Finally, differential binding scores were identified using TOBIAS BINDetect by integrating footprinting scores with known motif positions. Transferred transcription factor DNA-binding motifs (see above) were scanned genome-wide across ATAC-seq peaks, and binding occupancy was compared across developmental time points to identify differential binding scores (Supplementary Tables 38 and 39). At the predicted Hox cluster stripe anchor, footprinting scores were assessed on common motifs found in both *O. fusiformis* and *C. teleta* (Supplementary Table 40). For any motifs with multiple hits in the stripe anchor, we took the mean footprinting scores at each time point and used Pearson’s correlation coefficient to assess similarities in transcription factor binding scores across these motifs, species, and time points.

### Insulator and YY1 protein evolution

The presence and absence of metazoan insulator and chromatin regulator proteins were inferred from previously precomputed gene families^91^ and through manual sequence similarity searches on the predicted proteomes of *O. fusiformis*, *C. teleta* and *D. gyrociliatus*. To resolve the orthology of metazoan YY1 proteins, we performed multiprotein sequence alignment with MAFFT^113^ (v.7) (Supplementary Figure 11), removed poorly aligned regions with trimAl^125^ and conducted phylogenetic reconstruction with IQ-TREE2^119^. AlphaFold 3^126^ was used to reconstruct the protein structure of the annelid and human YY1 orthologs.

### Microsynteny analyses

Blocks of collinear microsynteny between *O. fusiformis* and *C. teleta* were inferred with MCScanX (Supplementary Table 41) with parameters -b 2 and -s 2 to report only inter-species conserved collinearity with at least two genes. This identified 11 blocks of conserved microsyntenic collinearity, including the Hox cluster. BEDTools^100^ (v2.31.1) intersect with consensus TADs and chromatin loops indicated how many of these blocks overlap with high-order chromatin conformations.

### pA-Tn5 protein production and transposome assembly

The pTXB1-3xFLAG-pA-Tn5-Fl (Addgene, cat#: 124601) was a gift from Steve Henikoff (Fred Hutchinson Cancer Center, Seattle, WA, USA). pA-Tn5 expression and purification were performed as previously described with minor modifications^127–129^. Homemade BL21 *Escherichia coli* cells were transformed with the expression plasmid. Lysogeny broth medium with 100 mg·l^−1^ ampicillin was used to grow a culture at 37 °C and 250 rpm agitation to an optical density at 600 nm of 0.5 to 0.7. Protein expression was induced with 0.25 mM isopropyl-β-D-1-thiogalactopyranoside (IPTG) overnight at 18 °C and 250 rpm of agitation. Cells were harvested at 9,500 × *g* for 1 h, and pellets were resuspended in ice-cold purification buffer, i.e., 1× HEGX buffer (20 mM 4-(2-hydroxyethyl)-1-piperazineethanesulfonic acid (HEPES)-KOH pH 7.2, 1 M NaCl, 1 mM ethylenediaminetetraacetic acid (EDTA), 10 % glycerol, 0.2 % Triton X-100, supplemented with 1× cOmplete™ EDTA-free Protease Inhibitor Cocktail Tablets). Lysis was achieved by a combination of chemical lysis (Triton X-100 in buffer), enzymatic lysis (lysozyme, 100 mg·l^−1^), and physical lysis using ultrasonic probe-tip sonication (VCX 130 Vibra-Cell™ Ultrasonic Liquid Processor, Sonics). Crude lysates were centrifuged at 16,000 × *g* for 30 min at 4 °C and frozen at –80 °C until purification. pA-Tn5 purification was performed using chitin affinity chromatography (NEB). Protein was bound and washed twice in purification buffer and eluted through a 48-h incubation at 4 °C of the protein-bound resin with purification buffer supplemented with 100 mM dithiothreitol (DTT). The eluate was dialysed in SnakeSkin™ Dialysis Tubing with a 7 kDa molecular weight cut-off (MWCO) (Thermo-Fisher) against dialysis buffer (100 mM HEPES-KOH pH 7.2, 200 mM NaCl, 0.2 mM EDTA, 1.7 mM DTT, 0.2 % Triton X-100, 20 % glycerol). Protein was concentrated using 5–10 ml Vivaspin™ concentrators with a 10 kDa MWCO (Sartorius) and glycerol was added to a final concentration of 50% for long-term storage at –20 °C. Protein expression, size, purity, and concentration were assessed via SDS-PAGE.

To assemble the active transposome, Tn5ME-A, Tn5ME-B and Tn5MErev single-stranded oligonucleotides (Supplementary Table 42) were resuspended to 200 µM in annealing buffer (10 mM tris(hydroxymethyl)aminomethane (Tris)-HCl, pH 8.0, 50 mM NaCl, 1 mM EDTA). As described before^127–129^, Tn5ME-A/rev and Tn5ME-B/rev mosaic end double-stranded (MEDS) oligonucleotides were pre-annealed by mixing equal volumes of the single-stranded oligonucleotides, incubated at 95 °C for 2 min and let cool at RT. Pre-annealing was verified through native PAGE. pA-Tn5 transposome assembly was performed by incubating MEDS oligonucleotides with 12.5 volumes of pA-Tn5 fusion protein at RT for 50 min before long-term storage at –20 °C. Transposome activity was assessed *in vitro* using phage λ DNA (7.5 µl of 20 ng·µl^-1^ stock) as previously described^127–129^.

### Cleavage Under Targets and Tagmentation (CUT&Tag)

CUT&Tag was performed as previously described^127,130,131^, with minor adaptations for our biological material. *Owenia fusiformis* embryos and larvae were collected and reared until the desired developmental stage at 19 °C in ASW supplemented with penicillin (100 units·ml^-1^) and streptomycin (100 mg·l^-1^). To obtain lysates of 50,000 cells, ∼850 blastulae (5 hpf), 500 gastrulae (9 hpf), 200 mitraria larvae (24 hpf), and 60 competent larvae (3 weeks post-fertilisation (wpf)) were combined. Two biological replicates for H3K4me3, H3K27me3, H3K4me1, and H3K27ac, and a control dataset for non-specific antibody profiling (IgG-control) were carried out for each developmental time point (3 biological replicates for H3K4me3, H3K27me3, and H3K4me1 for the competent larva stage), with each sample coming from the same *in vitro* fertilisation batch. Embryos/larvae were incubated for 10 min in 0.5 ml CMF-ASW, spun down for 3 min at 5,000 × *g*, resuspended and washed in 0.5 ml 1× PBS and harvested again. Nuclei were isolated by resuspending and incubating for 10 min in 0.5 ml NE1 buffer, mechanically disrupting the embryos/larvae with a pellet pestle motor for approx. 1 min in pulses of 2 s on–1 s off, and incubating the samples in NE1 buffer for 10 min again. Nuclei were harvested for 5 min at 5,000 × *g*, washed once in 0.5 ml 1× PBS, harvested again, and resuspended in 50 µl wash buffer (20 mM HEPES-KOH pH 7.5, 150 mM NaCl, 0.5 mM spermidine) per 50,000 nuclei, i.e., 250 µl per replicate per developmental stage. 27.78 µl DMSO were added to the 250 µl sample (for a 9:1 wash buffer:DMSO ratio) for nuclei cryopreservation at –80 C in a Mr Frosty™ Freezing Container (Thermo Fisher Scientific) filled with isopropanol, until ready for library preparation. Samples were gently thawed in a water bath at RT. Concanavalin A-coated magnetic beads (Bangs Laboratories) were washed twice in binding buffer (20 mM HEPES-KOH, pH 7.9, 10 mM KCl, 1 mM CaCl_2_, 1 mM MnCl_2_), allowing for the beads to clear after each wash by placing them on a magnet stand, and finally resuspended in the initial volume of beads of binding buffer. We added 10 µl of activated beads per 50,000 nuclei and incubated them at RT for 5 min. The unbound supernatant was removed, and bead-bound nuclei were resuspended in 50 µl dig-wash buffer (0.05 % digitonin in wash buffer) supplemented with 2 mM EDTA, 0.1 % BSA and a 1:50 dilution of the primary antibody, or no antibody for the non-specific antibody profiling (IgG-control). The primary antibody was incubated overnight at 4 °C with rocking. Samples were placed on a magnet stand to clear, and the solution was withdrawn. Nuclei were incubated with 100 µl of a 1:100 dilution of the secondary antibody in dig-wash buffer for 30 min at RT, then washed 3× in 1 ml dig-wash buffer using the magnet stand to remove unbound antibodies. The pA-Tn5 transposome was let bind to the antibodies by incubating the nuclei at RT for 1 h with 100 µl of a 1:100 dilution of pA-Tn5 transposome (∼50 nM) in dig-300 buffer (20 mM HEPES-KOH pH 7.5, 300 mM NaCl, 0.5 mM spermidine, 0.01 % digitonin, supplemented with 1× cOmplete™ EDTA-free Protease Inhibitor Cocktail Tablets). Nuclei were washed 3× in 1 ml dig-300 buffer, then 300 µl of dig-300 buffer with 10 mM MgCl_2_ was added for tagmentation for 1 h at 37 °C. Tagmentation was stopped, and DNA fragments were solubilised by adding 10 µl of 0.5 M EDTA, 3 µl of 10 % SDS and 2.5 µl of proteinase K 20 mg·ml^-1^ to the 300 µl samples, which were incubated at 55 °C for 1 h. DNA fragments were purified through a standard phenol:chloroform:isoamyl alcohol precipitation and resuspended in a 1/10 dilution of Tris-EDTA (TE) buffer (10 mM Tris-HCl, pH 8.0, 1 mM EDTA). Libraries were amplified for 14 PCR cycles using a different i7 barcode for each sample (Supplementary Table 43) and the NEBNext HiFi 2× PCR Master mix (not hot-start) (NEB). After PCR clean-up with SPRIselect beads (Beckman Coulter), libraries were analysed using a High Sensitivity D1000 ScreenTape on an Agilent 2200 TapeStation to estimate fragment size distribution before pooling at equal molecular weight. Primary antibodies used were as follows: 1:50 rabbit anti-trimethyl-histone H3 (Lys4) antibody (H3K4me3 polyclonal antibody) (Millipore-Sigma Aldrich, cat#: 07-473, RRID: AB_1977252), 1:50 rabbit anti-trimethyl-histone H3 (Lys27) antibody (H3K27me3 polyclonal antibody) (Millipore-Sigma Aldrich, cat#: 07-449, RRID: AB_310624), 1:50 rabbit recombinant anti-histone H3 (mono methyl K4) antibody [ERP16597] – ChIP grade (H3K4me1, monoclonal antibody) (Abcam, cat#: 176877, RRID: AB_2637011), 1:50 rabbit anti-acetyl-histone H3 (Lys27) antibody (H3K27ac, polyclonal antiserum) (Millipore-Sigma Aldrich, cat#: 07-360, RRID: AB_310550). Secondary antibody used was guinea pig anti-rabbit IgG (heavy & light chain) antibody – preadsorbed (IgG-control, polyclonal antibody) (Antibodies-Online, cat#: ABIN101961, RRID: AB_10775589). Sequencing was performed over 4 different lanes of a NextSeq2000 platform in 2 × 50 bases mode at the Barts and The London Genome Centre (Queen Mary University of London, UK) (Supplementary Table 44).

### CUT&Tag data analysis

The core CUT&Tag analysis pipeline was similar to previous published work^127,130,132^. Trimmomatic^133^ (v.0.39) was used to remove sequencing adaptors. Surviving reads were mapped against both the chromosome-level assembly of *O. fusiformis*^2^ (GenBank: GCA_903813345.2) and the *E. coli* strain K-12 substrain MG1655 genome assembly (GenBank: GCA_000005845.2) using bowtie2^134^ (v.2.4.4) in paired-end mode (*O. fusiformis*: --local --very-sensitive --no-mixed --no-discordant -I 10 -X 700; *E. coli*: --local --very-sensitive --no-overlap --no-dovetail --no-mixed --no-discordant -I 10 -X 700). Apparent duplication rate was estimated using Picard (v.2.25.6) (http://broadinstitute.github.io/picard/) (Supplementary Tables 44 and 45). We used SAMtools^124^ (v.1.12) to convert .sam to .bam files, from which we estimated the fragment size distribution. We obtained the associated .bed files and retained only read pairs on the same chromosome/scaffold and with a fragment length of less than 1 kb using the bamtobed command from BEDTools^100^ (v.2.30.0). We used the genomecov command from BEDTools to generate normalised .bedgraph files using the number of fragments mapped to the *E. coli* genome as the spike-in/normalising factor for each sample and the bedGraphToBigWig^135^ (v.385) utility to create the final coverage files in .bw format. Peak calling was done using MACS2^136^ (v.2.2.7.1) in both narrow (-f BAMPE -g 0.44e9 --keep-dup all --nolambda --nomodel --min-length 100 --max-gap 75) and broad peak modes (including the –broad argument as well) using the IgG experiment as the negative control (-c), to unbiasedly assess which peak type each hPTM displays in previously uncharacterised annelids (Supplementary Table 46). For each combination of stage and hPTM, reproducible peaks between biological replicates were obtained using the Irreproducible Discovery Rate (IDR)^137^ (v.2.0.4.2) (*P* values < 0.05) (Supplementary Table 47). Afterwards, we used DiffBind^138^ (v.3.4.11) to generate the consensus peak sets. FRiP score-based benchmarking proved that the narrow peak-calling mode was optimal for all hPTMs (Supplementary Tables 48–50). We produced a consensus peak set for each hPTM, from which Tn5-biased peaks shared across all hPTM peak sets and the consensus ATAC-seq peak set were removed (Supplementary Table 51). The final filtered consensus peak sets contained 20,486 (H3K4me3), 8,630 (H3K27me3), 28,471 (H3K4me1), and 13,474 (H3K27ac) peaks, over which the CUT&Tag score was normalised with the DESeq2 method (Supplementary Tables 52–55). Both the IDR peak sets and the DiffBind peak sets were classified by genomic feature, and their nearest transcript and/or promoter region were annotated with HOMER^108^ (v.4.11) (Supplementary Tables 56–59). Peak size comparisons were done using a one-way ANOVA, followed by two-tailed post-hoc Tukey tests for pair-wise comparisons. All *P* values derived from pair-wise comparisons were adjusted using the stringent Bonferroni method for multiple testing correction (Supplementary Table 60). UpSet plots were used to identify stage-specific and constitutive peaks and genomic regions. The Pearson correlation coefficient between CUT&Tag peak dynamics and gene expression was computed individually for each CUT&Tag peak and its nearest annotated transcript (or the nearest transcript associated with the nearest promoter for peaks in intergenic regions) using two-sided Pearson tests. The predictive effect of the genomic feature and the hPTM on this correlation was analysed using linear models of each predictor alone (i.e., lm(correlation∼hPTM) and lm(correlation∼feature)) and as a bivariate model with a modelled interaction (i.e., lm(correlation∼hPTM*feature) (Supplementary Tables 61 and 62). Lastly, we used the intersect command in BEDTools^100^ (v.2.30.0) to subset the consensus peaks within the *Hox* cluster and analyse the signal dynamics at the peaks of interest (Supplementary Table 63). pyGenomeTracks^100^ (v.2.1) and deepTools^139^ (v.3.4.3) were used to visualise peak tracks and gene structures.

### Chromatin states annotation

ChromHMM^140^ (v.1.24) was used to identify and annotate chromatin states in *O. fusiformis*. For that, in addition to the CUT&Tag data generated in this study (see above), we used previously published RNA-seq^2^, ATAC-seq^2^, and whole-genome bisulfite sequencing^103^. Genomic features such as TSSs, exons, and intergenic regions were inferred from the genome annotation, and CpG islands were identified using cpgplot in the EMBOSS package^141^. After signal binarisation, a model was trained with ChromHMM’s LearnModel function with default parameters on both a stacked and a concatenated approach. After testing varying state numbers, a 12-state model was selected, and states were annotated based on their signal combinations and genomic feature enrichment (Supplementary Tables 64 and 65).

### METAloci

METAloci^142^ (v.1.3.6) was used to identify the spatial distribution and autocorrelation of CUT&Tag signal (H3K4me3, H3K4me1, H3K27me3, and H3K27ac) at the early and late larval stages along the Hox cluster of *O. fusiformis* at a contact map resolution of 2 kb with default parameters. Signal tracks were binned to 2 kb windows, combining replicates by calculating the statistical mean for each window. Spatial and epigenetic signals were visualised as Gaudí plots, applying the local Moran’s index for each bin of the plot.

### Chromatin architecture analyses in spiralian *Hox* gene clusters

Available Hi-C data (Supplementary Table 66) for five spiralian species (the chaetognath *Paraspadella gotoi*, the mollusc *Lepidochitona cinerea*, the phoronid *Phoronis australis*, the brachiopod *Lingula anatina*, and the nemertean *Lineus longissimus*) with chromosome-level genome assemblies^27,65,66,143^ were processed as described above. For *Lepidochitona cinerea* and *Lineus longissimus*, the Hox cluster was identified through sequence similarity searches on the genomic assembly using *Hox* genes from closely related species. Hi-C contact maps were plotted with plotardener^102^ (v.1.15.1).

### Gene expression analyses

Fragments of *meis*, *runx*, *sox2*, *pax6*, *Notch*, *prospero*, *foxA*, *musashi*, and *Hox* genes for *O. fusiformis* and *C. teleta* (Supplementary Table 67) were isolated as previously described^144^ using gene-specific oligonucleotides and a T7 adaptor. Riboprobes were synthesised using a T7 MEGAscript kit (ThermoFisher, AM1334) and stored at a concentration of 50 ng/µl in hybridisation buffer at –20 °C. Whole-mount *in situ* hybridisation in larvae of *O. fusiformis* and *C. teleta* was performed as described elsewhere^85,144^, but with the change of permeabilising with a detergent solution (150 mM NaCl, 160 mM Tris-HCl, pH 7.5, 1 mM EDTA, pH 8, 1 % SDS, 0.5 % Tween-20) for 30 min at RT. Differential interference contrast (DIC) images of the colourimetric *in situ* were obtained using a Leica 560 DMRA2 upright microscope equipped with an Infinity5 camera (Lumenera). DIC images were digitally stacked with Helicon Focus 7 (HeliconSoft). Brightness and contrast were edited with Adobe Photoshop CC (Adobe Inc.).

## Data availability

All sequencing data generated in this study are available at the European Nucleotide Archive (Pico-C libraries) and the Gene Expression Omnibus repository (CUT&Tag datasets).

## Code availability

All code generated in this study is available in Zenodo.

## Acknowledgements

We thank all members of the Martín-Durán lab for their help and support, as well as the technical staff at the Department of Biology at Queen Mary University of London. This work utilised Queen Mary’s Apocrita HPC facility, supported by QMUL Research-IT. This work was funded by grants to JMM-D from the European Union Horizon 2020 Framework Programme under the European Research Council (ERC Starting Grant agreement number 801669), the Biotechnology and Biological Sciences Research Council (BB/T008709/1 and BB/Y004221/1) and the Leverhulme Trust (RPG-2023-121). Research in AS-P’s group was supported by the European Research Council (ERC-COG 101170846) and the Spanish Ministry of Science and Innovation (PID2024-158058NB-I00). GZ was supported by the INPhINIT PhD fellowship from LaCaixa Foundation LCF/BQ/DI21/11860036. Y-JL was supported by Academia Sinica (AS-CDA-112-L06 and AS-GCS-114-L08). PJH and NRZ were supported by the Biotechnology and Biological Sciences Research Council (BBSRC Grant agreement numbers BB/V009311/1 and UKRI698, respectively). FM and EP are supported by a Royal Society University Research Fellowship (URF\R1\191161 and URF\R\241014), a Biotechnology and Biological Sciences Research Council research grant (BB/V01109X/1), a Leverhulme Trust research grant (RPG-2025-274) and a Newton International Fellowship from the Royal Society (NIF\R1\222125). Work in the Vaquerizas Laboratory is supported by the Medical Research Council, UK (award no. MC_UP_1605/10 to JMV), the Academy of Medical Sciences and the Department of Business, Energy and Industrial Strategy (award no. APR3\1017 to JMV).

## Author contributions

BED, FMM-Z, and JMM-D conceived and designed the study. BED, NE, and NM performed chromatin conformation experiments. FMM-Z performed histone post-translational modification profiling. TF did gene expression analyses. GZ and AS-P performed DNA-binding motif identification. BED, FMM-Z, EP, KG, and JMM-D performed computational analyses. Y-JL, FM, JMV, PJH, and NRZ contributed to the analysis of chromatin conformation data. BED, FMM-Z and JMM-D drafted the manuscript. All authors read and critically commented on the manuscript.

## Extended Data Figures

**Extended Data Figure 1.**
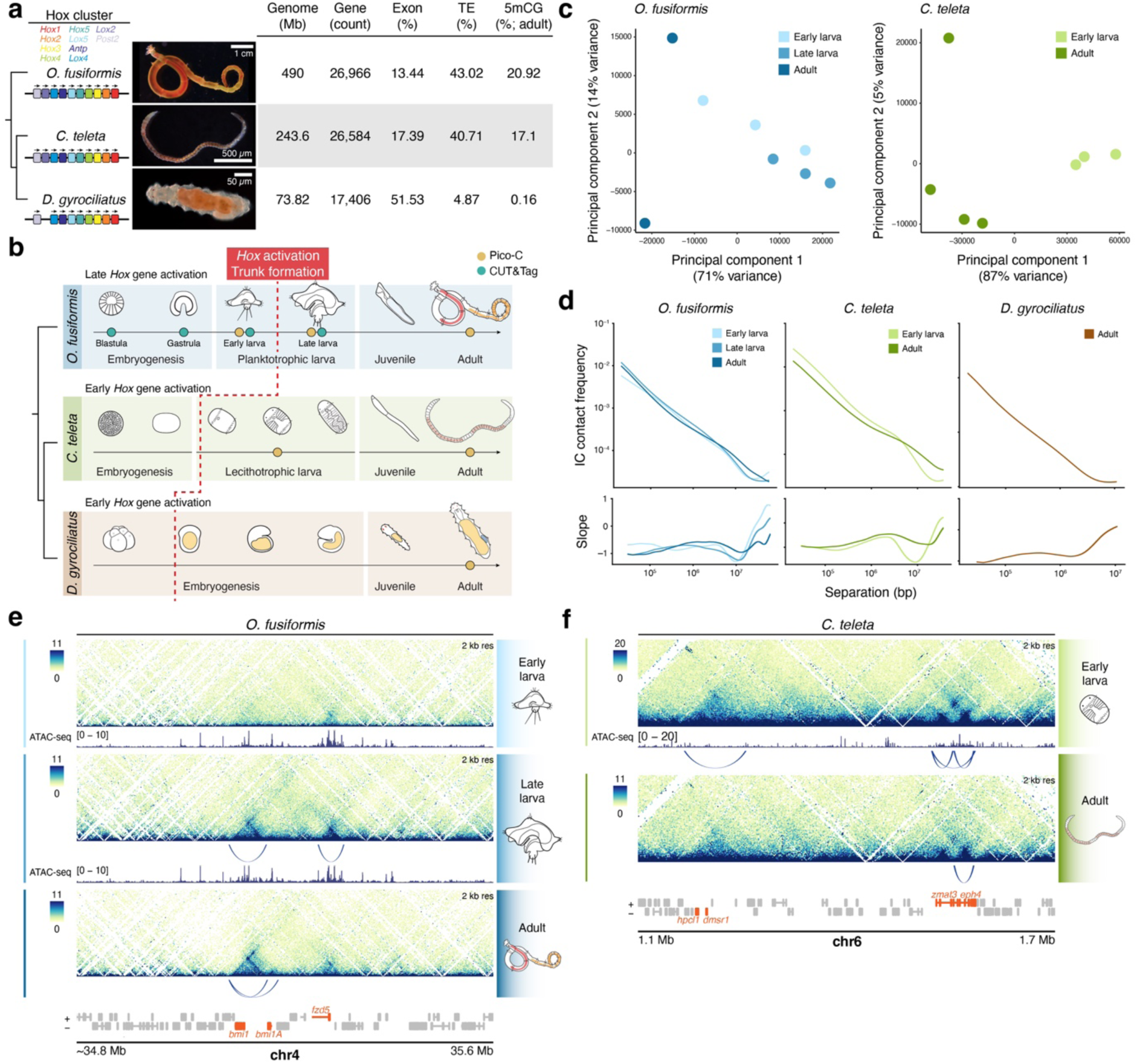
| The study of the annelid 3D chromatin landscape. (**a**) Species analysed in this study, their phylogenetic relationship, *Hox* gene complement (left), and basic genomic features (right). (**b**) Schematic drawings of the life cycle of *O. fusiformis*, *C. teleta*, and *D. gyrociliatus*, indicating the timing of activation of *Hox* genes and the samples taken for chromatin conformation analyses through Pico-C and histone modification profiling through CUT&Tag. (**c**) Principal component analysis segregates the different stages in O. fusiformis and C. teleta. (**d**) Top, line plots showing the rate of decay of contact frequency over genomic distance. The contact probability is averaged over all chromosomes. Bottom, log-derivative of the contact frequency probability plots. (**e**, **f**) Representative Pico-C chromatin interaction heatmaps of genomic regions with dynamic TADs and loops in the early and late larvae, as well as adults of *O. fusiformis* (**e**), and the early larva and adult of *C. teleta* (**f**). Genes contained in TADs and loops are highlighted in red. In (**e**, **f**), heatmaps represent normalised observed counts.

**Extended Data Figure 2.**
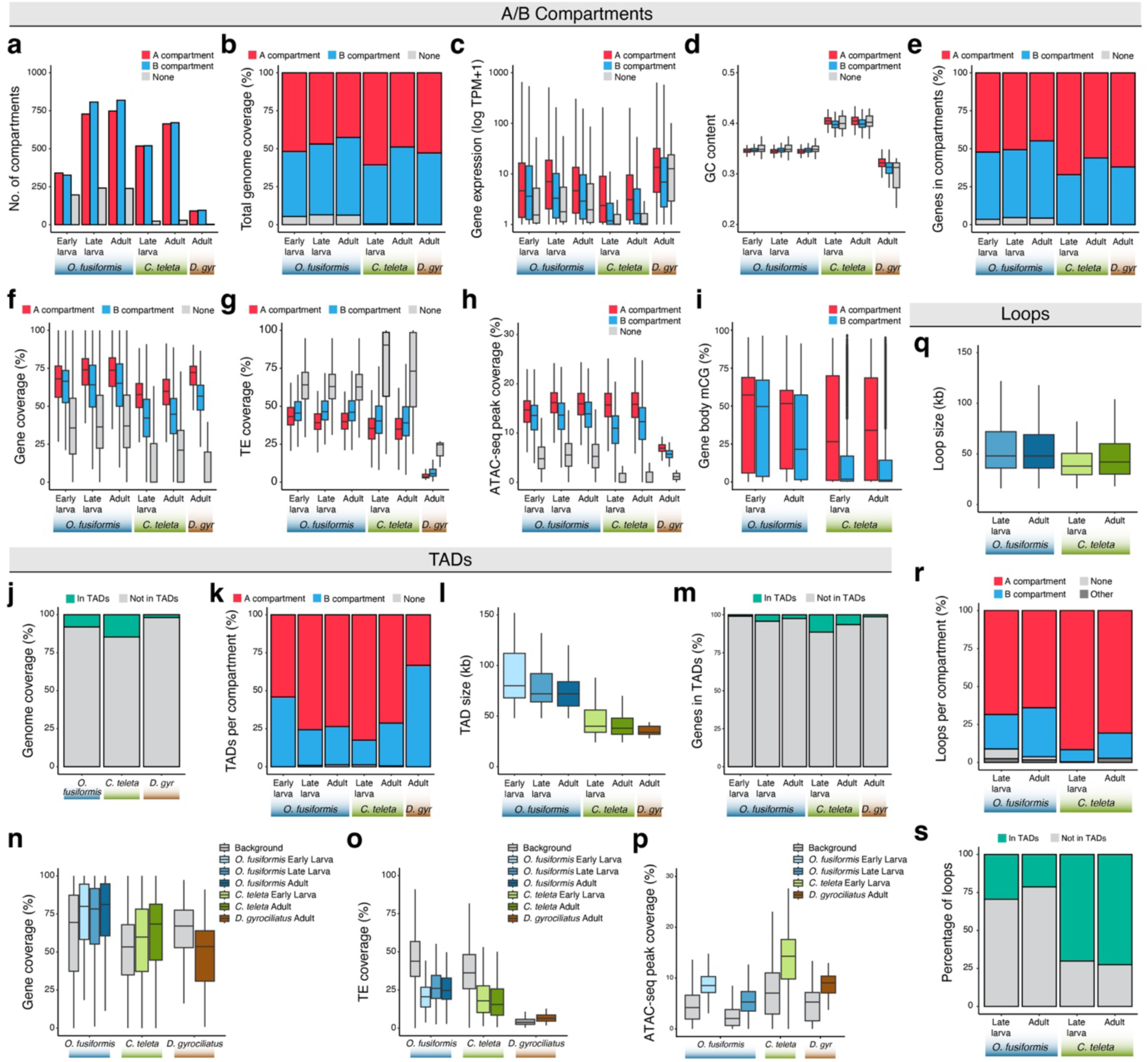
| Chromatin compartments, TADs, and chromatin loops in annelid genomes. (**a**) Bar plots indicating the number of A and B compartments and regions without any assignment in the studied time points. (**b**) Proportion of the genome occupied by A and B compartments. (**c**, **d**) Box plots showing gene expression levels (**c**) and GC content (**d**) in A and B compartments in *O. fusiformis*, *C. teleta* and *D. gyrociliatus*. (**e**) Proportion of genes located in A and B compartments. (**f**–**i**) Box plots showing the coverage of protein-coding genes (**f**), transposable elements (**g**), and open chromatin regions (**h**), as well as gene body methylation levels (**i**) in A and B compartments in *O. fusiformis*, *C. teleta* and *D. gyrociliatus*. (**j**) Proportion of the genome contained in TADs in *O. fusiformis*, *C. teleta* and *D. gyrociliatus*. (**k**) Proportion of TADs in A and B compartments. (**l**) Box plots depicting the size distributions of TADs at the analysed stages and species. (**m**) Proportion of genes in TADs. (**n**–**p**) Box plots showing the coverage of protein-coding genes (**n**), transposable elements (**o**), and open chromatin regions (**p**) in the TADs of *O. fusiformis*, *C. teleta* and *D. gyrociliatus*. (**q**) Box plots depicting the size distributions of chromatin loops at the analysed stages and species. (**r**, **s**) Bar plots showing the proportion of chromatin loops in A and B compartments (**r**) and in TADs (**s**) in *O. fusiformis* and *C. teleta*.

**Extended Data Figure 3.**
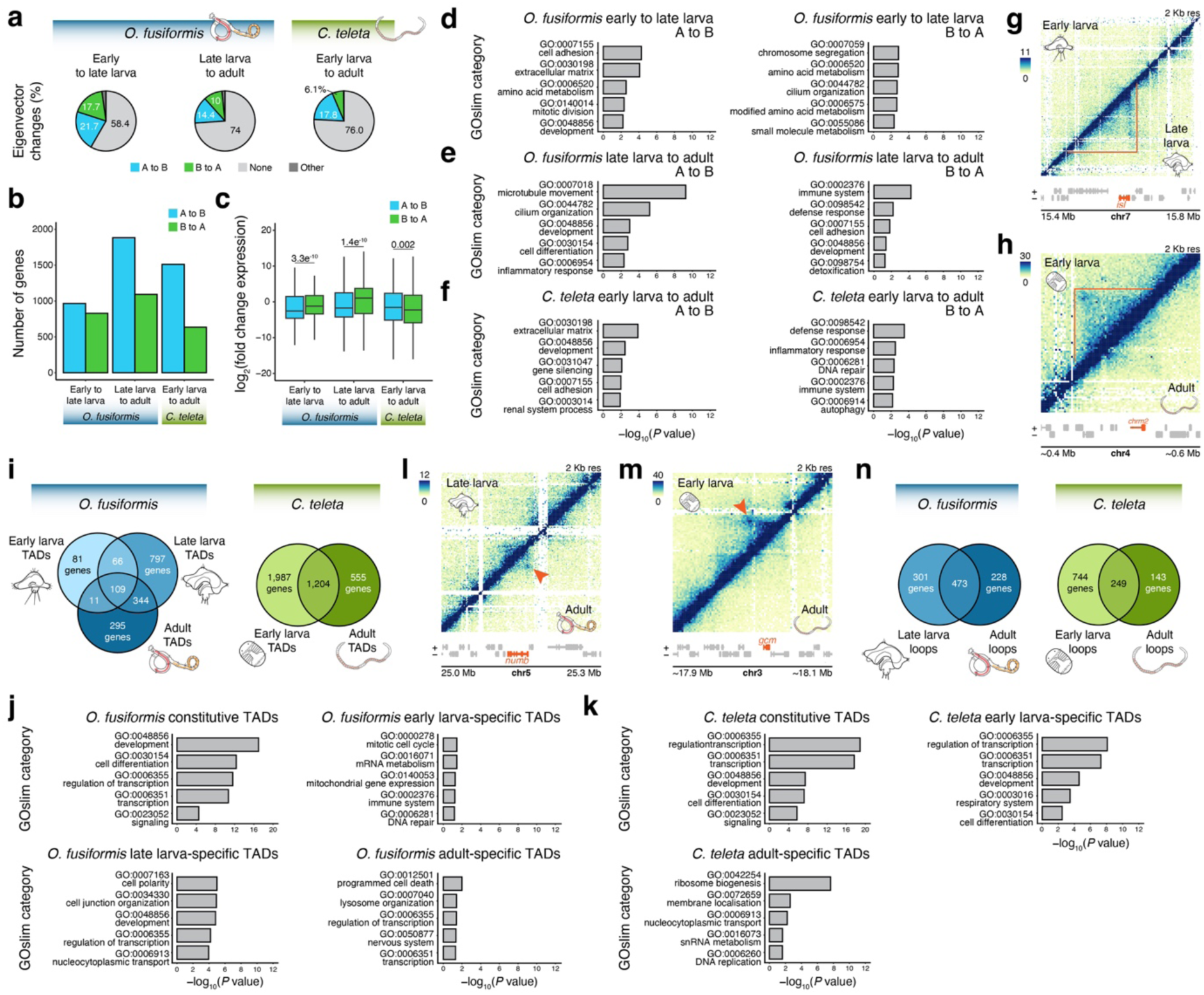
| The dynamic landscape of TADs and chromatin loops in annelids. (**a**) Pie charts depicting the proportion of compartment swaps between sequential pairwise comparisons of the studied time points in *O. fusiformis* and *C. teleta*. (**b**) Bar plots showing the number of genes changing compartment during the life cycles of *O. fusiformis* and *C. teleta*. (**c**) Box plots of the distribution of expression changes of differentially expressed genes that change compartment between stages in *O. fusiformis* and *C. teleta*. A swap to an A compartment is associated with a significantly higher fold change in expression in *O. fusiformis*, but the opposite is observed in *C. teleta*. *P* values are shown above the comparisons. (**d**–**f**) Bar plots showing the top five enriched Gene Ontology slim terms related to genes changing compartments between stages in *O. fusiformis* and *C. teleta*. (**g**, **h**) Representative Pico-C chromatin interaction heatmaps of genomic regions with dynamic TADs in O. fusiformis (**g**; *islet* locus) and *C. teleta* (**h**; *chrm2* locus). (**i**) Venn diagrams depicting the number of shared and stage-specific genes in TADs in *O. fusiformis* and *C. teleta*. (**l**, **m**) Representative Pico-C chromatin interaction heatmaps of genomic regions with dynamic loops in O. fusiformis (**g**; *numb* locus) and *C. teleta* (**h**; *gcm* locus). (**n**) Venn diagrams depicting the number of shared and stage-specific genes in chromatin loops in *O. fusiformis* and *C. teleta*. (**j**, **k**) Bar plots showing the top five enriched Gene Ontology slim terms related to genes that are always in TADs or at specific stages in *O. fusiformis* and *C. teleta*. In (**g**, **h**, **l**, **m**), heatmaps represent normalised observed counts. In (**d**–**f**, **j**, **k**), *P* values are obtained from two-tailed Fisher’s exact tests.

**Extended Data Figure 4.**
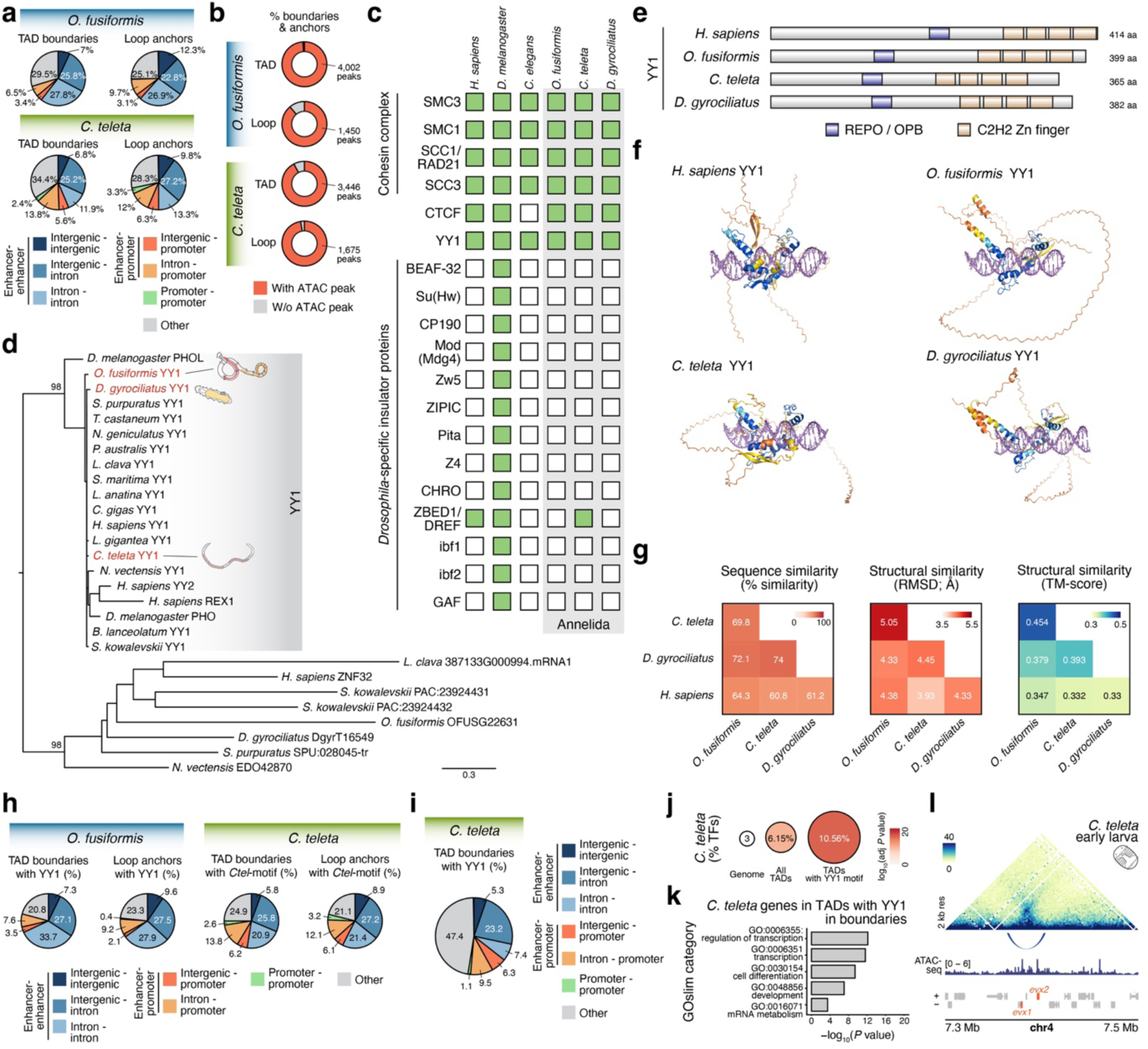
| The evolution of YY1 in Annelida. (**a**) Pie charts depicting the proportion of TAD boundaries and loop anchors that connect enhancers, enhancer and promoters, promoters, or other genomic regions in *O. fusiformis* and *C. teleta*. (**b**) Proportion of TAD boundaries and loop anchors that overlap at least one open chromatin (ATAC) peak. (**c**) Presence/absence heatmap of cohesin and insulator proteins in humans, *D. melanogaster*, *C. elegans*, and the three studied annelids. (**d**) Maximum likelihood phylogenetic tree of YY1 orthologs using related zinc fingers as the outgroup. (**e**) Schematic representation of YY1 proteins in human and the three annelids, indicating the four C2H2 zinc fingers and the Polycomb-interacting REPO domain. (**f**) AlphaFold reconstruction of those four YY1 proteins, showing the overall high conservation of the DNA-binding domains of the four proteins. (**g**) Heatmaps of sequence (left) and structural (centre and right) similarity scores between the human and annelid YY1 proteins. (**h**, **i**) Pie charts illustrating the genomic annotation of TAD boundaries and loop anchors with the YY1-like motif in *O. fusiformis* and the species-specific motif in *C. teleta* (**h**) and the TAD boundaries with the YY1-like motif in *C. teleta* (**i**). (**j**, **k**) TADs with the YY1-like motif in *C. teleta* are highly enriched for transcription factors (**j**; two-tailed Fisher’s exact test with Bonferroni correction) and Gene Ontology slim terms related to transcriptional regulation and development (**k**; two-tailed Fisher’s exact test). (**l**) Pico-C chromatin interaction heatmap of the *evx* locus in *C. teleta*. The anchors of the chromatin loop contain YY1-like motifs, and the loop brings the two paralogs together.

**Extended Data Figure 5.**
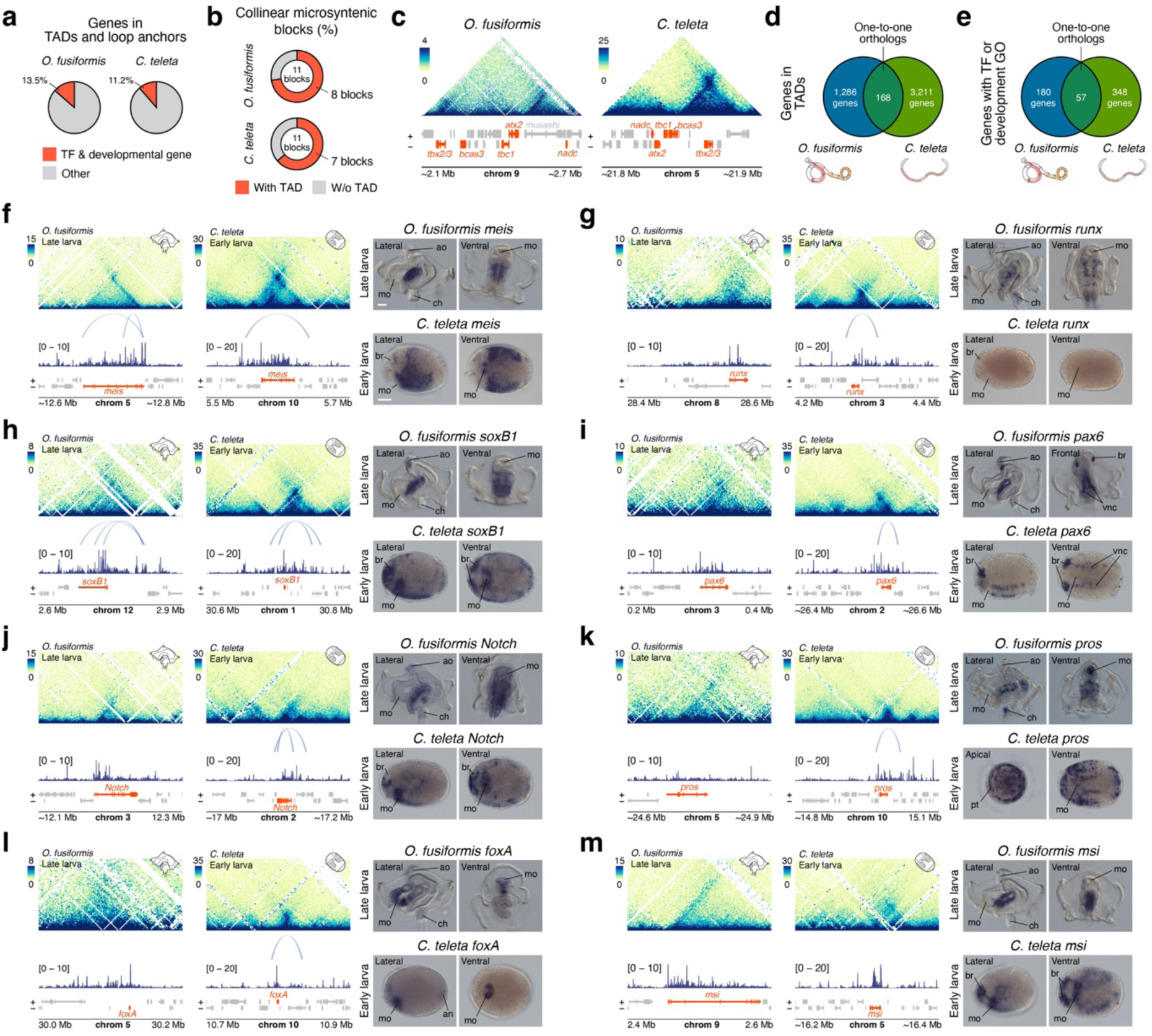
| The evolution of 3D chromatin regulation in developmental genes in Annelida. (**a**) Proportion of genes in TADs and loop anchors that are transcription factors and developmental genes in *O. fusiformis* and *C. teleta*. (**b**) Proportion of collinear microsyntenic blocks between *O. fusiformis* and *C. teleta* with a TAD in each species. (**c**) Representative Pico-C chromatin interaction heatmap around a collinear microsyntenic block in *O. fusiformis* and *C. teleta*. (**d**, **e**) Venn diagrams representing the number of shared one-to-one orthologs in TADs between *O. fusiformis* and *C. teleta* (**d**) and shared one-to-one orthologs that are either transcription factors or developmental genes (**e**). (**f**–**m**) For each panel on the left, Pico-C chromatin interaction heatmaps around orthologous developmental genes (*meis*, *runx*, *soxB1*, *pax6*, *Notch*, *pros*, *foxA*, and *msi*) in *O. fusiformis* and *C. teleta*, depicting chromatin loops and open chromatin data below. On the right, whole-mount *in situ* hybridisation of each gene at corresponding developmental stages, namely the late larva of *O. fusiformis* and the early larva of *C. teleta*. (**f**) *meis* is expressed in the trunks of *O. fusiformis* and *C. teleta*, as well as the brain and mouth of *C. teleta*. (**g**) In the late larva of *O. fusiformis*, *runx* is expressed in isolated cells of the ventral nerve cord. In the early larva of *C. teleta* (stage 5), *runx* is weakly expressed and mainly detected in the brain^42^. At later larval stages, the expression is, however, like that of *O. fusiformis*^42^. (**h**) *soxB1* is expressed in the brain, trunk and ventral nerve cords in the two annelids, as well as in the mouth of *C. teleta*. (**i**) *pax6* is detected in the annelid brain and ventral nerve cord. (**j**) *Notch* is expressed in the trunks of *O. fusiformis* and *C. teleta*, as well as in the anterior ectoderm, brain and mouth of *C. teleta*. (**k**) *prospero* (*pros*) is expressed in isolated ectodermal cells, mostly restricted to the trunk in *O. fusiformis* but more broadly distributed in *C. teleta*. (**l**) *foxA* is expressed in the foregut in *O. fusiformis* and *C. teleta*, as well as in the hindgut in the latter. (**f**) *musashi* (*msi*) is detected in the annelid trunk, as well as brain and mouth in *C. teleta*. In (**c**, **f**–**m**), heatmaps represent normalised observed counts. an, anus; ao, apical organ; br, brain; ch, chaetae; mo, mouth; pt, prototroch; vnc, ventral nerve cord. Scale bars in (**f**) are 50 µm and apply to panels (**f**–**m**).

**Extended Data Figure 6.**
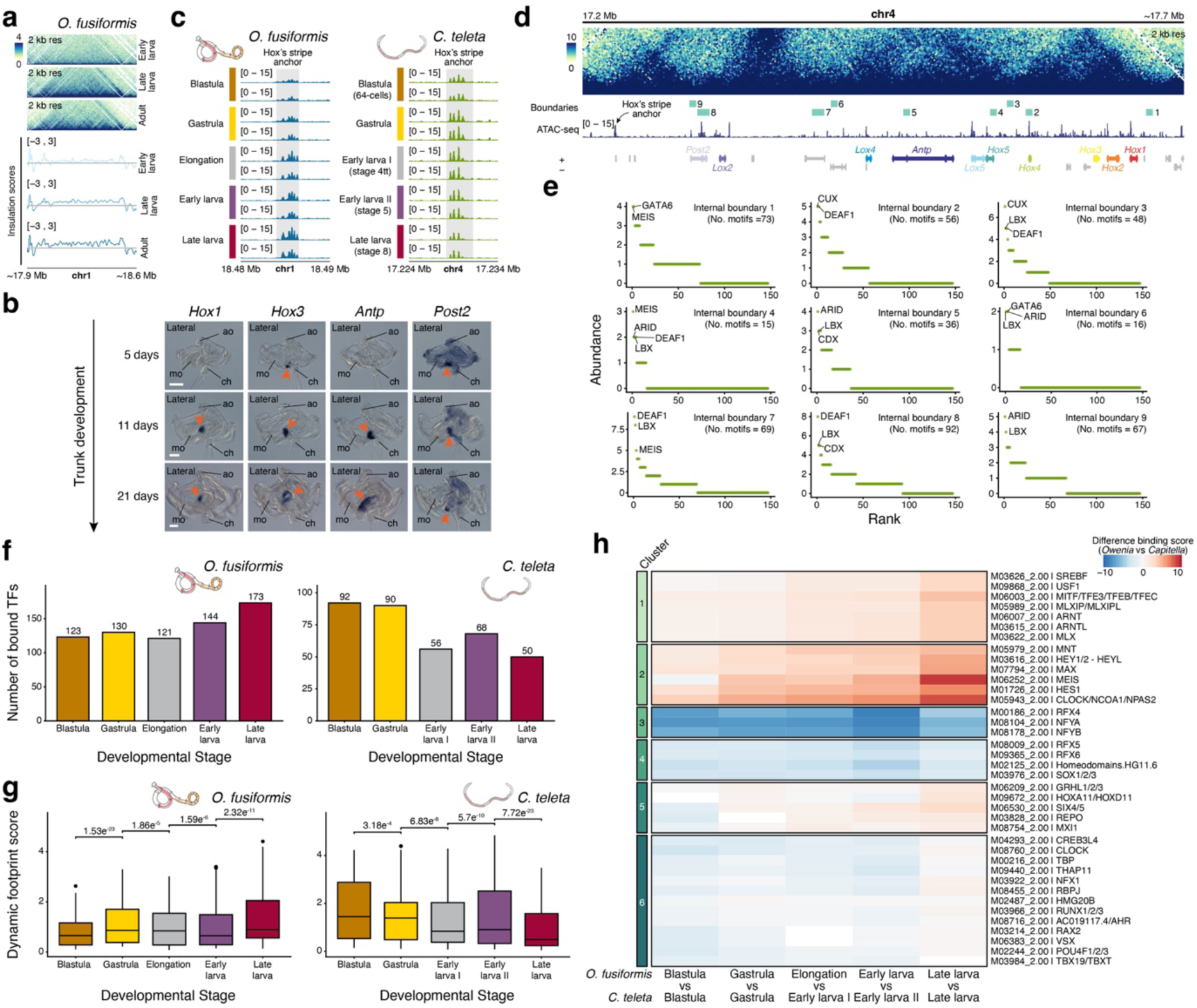
| *Hox* gene regulation in annelids. (**a**) On top, Pico-C contact heatmap of normalised observed counts in the Hox cluster of *O. fusiformis* at the early and late larval stages and adults. Bottom, insulation scores in that region at those stages, showing the increase in self-insulation of the Hox cluster as *Hox* genes become active. (**b**) Whole-mount *in situ* hybridisation of *Hox1*, *Hox3*, *Antp*, and *Post2* at three time points during larval growth in *O. fusiformis*. At 5 days post fertilisation (dpf), *Hox3* and *Post2* are expressed at the ventral ectoderm where the infolding of the trunk rudiment is forming (red arrowheads). At 11 dpf, when the trunk rudiment is well-formed, and in the late competent larva at 21 dpf, when the trunk is fully differentiated, these four *Hox* genes are expressed in a spatially collinear fashion (red arrowheads). (**c**) For each species, the tracks show chromatin accessibility dynamics at the stripe anchors of the Hox clusters. (**d**) Pico-C contact heatmap of normalised observed counts in the Hox cluster of *C. teleta*, indicating the internal boundaries in relation to *Hox* genes. (**e**) Scattered plots of motif count in each internal boundary of *C. teleta*’s Hox cluster. (**f**) Bar plots indicating the number of bound transcription factors in the stripe anchor of the Hox clusters of *O. fusiformis* and *C. teleta*. (**g**) Boxplots depicting the distributions of footprinting scores at the Hox cluster’s stripe anchor in *O. fusiformis* and *C. teleta*. In *O. fusiformis*, there are more transcription factors and more strongly bound in the late larval stage, when *Hox* genes are expressed, whereas the opposite occurs in *C. teleta*. (**h**) Clustered heatmap of the differential footprinting scores of shared bound transcription factors in the stripe anchors of the Hox clusters of *O. fusiformis* and *C. teleta*. Clusters 1 and 2 include transcription factors that are generally more strongly bound in *O. fusiformis* than in *C. teleta*. Clusters 3 and 4 comprise transcription factors that are generally more bound in *C. teleta* than in *O. fusiformis*. Clusters 5 and 6 include transcription factors that are more strongly bound at earlier developmental time points in *C. teleta* but more strongly bound at late larval stages in *O. fusiformis*. ao, apical organ; ch, chaetae; mo, mouth. Scale bars in (**a**) are 50 µm.

**Extended Data Figure 7.**
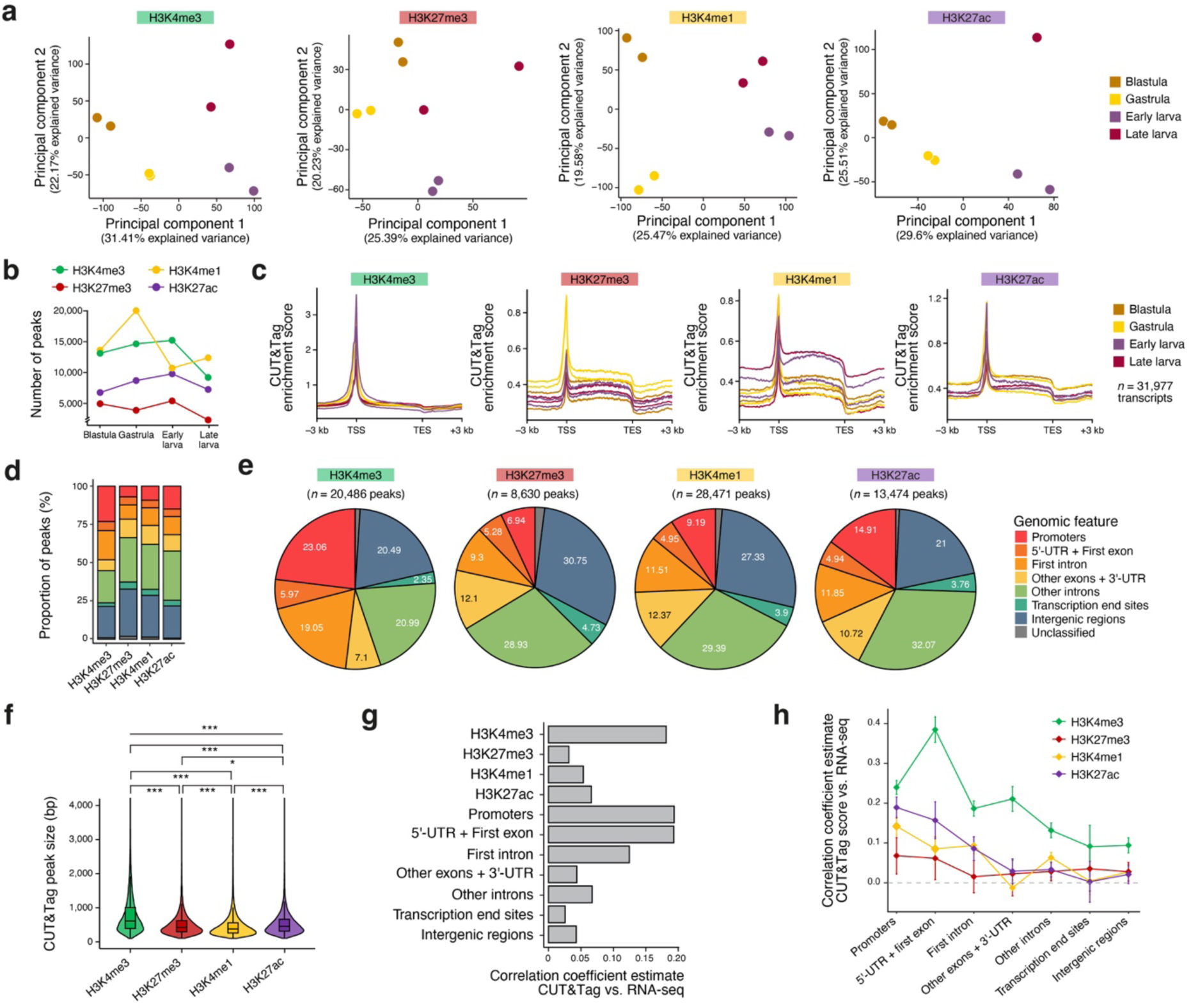
| Histone post-translation modification profiling in *O. fusiformis*. (**a**) Principal component analysis of the profiled histone marks at the blastula, gastrula, early and late larval stages. Duplicates are highly correlated. (**b**) Peak counts during embryonic and larval development. (**c**) Per-replicate metagene profiles for H3K4me3, H3K27me3, H3K4me1, and H3K27ac and 3 kb flanking regions upstream of the transcription start site (TSS) and downstream of the transcription end site (TES). (**d**, **e**) Bar plots (left) and pie charts (right) depicting the genomic annotation of the consensus peaks for each profiled histone mark. (**f**) Violin and box plots showing the size distributions of consensus peaks for H3K4me3, H3K27me3, H3K4me1, and H3K27ac. (**g**) Correlation coefficients between CUT&Tag and RNA-seq data for each histone mark and genomic location. (**h**) Correlation coefficients between CUT&Tag and RNA-seq data at different genomic locations for each of the profiled histone marks.

**Extended Data Figure 8.**
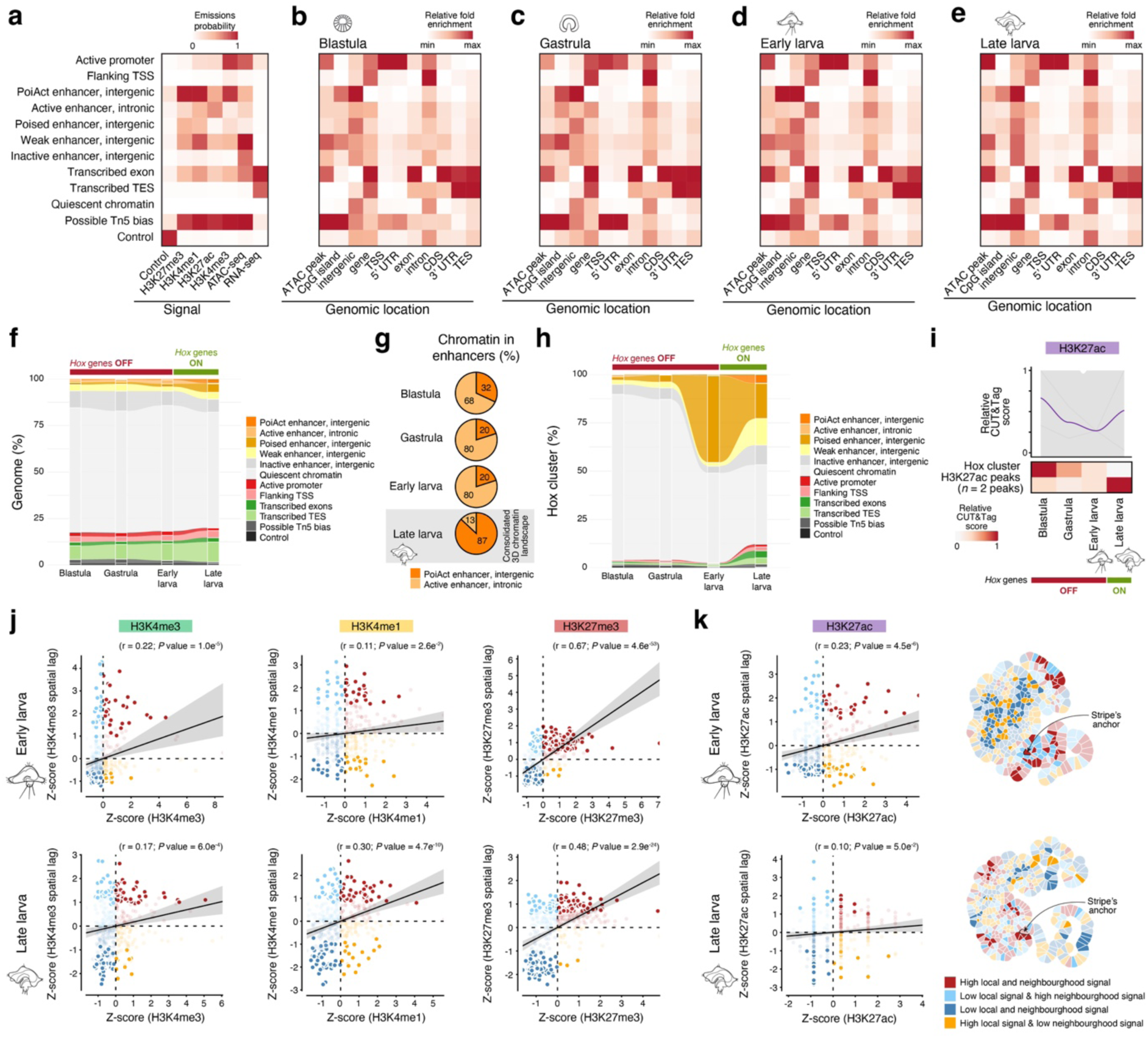
| Chromatin state dynamics during *O. fusiformis* embryogenesis. (**a**–**e**) Heatmaps of emission probabilities (**a**) and genomic enrichment of the 12 (including Tn5 bias and control) chromatin states defined by integrating histone profiling (CUT&Tag) data for H3K4me3, H3K4me1, H3K27me3, and H3K27ac, open chromatin data (ATAC-seq), and gene expression (RNA-seq) at the blastula (**b**), gastrula (**c**), early larval (**d**) and late larval (**e**) stages. (**f**) Alluvial plot of the proportion of each regulatory state, including the potential Tn5 and control states, at the blastula, gastrula, early larval and late larval stages. (**g**) Pie charts depicting the proportion of strongly active intergenic and intronic enhancers during *O. fusiformis*’ life cycle. There is a shift from mostly intronic to mostly intergenic active enhancers as the 3D regulatory landscape consolidates in the late larva. (**h**) Alluvial plot of the proportion of the Hox cluster of *O. fusiformis* exhibiting each chromatin state, including the potential Tn5 and control states, during embryogenesis and larval growth. (**i**) Dynamics of CUT&Tag scores in the H3K27ac peaks located in the Hox cluster during embryogenesis and larval development. On top, the regression line is coloured. (**j**) Scattered plots correlating local (x axis) and neighbourhood (y axis) signal at each node in the Kamada-Kawai graph layout of the Hox cluster in early (top row) and late (bottom row) larva. Solid colours indicate statistically significant correlations (*P* value < 0.05). (**k**) METALoci integration of H3K27ac signal with Pico-C data in the Hox cluster. On the left, scattered plots correlating local (x axis) and neighbourhood (y axis) signal at each node in the Kamada-Kawai graph layout of the Hox cluster in early (top row) and late (bottom row) larva. On the right, Gaudí plots projecting the H3K27ac signal onto a two-dimensional Kamada-Kawai graph layout. In all cases, solid colours indicate statistically significant (*P* value < 0.05) 2 Kb bins identified using a one-sided permutation test.

**Extended Data Figure 9.**
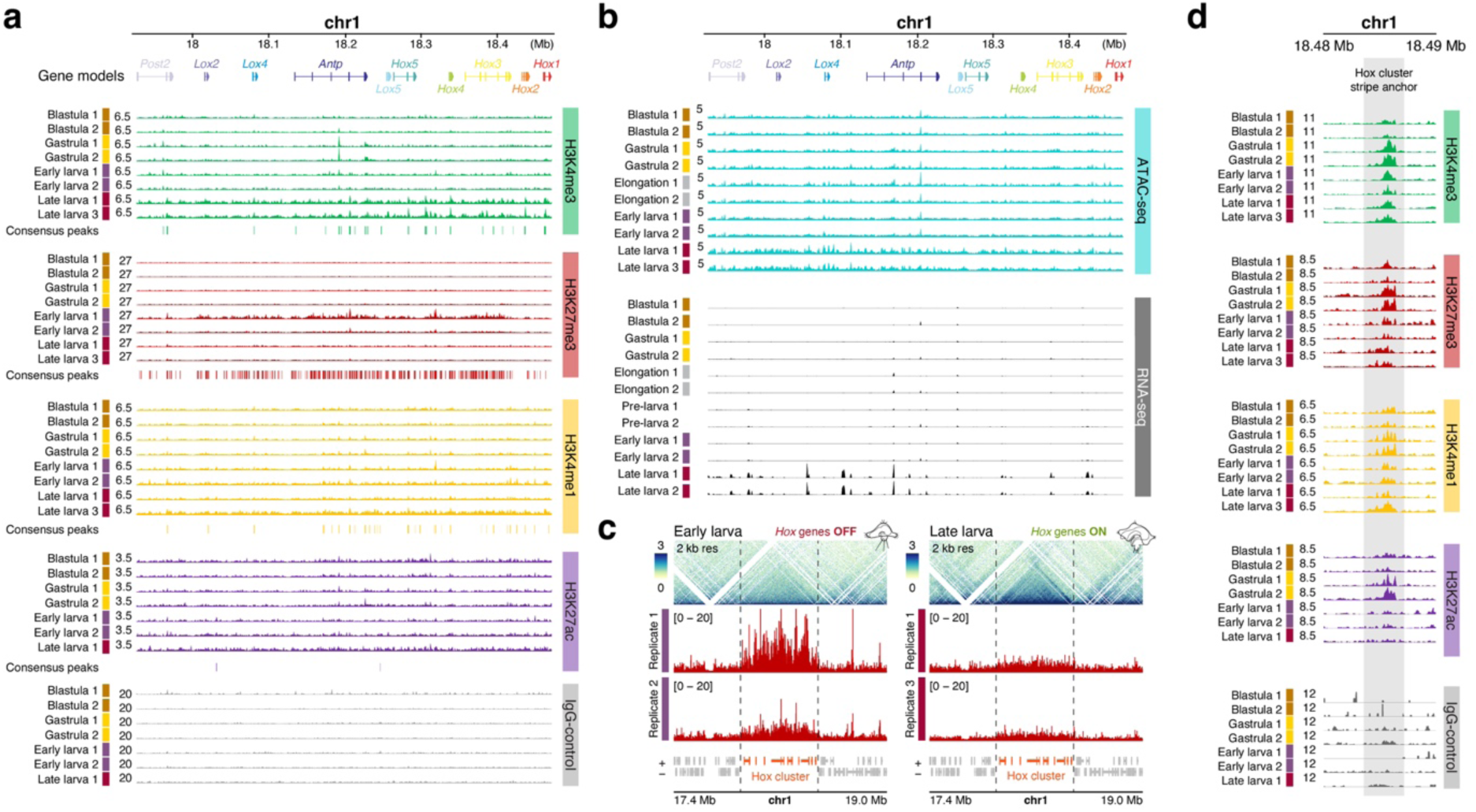
| The dynamics of histone post-translation modifications in the Hox cluster of *O. fusiformis*. (**a**, **b**) Tracks depicting the abundance of the studied histone modifications (**a**), as well as open chromatin (ATAC-seq data) and transcription (RNA-seq data) (**b**) across the Hox cluster of *O. fusiformis*. (**c**) Zoom outs of the genomic region comprising the Hox cluster in the early (left) and late (right) larval stages in *O. fusiformis*, depicting the Pico-C contact heatmaps of normalised observed counts on the top and the abundance of the Polycomb-mediated H3K27me3 repressive mark on the bottom. Not the reduction of the repressive mark as *Hox* genes become active. (**d**) Tracks depicting the abundance of the studied histone modifications at the Hox cluster’s stripe anchor during *O. fusiformis* embryonic development and larval growth.

