## Supplementary Figure for "Convergent evolution of cluster-wide *Hox* gene regulation in Bilateria"

**Supplementary Information includes:**

- 20 Supplementary Figures
- 67 Supplementary Tables (as an Excel spreadsheet)

**Index of Supplementary Figures:**

- **Supplementary Figure 1 |** Pico-C quality check.
- **Supplementary Figure 2 |** The new genome assemblies of *C. teleta* and *D. gyrociliatus*.
- **Supplementary Figure 3 |** Chromatin compartmentalisation in the early larva of *O. fusiformis*.
- **Supplementary Figure 4 |** Chromatin compartmentalisation in the late larva of *O. fusiformis*.
- **Supplementary Figure 5 |** Chromatin compartmentalisation in the adult of *O. fusiformis*.
- **Supplementary Figure 6 |** Chromatin compartmentalisation in the early larva of *C. teleta*.
- **Supplementary Figure 7 |** Chromatin compartmentalisation in the adult of *C. teleta*.
- **Supplementary Figure 8 |** Chromatin compartmentalisation in the adult of *D. gyrociliatus*.
- **Supplementary Figure 9 |** Chromatin compartments and gene expression in annelids.
- **Supplementary Figure 10 |** 3D chromatin architecture and DNA methylation in *O. fusiformis* and *C. teleta*.
- **Supplementary Figure 11 |** YY1 evolution in annelids.
- **Supplementary Figure 12 |** Multi-species full-genome alignment.
- **Supplementary Figure 13 |** CUT&Tag library statistics.
- **Supplementary Figure 14 |** CUT&Tag insert size distributions.
- **Supplementary Figure 15 |** Reproducibility assessment and peak calling benchmarking.
- **Supplementary Figure 16 |** CUT&Tag enrichment around hPTM peaks.
- **Supplementary Figure 17 |** Correlation and developmental dynamics of histone marks.
- **Supplementary Figure 18 |** Histone mark enrichment in *O. fusiformis* genes.
- **Supplementary Figure 19 |** Developmental dynamics of histone marks.
- **Supplementary Figure 20 |** Correlation between hPTM dynamics and gene expression dynamics.

**Index of Supplementary Tables and Legends**

- **Supplementary Table 1 |** Sequencing and mapping statistics for Pico-C libraries.
- **Supplementary Table 2 |** Annelid compartments and associated genomic features.
- **Supplementary Table 3 |** Annelid TADs and associated genomic features.
- **Supplementary Table 4 |** Annelid TADs and ATAC-seq coverage.
- **Supplementary Table 5 |** Gene expression in annelid TADs.
- **Supplementary Table 6 |** Gene content in the TADs of *O. fusiformis* (only considering genes overlapping at least 50% with the TAD span).
- **Supplementary Table 7 |** Gene content in the TADs of *C. teleta* (only considering genes overlapping at least 50% with the TAD span).
- **Supplementary Table 8 |** Gene content in the loop anchors of *O. fusiformis*.
- **Supplementary Table 9 |** Gene content in the loop anchors of *C. teleta*.
- **Supplementary Table 10 |** Genes changing compartments in *O. fusiformis*.
- **Supplementary Table 11 |** Genes changing compartments in *C. teleta*.
- **Supplementary Table 12 |** Transcription factor annotation in *O. fusiformis*.
- **Supplementary Table 13 |** Transcription factor annotation in *C. teleta*.
- **Supplementary Table 14 |** Gene Ontology term annotation in *O. fusiformis*.
- **Supplementary Table 15 |** Gene Ontology term annotation in *C. teleta*.
- **Supplementary Table 16 |** List of Gene Ontology terms used to annotate developmental genes.
- **Supplementary Table 17 |** One-to-one ortholog genes between *O. fusiformis* and *C. teleta*.
- **Supplementary Table 18 |** Gene age annotation in *O. fusiformis*.
- **Supplementary Table 19 |** Gene age annotation in *C. teleta*.
- **Supplementary Table 20 |** Genomic feature annotation of *O. fusiformis*’ TAD boundaries.
- **Supplementary Table 21 |** Genomic feature annotation of *O. fusiformis*’ loop anchors.
- **Supplementary Table 22 |** Genomic feature annotation of *C. teleta*’s TAD boundaries.
- **Supplementary Table 23 |** Genomic feature annotation of *C. teleta*’s loop anchors.
- **Supplementary Table 24 |** Genomes used in sequence conservation analyses.
- **Supplementary Table 25 |** PhastCons values in annelid TAD boundaries and loop anchors.
- **Supplementary Table 26 |** Transcription factor motif transfer for *O. fusiformis*.
- **Supplementary Table 27 |** Transcription factor motif transfer for *C. teleta*.
- **Supplementary Table 28 |** Transcription factor motif transfer for *D. gyrociliatus*.
- **Supplementary Table 29 |** Annotation of YY1-like motif in TAD boundaries in *O. fusiformis*.
- **Supplementary Table 30 |** Annotation of YY1-like motif in loop anchors in *O. fusiformis*.
- **Supplementary Table 31 |** Annotation of the *C. teleta*-specific motif in TAD boundaries in *C. teleta*.
- **Supplementary Table 32 |** Annotation of the *C. teleta*-specific motif in loop anchors in *C. teleta*.
- **Supplementary Table 33 |** Annotation of YY1-like motif in TAD boundaries in *C. teleta*.
- **Supplementary Table 34 |** Motif enrichment in annelid TAD boundaries and loop anchors.
- **Supplementary Table 35 |** Motif annotation in the stripe anchor of the Hox cluster in *O. fusiformis*.
- **Supplementary Table 36 |** Motif annotation in the stripe anchor of the Hox cluster in *C. teleta*.
- **Supplementary Table 37 |** Motif annotation in the internal boundaries of the Hox cluster in *C. teleta*.
- **Supplementary Table 38 |** Motif footprinting in the stripe anchor of the Hox cluster in *O. fusiformis*.
- **Supplementary Table 39 |** Motif footprinting in the stripe anchor of the Hox cluster in *C. teleta*.
- **Supplementary Table 40 |** Footprinting scores of shared motifs in the stripe anchors of the Hox clusters of *O. fusiformis* and *C. teleta*.
- **Supplementary Table 41 |** Conserved microsyntenic collinear blocks between *O. fusiformis* and *C. teleta*.
- **Supplementary Table 42 |** Tn5 transposome oligonucleotides and sequencing primers used for CUT&Tag experiments.
- **Supplementary Table 43 |** Primers and sequencing indices used for each CUT&Tag library.
- **Supplementary Table 44 |** Read duplication statistics for CUT&Tag libraries.
- **Supplementary Table 45 |** Sequencing and mapping statistics for CUT&Tag experiments.
- **Supplementary Table 46 |** MACS2 peak calling statistics for CUT&Tag experiments.
- **Supplementary Table 47 |** IDR peak set calling statistics for CUT&Tag experiments.
- **Supplementary Table 48 |** FRiP score-based peak calling benchmarking in hPTM-specific DiffBind consensus peak sets for CUT&Tag experiments
- **Supplementary Table 49 |** DiffBind consensus peak set calling statistics for CUT&Tag experiments.
- **Supplementary Table 50 |** Different peak calling merging.
- **Supplementary Table 51 |** Number of peaks after ATAC-seq-based filtering.
- **Supplementary Table 52 |** Developmental dynamics of normalised DiffBind H3K4me3 CUT&Tag score in *O. fusiformis*.
- **Supplementary Table 53 |** Developmental dynamics of normalised DiffBind H3K27me3 CUT&Tag score in *O. fusiformis*.
- **Supplementary Table 54 |** Developmental dynamics of normalised DiffBind H3K4me1 CUT&Tag score in *O. fusiformis*.
- **Supplementary Table 55 |** Developmental dynamics of normalised DiffBind H3K27ac CUT&Tag score in *O. fusiformis*.
- **Supplementary Table 56 |** Genomic feature and nearest gene annotation of the H3K4me3 DiffBind consensus peak set.
- **Supplementary Table 57 |** Genomic feature and nearest gene annotation of the H3K27me3 DiffBind consensus peak set.
- **Supplementary Table 58 |** Genomic feature and nearest gene annotation of the H3K4me1 DiffBind consensus peak set.
- **Supplementary Table 59 |** Genomic feature and nearest gene annotation of the H3K27ac DiffBind consensus peak set.
- **Supplementary Table 60 |** Statistics from comparison analysis of hPTM peak size.
- **Supplementary Table 61 |** ANOVA table for the bivariate linear model based on hPTM and genomic feature.
- **Supplementary Table 62 |** Coefficient estimates table for the bivariate linear model based on hPTM and genomic feature.
- **Supplementary Table 63 |** CUT&Tag peaks and their normalised score dynamics located in the *Hox* genes cluster locus of *O. fusiformis*.
- **Supplementary Table 64 |** Chromatin states emission probabilities for *O. fusiformis*.
- **Supplementary Table 65 |** Fold enrichment of chromatin states in different genomic contexts at the blastula, gastrula, early and late larval stages of *O. fusiformis*.
- **Supplementary Table 66 |** Genome assemblies and Hi-C datasets of spiralian species used to explore the evolution of the Hox cluster chromatin architecture.
- **Supplementary Table 67 |** Gene IDs of orthologous genes used in gene expression analyses in *O. fusiformis* and *C. teleta*.

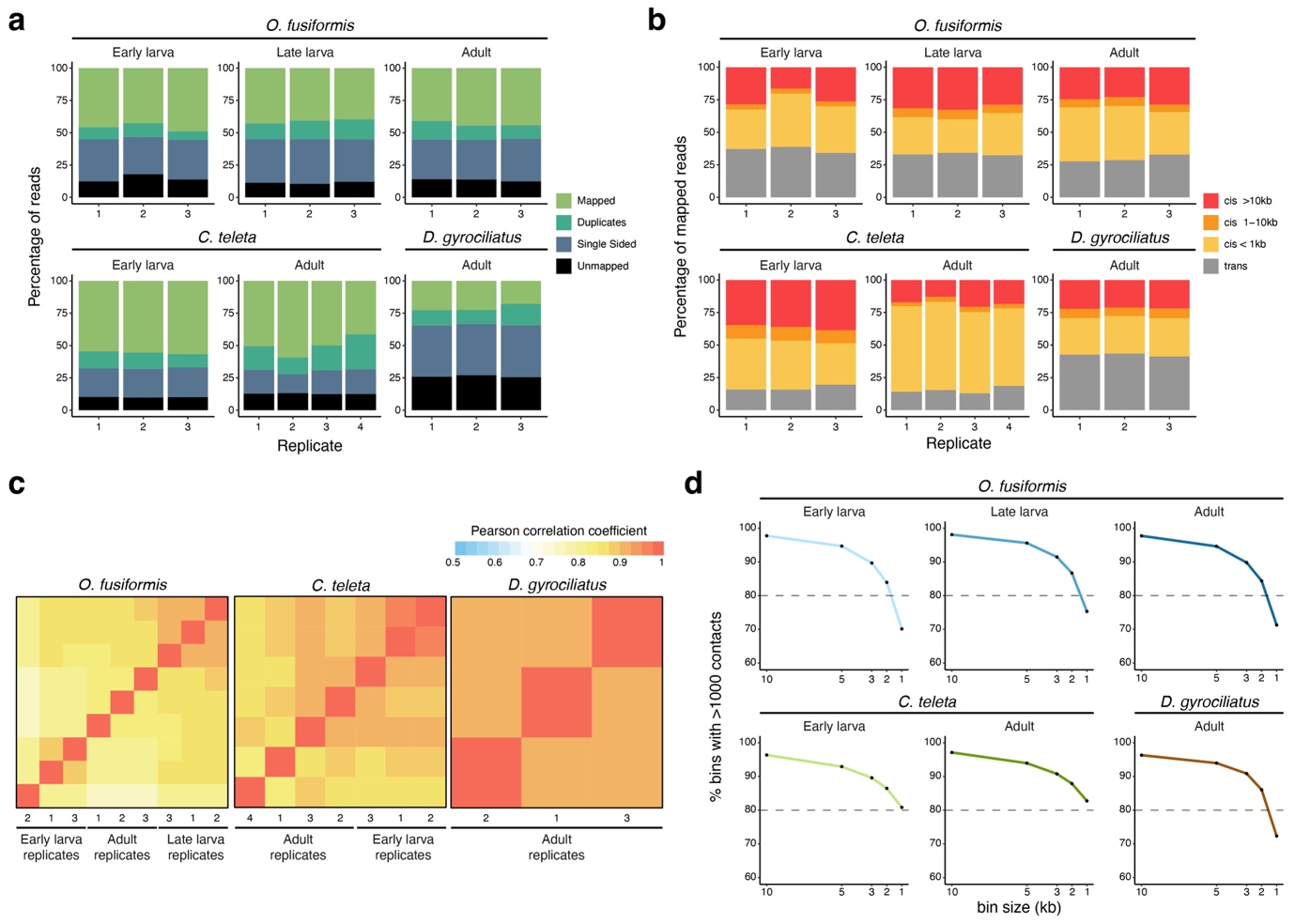

**Supplementary Figure 1** **| Pico-C quality check.** (**a**, **b**) Bar plots showing the percentages of mapped reads (**a**) and mapping patterns (**b**) per stage and replicate. (**c**) Heatmaps of Pearson correlation coefficients between replicates. (**d**) Line plots showing library complexity at different resolutions, used as an estimate of the minimal ideal resolution (bin size with > 80% of bins having more than 1000 contacts).

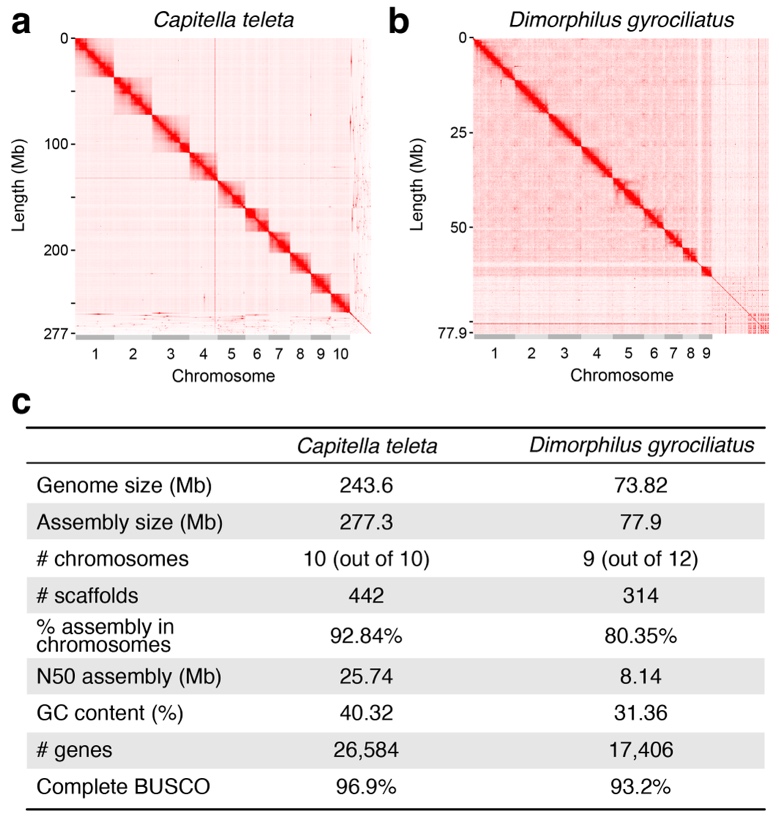

**Supplementary Figure 2 | The new genome assemblies of *C. teleta* and *D. gyrociliatus*.** (**a**, **b**) Assembly-wide contact maps showing the new chromosome-level scaffolding of *the C. teleta* and *D. gyrociliatus* genomes. The new assembly resolves chromosome 3 in *C. teleta* and assembles nine of the 12 chromosomes in *D. gyrociliatus*. (**c**) Table of the new assembly statistics.

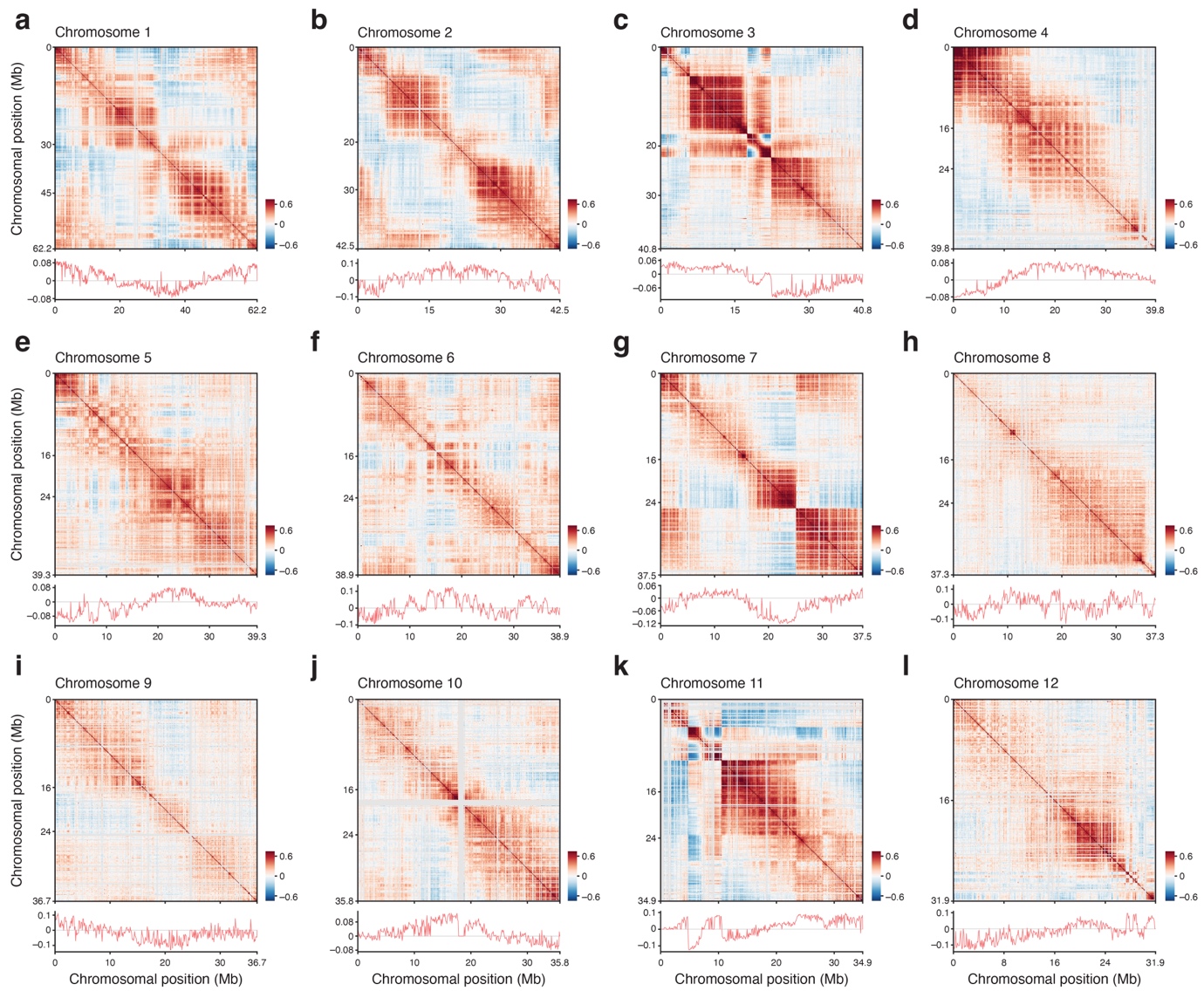

**Supplementary Figure 3 | Chromatin compartmentalisation in the early larva of *O. fusiformis*.** (**a**–**l**) For each panel, the top shows a Pico-C contact heatmap for each chromosome. The bottom panel shows a line plot of eigenvalues along the chromosome.

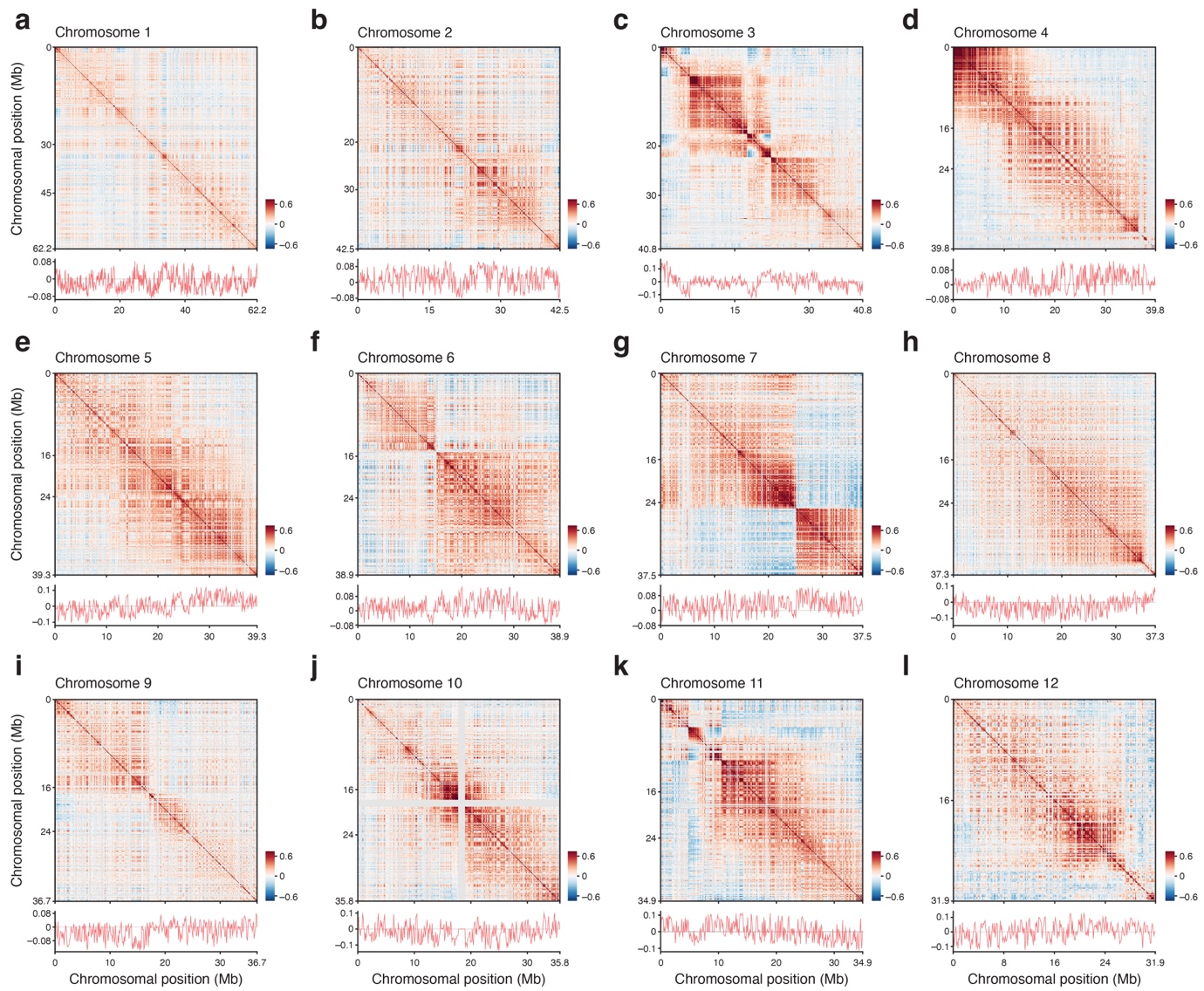

**Supplementary Figure 4 | Chromatin compartmentalisation in the late larva of *O. fusiformis*.** (**a**–**l**) For each panel, the top shows the Pico-C contact heatmap for each chromosome. The bottom panel shows a line plot of eigenvalues along the chromosome.

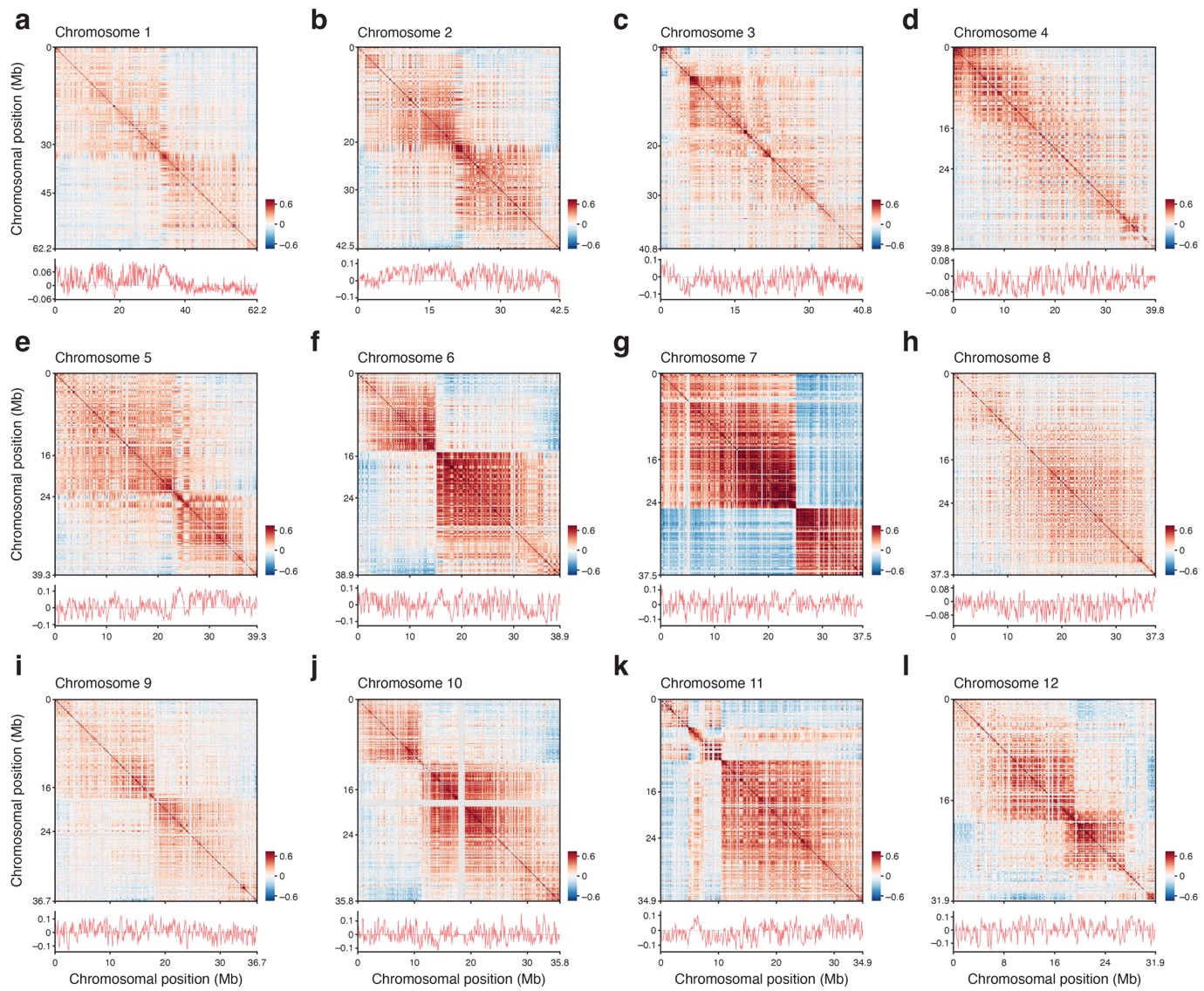

**Supplementary Figure 5 | Chromatin compartmentalisation in the adult of *O. fusiformis*.** (**a**–**l**) For each panel, the top shows the Pico-C contact heatmap for each chromosome. The bottom panel shows a line plot of eigenvalues along the chromosome.

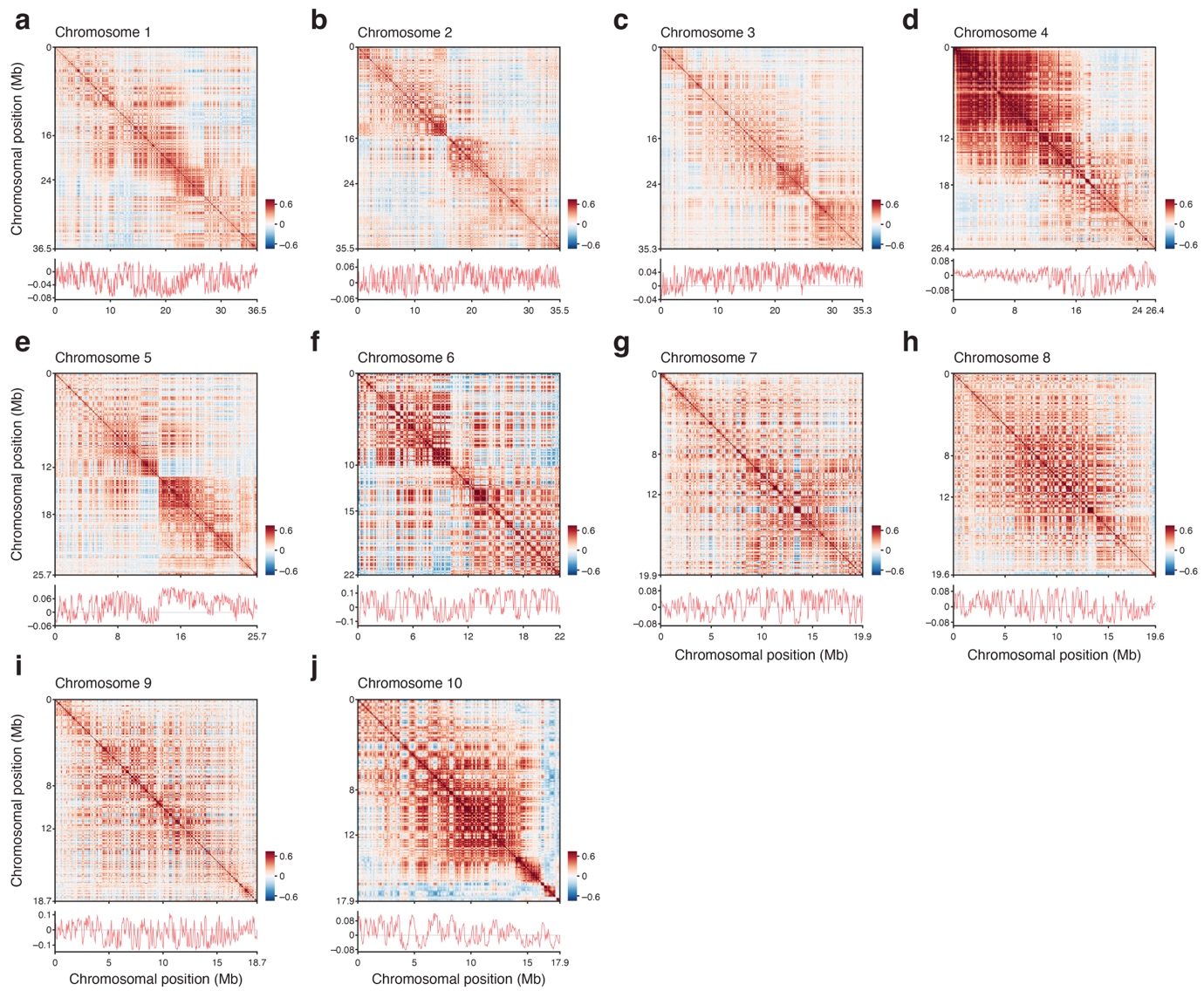

**Supplementary Figure 6 | Chromatin compartmentalisation in the early larva of *C. teleta*.** (**a**–**j**) For each panel, a Pico-C contact heatmap for each chromosome is shown at the top. The bottom panel shows a line plot of eigenvalues along the chromosome.

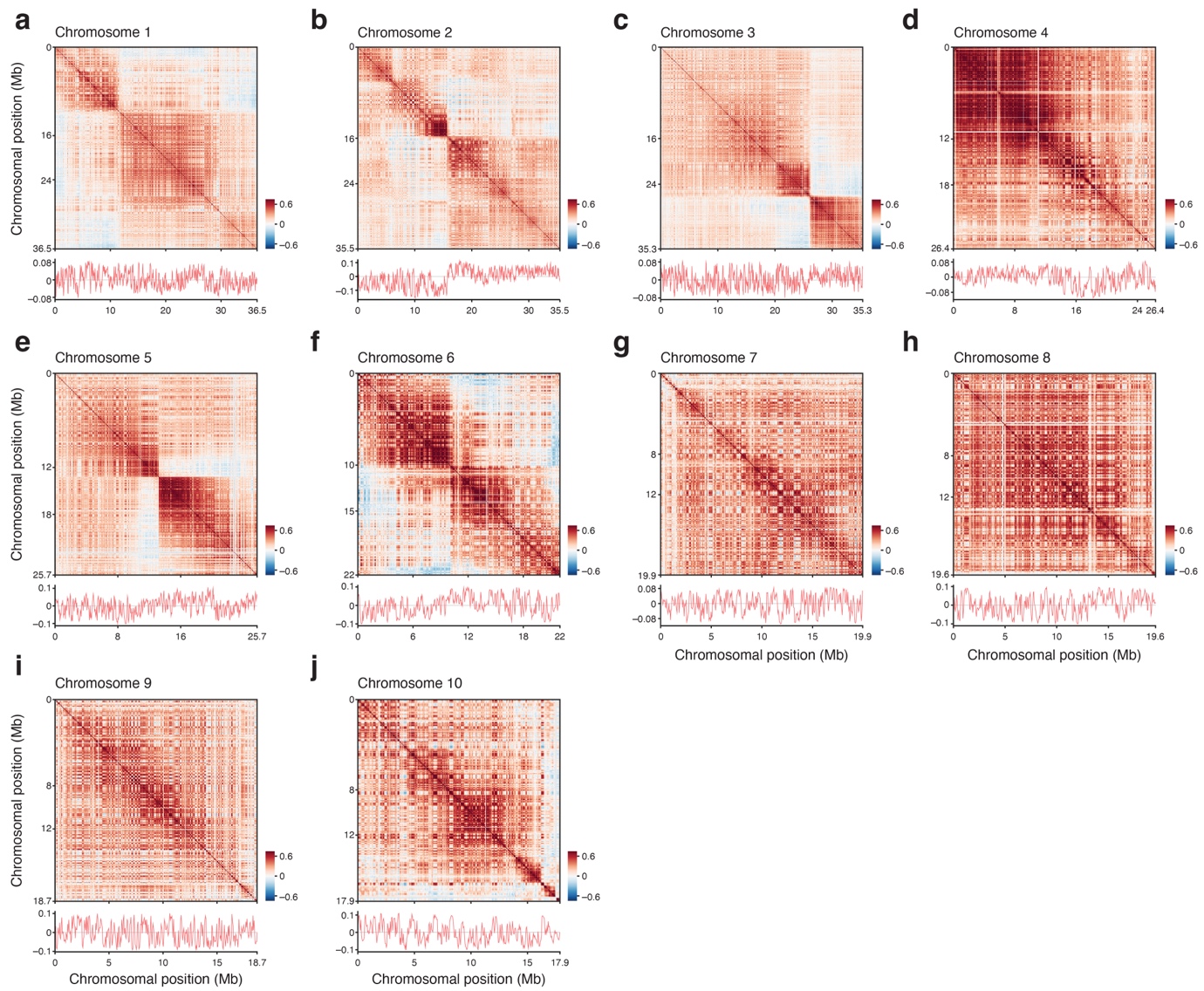

**Supplementary Figure 7 | Chromatin compartmentalisation in the adult of *C. teleta*.** (**a**–**j**) For each panel, a Pico-C contact heatmap for each chromosome is shown at the top. The bottom panel shows a line plot of eigenvalues along the chromosome.

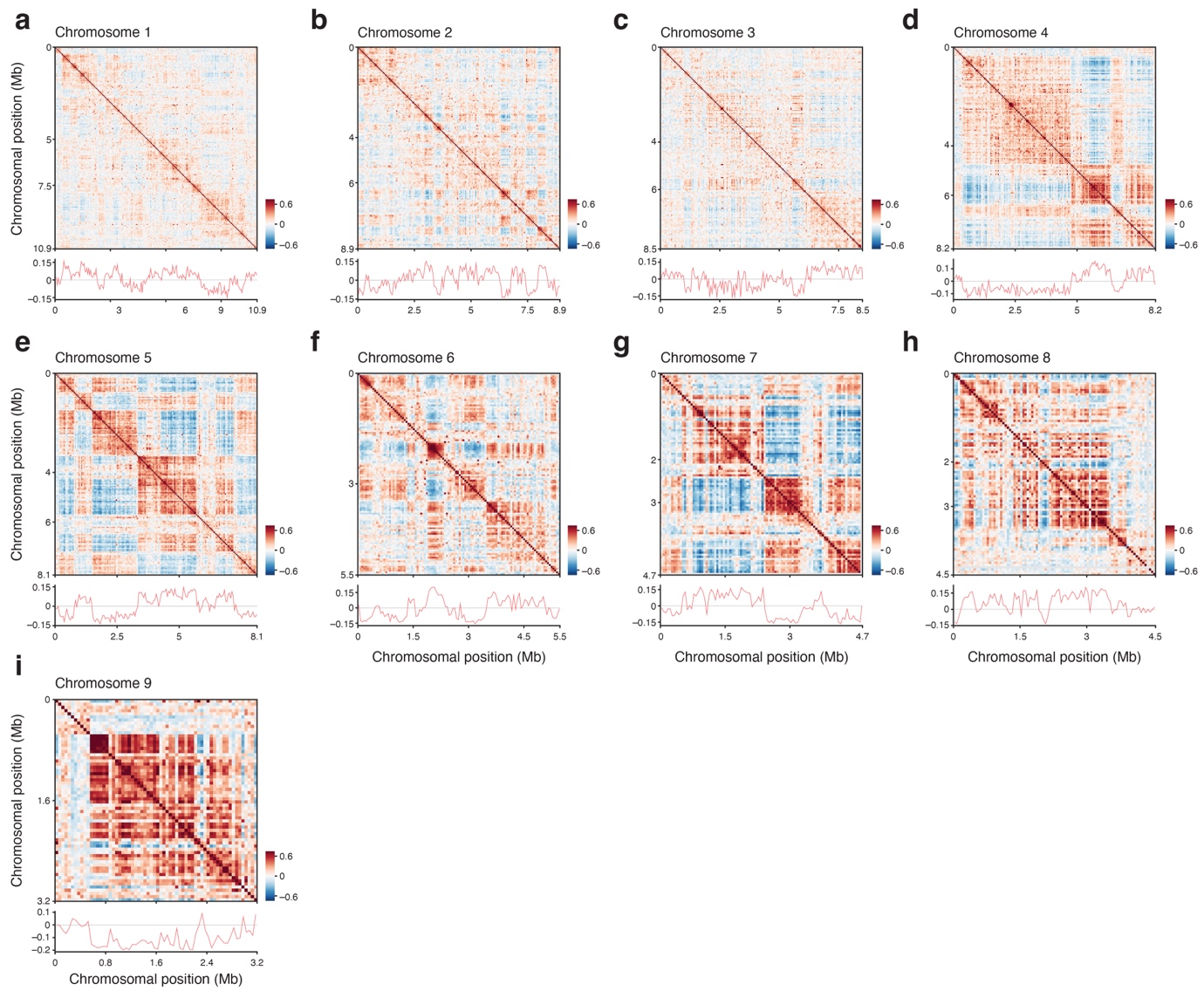

**Supplementary Figure 8 | Chromatin compartmentalisation in the adult of *D. gyrociliatus*.** (**a**–**i**) For each panel, a Pico-C contact heatmap for each chromosome is shown at the top. The bottom panel shows a line plot of eigenvalues along the chromosome.

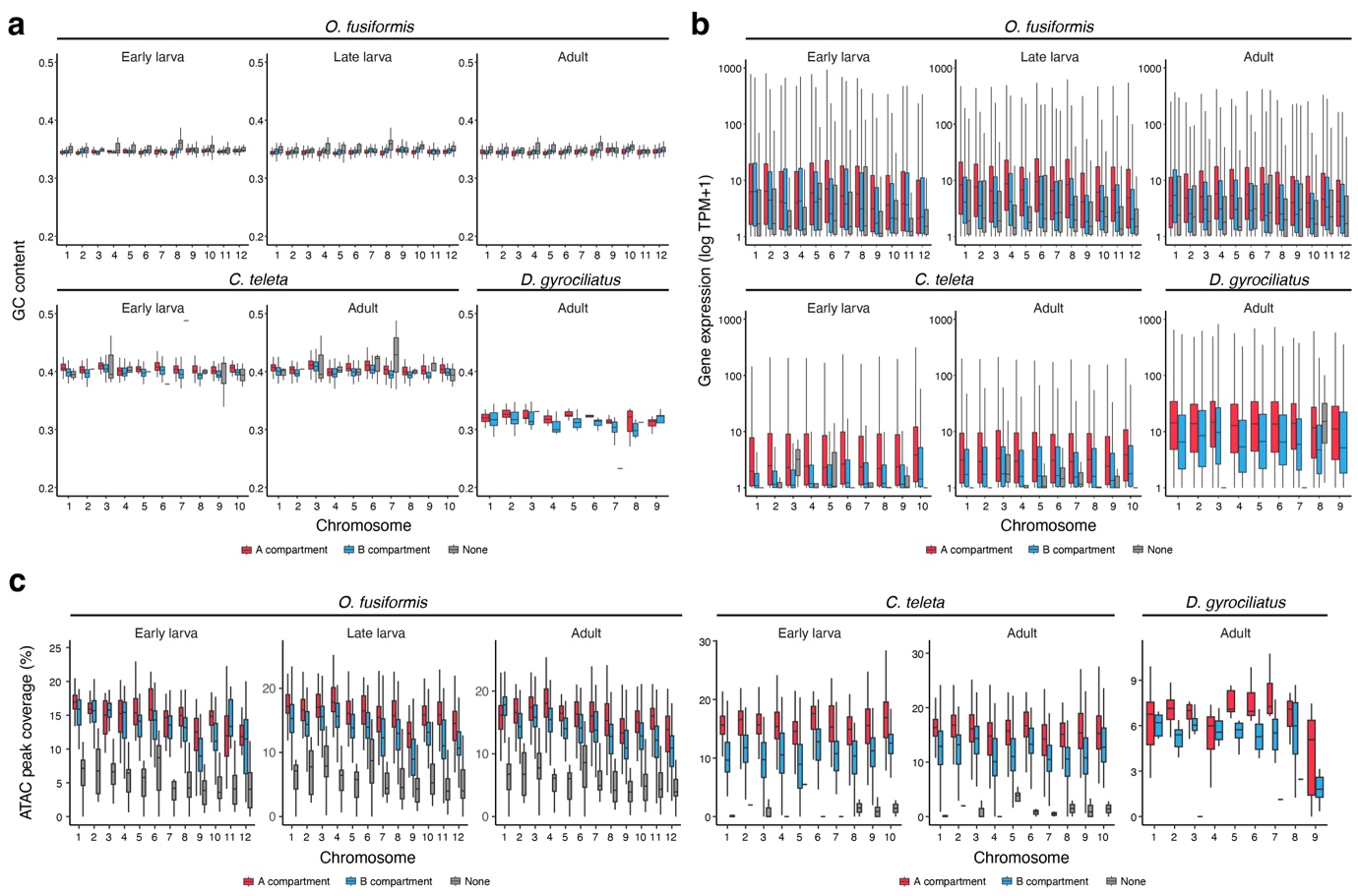

**Supplementary Figure 9 | Chromatin compartments and gene expression in annelids.** (**a**–**c**) Box plots showing the GC content (**a**), gene expression (**b**) and ATAC-seq peak coverage (**c**) in compartments A and B, and in regions not assigned to any compartment, per chromosome across the different stages and annelid species. For box plots, the centre line shows the median; the box shows the interquartile range (IQR); and the whiskers extend to the first or third quartile ± 1.5 × IQR.

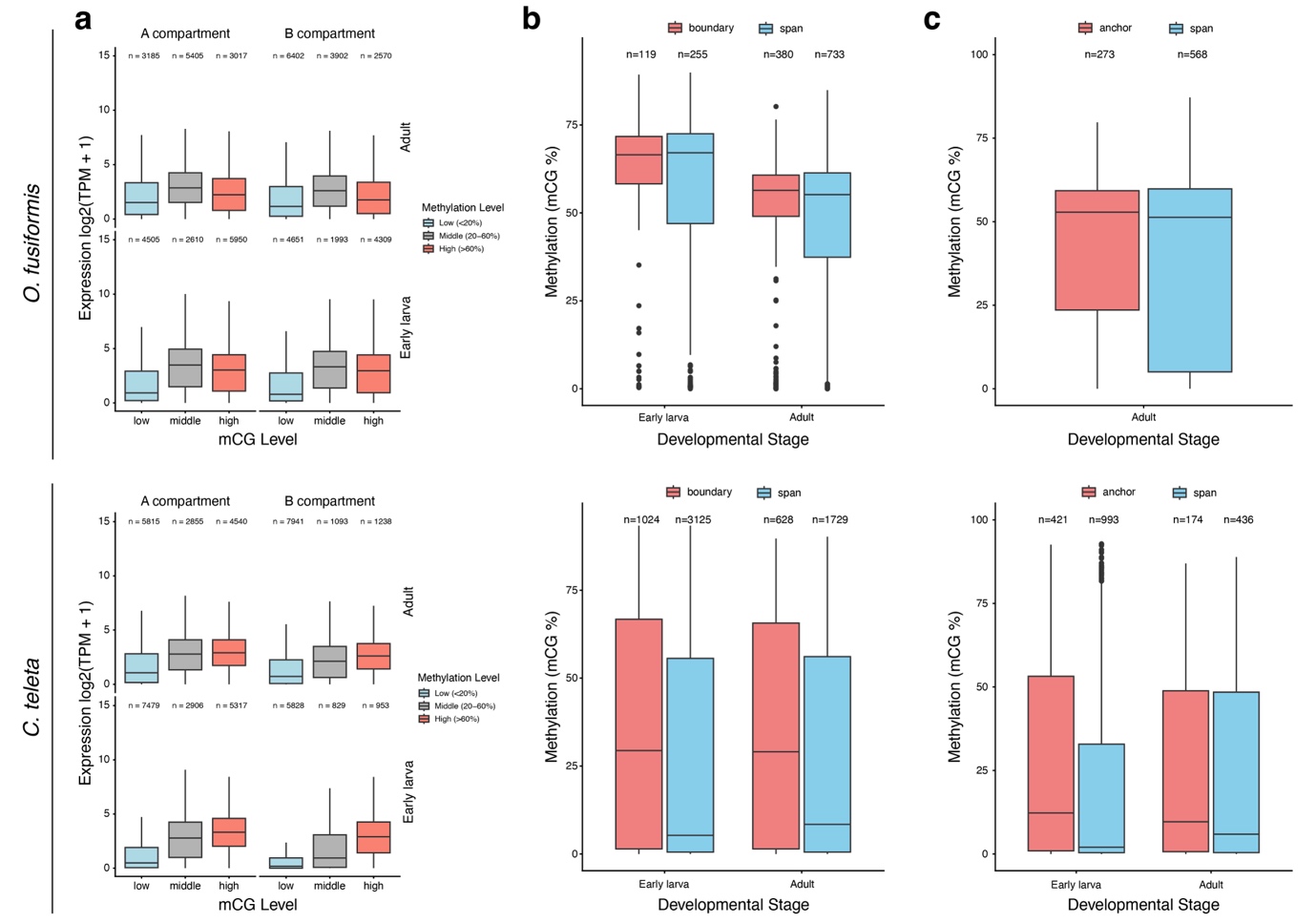

**Supplementary Figure 10 | 3D chromatin architecture and DNA methylation in *O. fusiformis* and *C. teleta*.** (**a**) Box plots showing gene expression levels in A and B compartments, grouped by gene body methylation (low, medium, and high). (**b**, **c**) Box plots showing gene body methylation levels at boundaries and TAD spans (**b**) and at anchors and chromatin loop spans (**c**) in *O. fusiformis* (top) and *C. teleta* (bottom). For box plots, the centre line shows the median; the box shows the interquartile range (IQR); and the whiskers extend to the first or third quartile ± 1.5 × IQR.

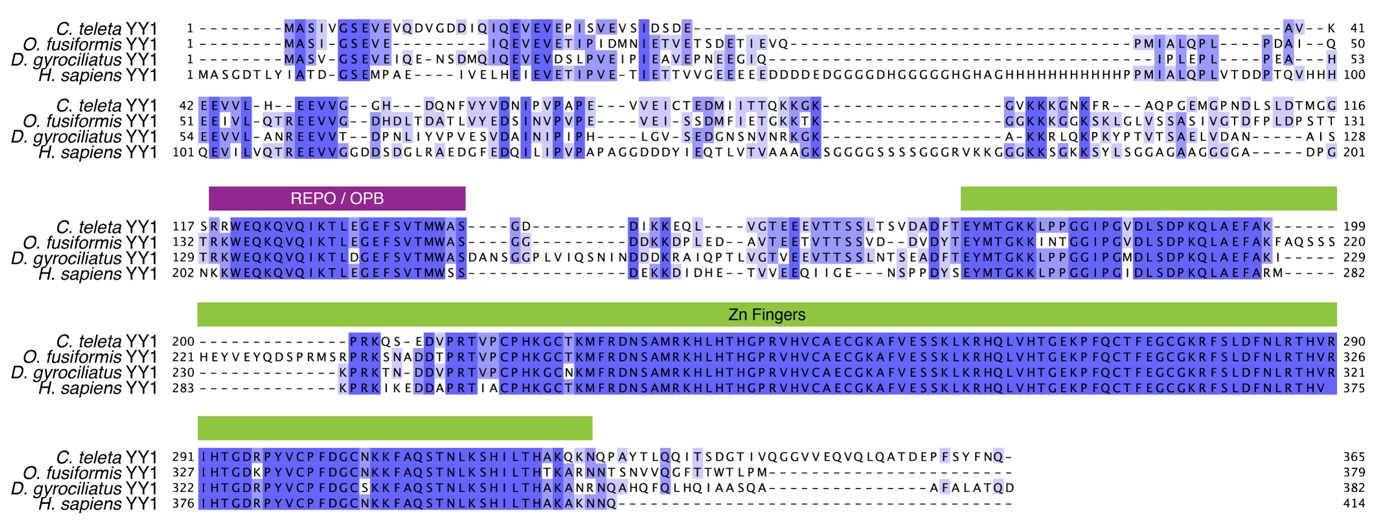

**Supplementary Figure 11 | YY1 evolution in annelids.** Multiple full-sequence alignment of the YY1 proteins from the annelids *C. teleta*, *O. fusiformis* and *D. gyrociliatus*, and the human orthologue. The REPO/OPB and Zn finger domains are highlighted with purple and green boxes, respectively. Sequence similarity per position is indicated in blue (dark blue indicates high similarity, light blue and white indicate low or no similarity).

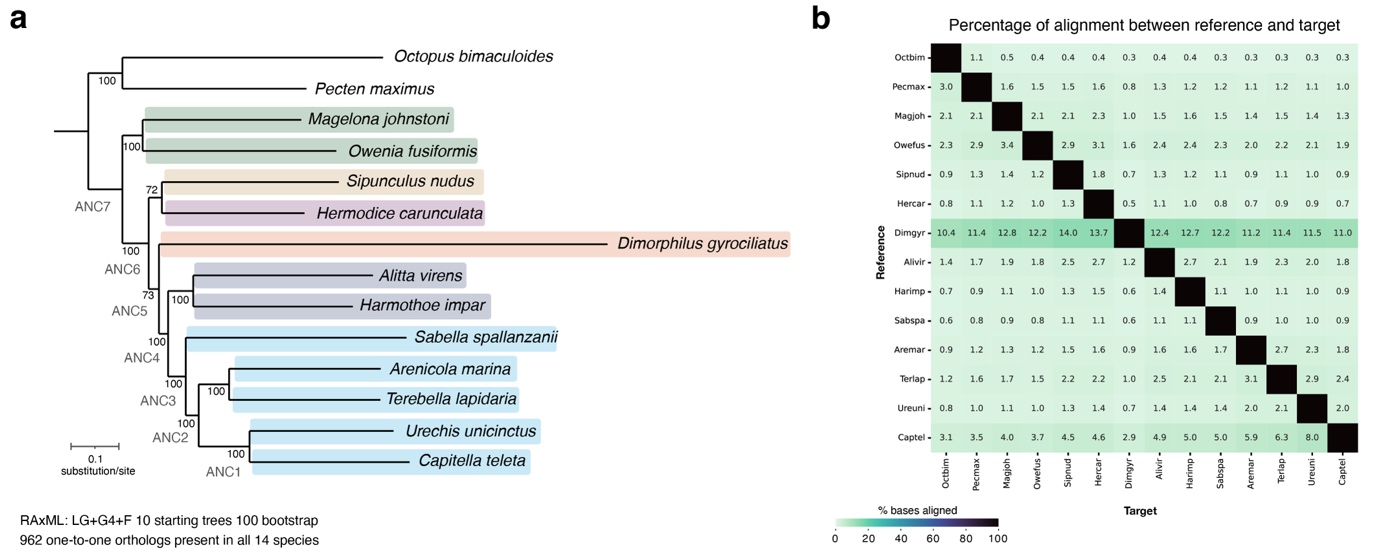

**Supplementary Figure 12 | Multi-species full-genome alignment.** (**a**) Phylogenetic tree based on full-genome alignments of the species included in the study, used to infer sequence conservation scores and phylogenetic origin. (**b**) Heatmap showing the percentage of bases aligned between pairs of species.

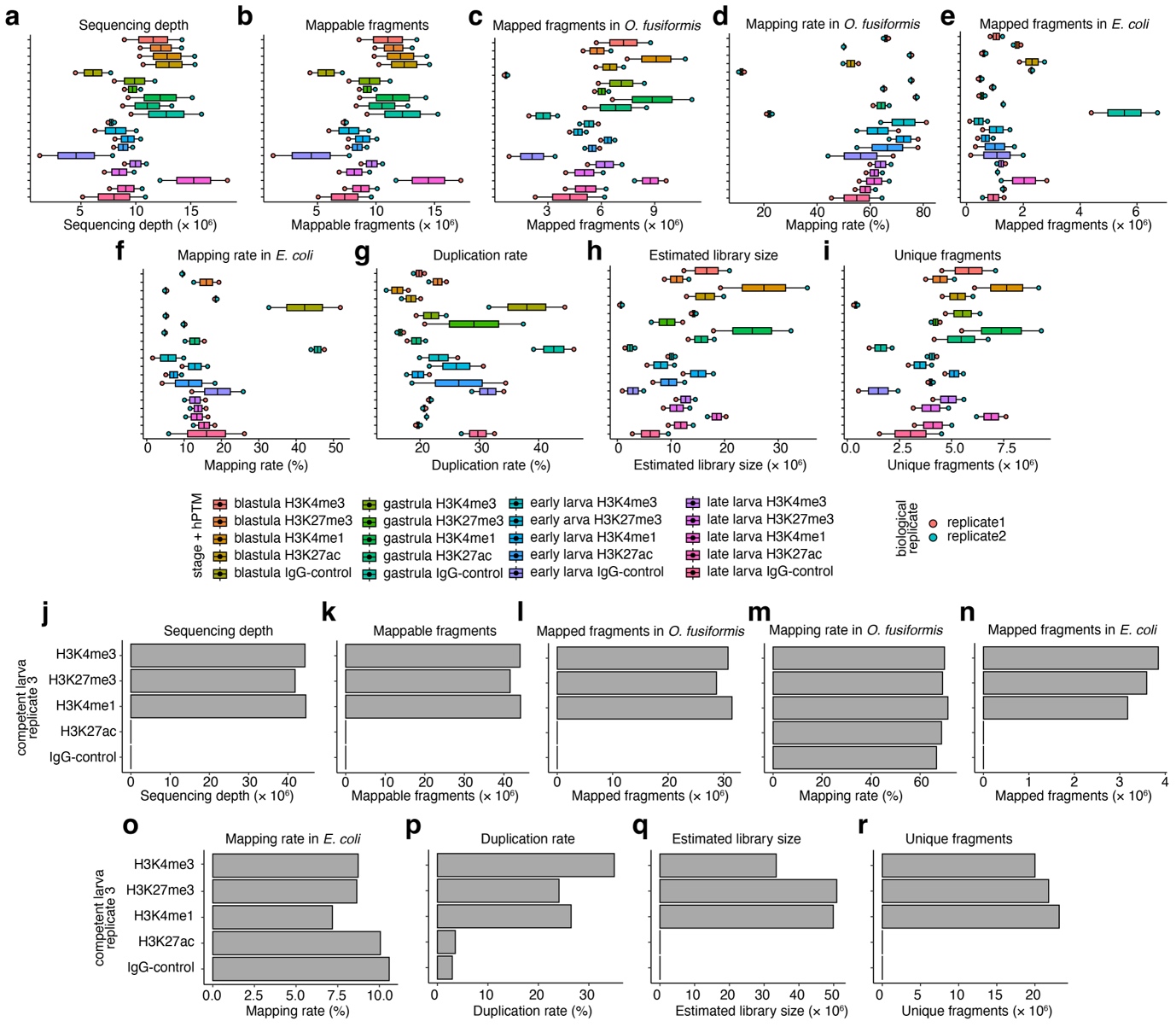

**Supplementary Figure 13 | CUT&Tag library statistics.** (**a**–**i**) Box plots showing sequencing depth (**a**), mappable fragments (**b**), mapped fragments (**c**), mapping rate (**d**) in *O. fusiformis*, mapped fragments (**e**), mapping rate (**f**) in *E. coli*, percentage of duplicates (**g**), estimated library size (**h**), and unique fragments (**i**) across the different CUT&Tag libraries. For box plots, the centre line shows the median; the box shows the interquartile range (IQR); and the whiskers extend to the first or third quartile ± 1.5 × IQR. (**j**–**r**) Bar plots showing similar statistics per histone modification.

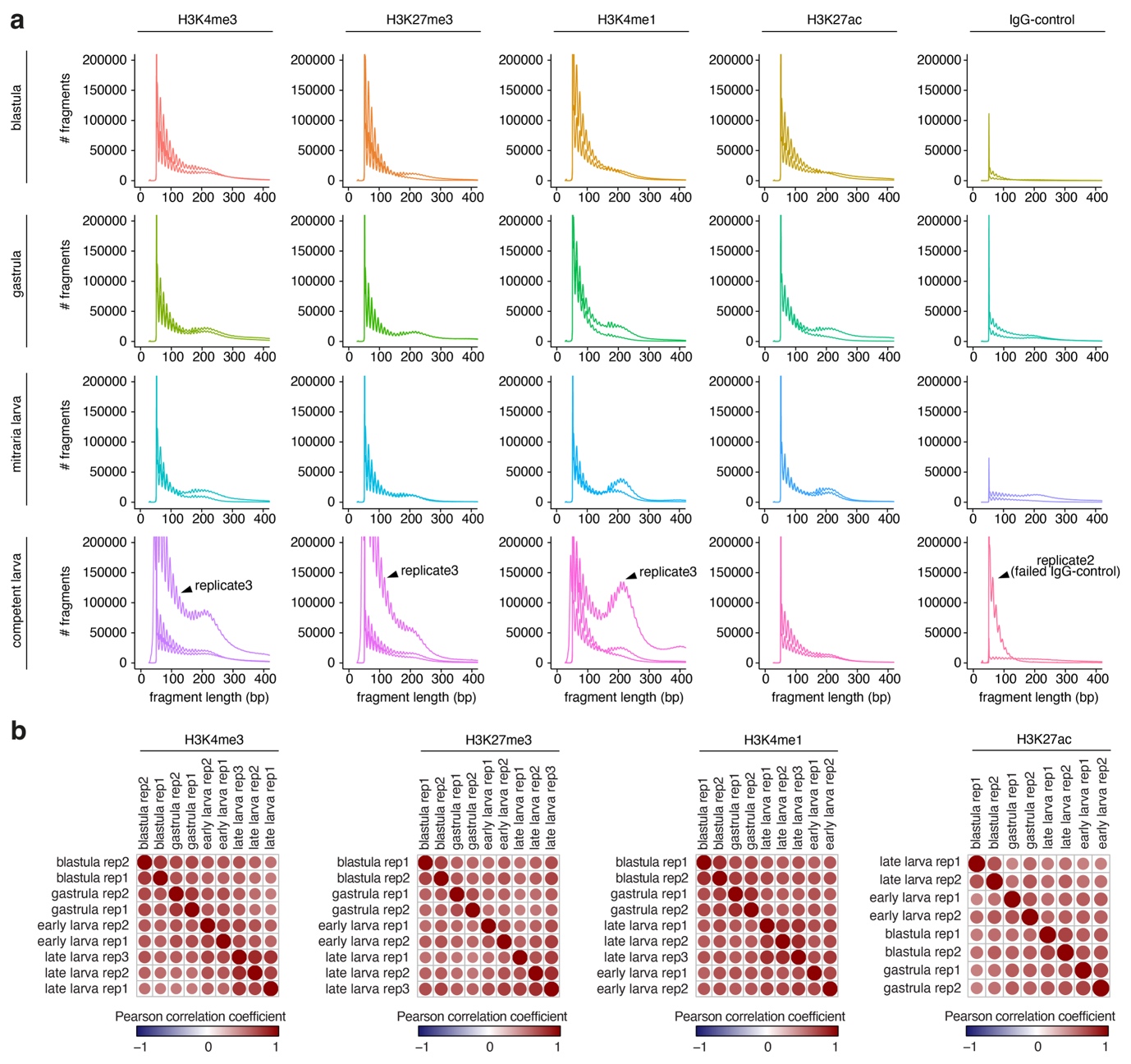

**Supplementary Figure 14 | CUT&Tag insert size distributions.** (**a**) Insert size distributions of CUT&Tag libraries from *O. fusiformis*. From left to right: H3K4me3, H3K27me3, H3K4me1, and H3K27ac libraries, and the IgG-negative control library. From top to bottom: blastula, gastrula, early larva, and late larva stages. Arrows point to the distributions of replicate 3 samples from the competent larva stage (H3K4me3, H3K27me3, and H3K4me1), which show a significantly higher number of fragments across almost every fragment length, simply due to higher sequencing depth. Note that the IgG-negative control sample from replicate 2 failed, as it contained a larger number of DNA fragments than any other control library. (**b**) Bubble plots showing Pearson correlation coefficients between replicates for each histone modification.

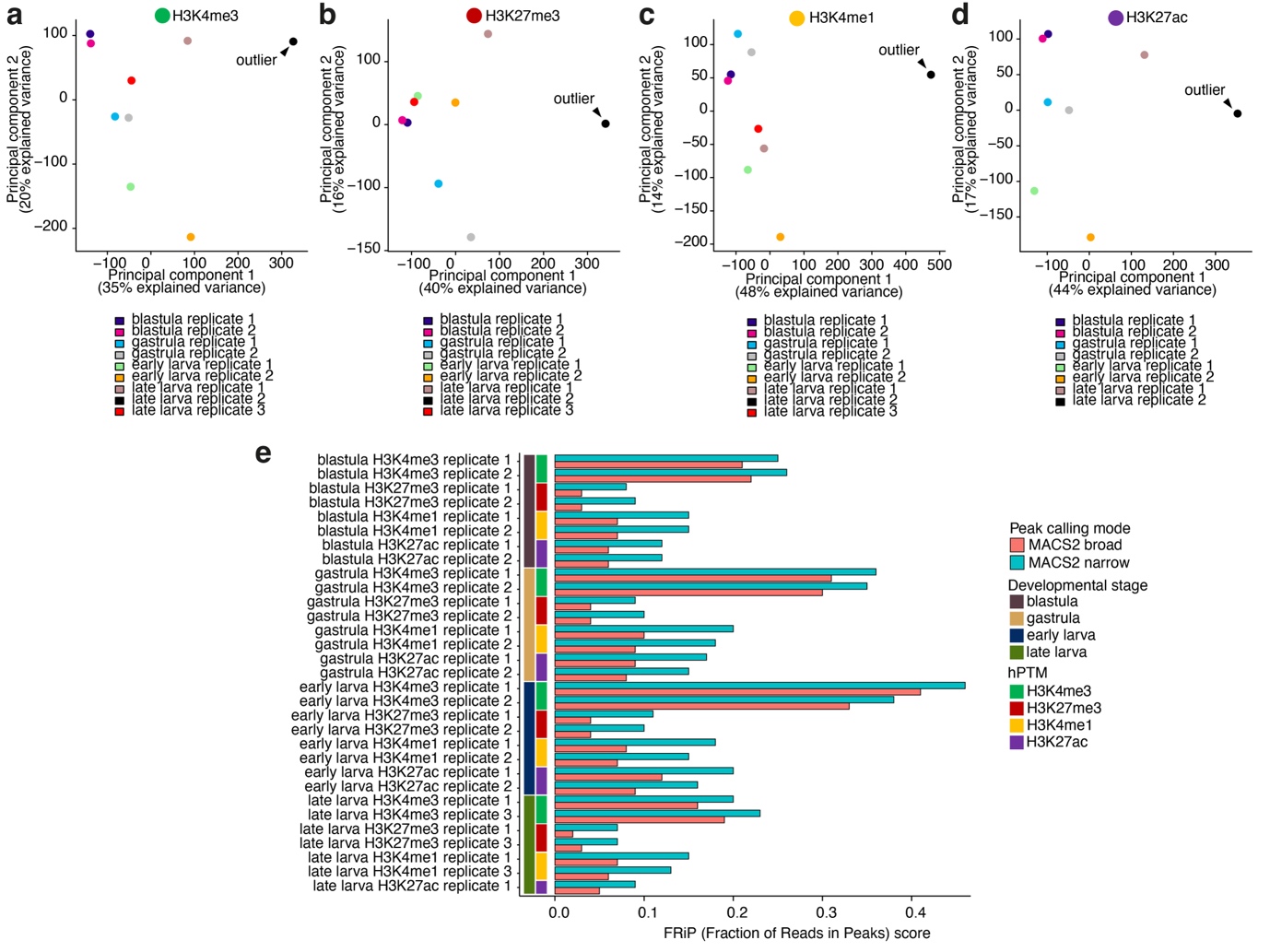

**Supplementary Figure 15 | Reproducibility assessment and peak calling benchmarking.** (**a**–**d**) Principal component analysis of CUT&Tag score dynamics across the hPTM-specific DiffBind consensus peak sets, following peak calling with MACS2 in narrow mode. The two biological replicates for the blastula, gastrula, and early larva stages are well correlated across all hPTMs, whereas this is not the case for the late larva stage. After including a third replicate, we found that replicate 2 of the late larval stage not only failed (due to a failed IgG-negative control library) but also served as an outlier for every hPTM. We therefore discarded replicate 2 for the late larva stage and continued our analysis with replicates 1 and 3 for H3K4me3, H3K27me3, and H3K4me1, and with replicate 1 only for H3K27ac. (**e**) FRiP (Fraction of Reads in Peaks) scores for all CUT&Tag samples in the hPTM-specific DiffBind consensus peak sets, following peak calling with MACS2 in either narrow (blue) or broad mode (pink). Narrow peak calling was the optimal mode across all 31 samples.

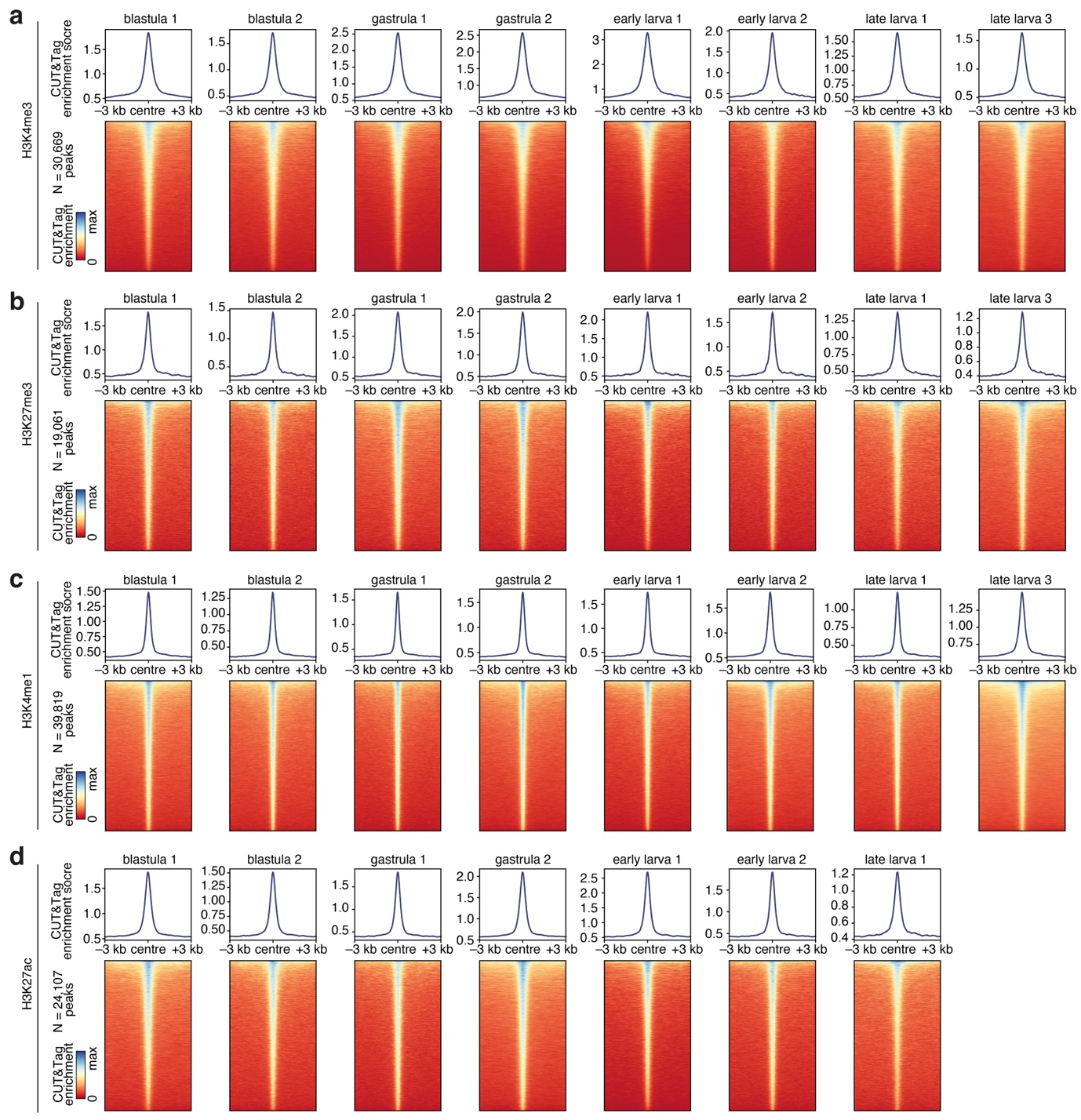

**Supplementary Figure 16 | CUT&Tag enrichment around hPTM peaks.** (**a**–**d**) CUT&Tag enrichment profiles (top) and heatmaps (bottom) around the peak centres (± 3 kb) of the hPTM-specific DiffBind consensus peak sets obtained by peak calling with MACS2 in narrow mode, for H3K4me3 (**a**), H3K27me3 (**b**), H3K4me1 (**c**), and H3K27ac (**d**). For each hPTM, from left to right: blastula replicates 1 and 2, gastrula replicates 1 and 2, early larva replicates 1 and 2, and late larva replicates 1 and 3 (except for H3K27ac, which has only replicate 1). The CUT&Tag enrichment score scale is sample-specific. All samples show at least 3-fold enrichment at the peaks.

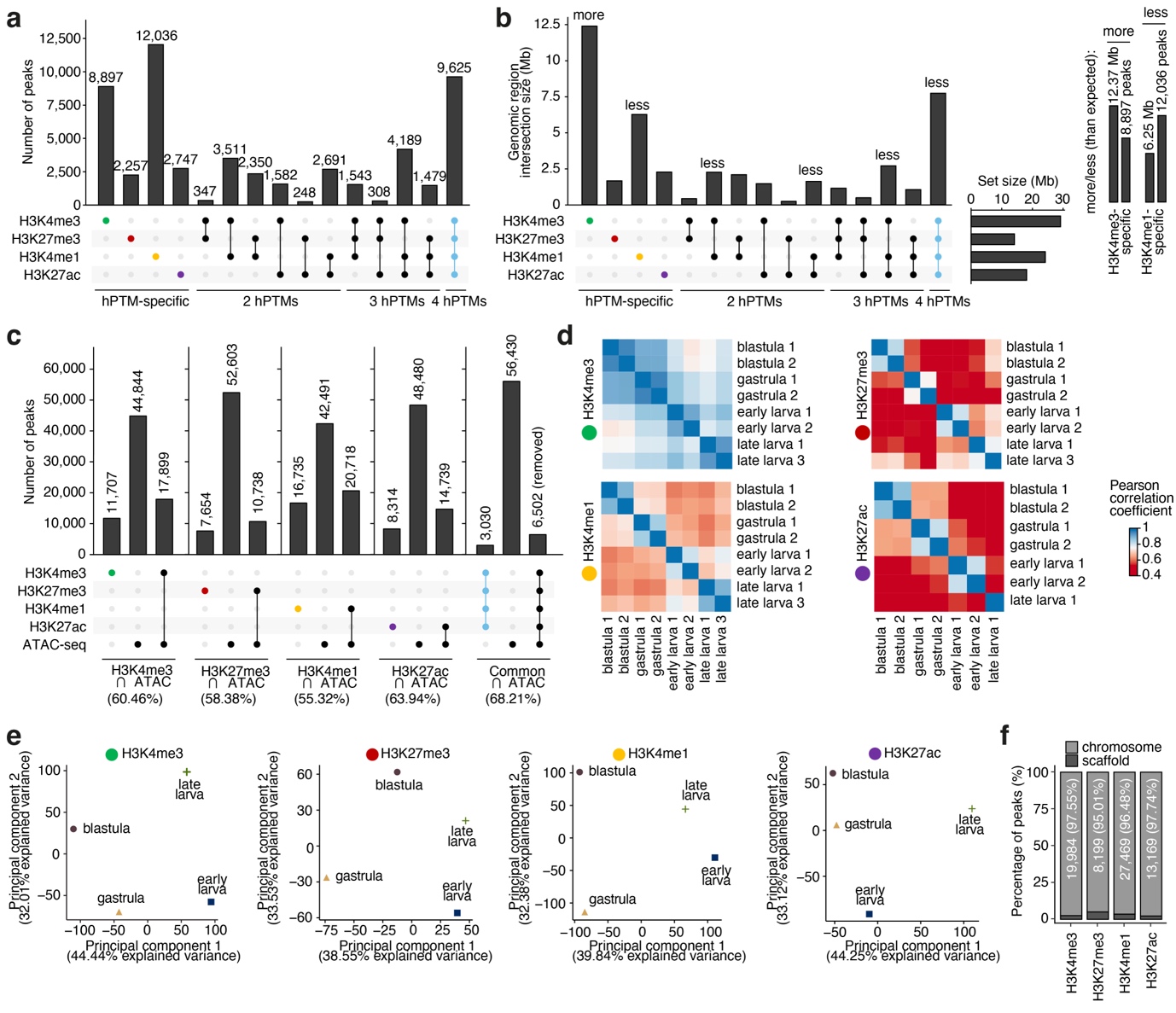

**Supplementary Figure 17 | Correlation and developmental dynamics of histone marks.** (**a**, **b**) UpSet plot of the four CUT&Tag hPTM-specific DiffBind consensus peak sets. Overlap is shown as the number of peaks (**a**) and as genomic distances (**b**; in Mb) between peak sets. hPTM-specific regions are shown to the right, in the colours of each hPTM. Constitutive peaks are shown to the left, in blue. In (**b**), the inset on the right illustrates how regions marked with “more” or “less” refer to those that are larger or smaller than expected, given the number of peaks in each intersection. (**c**) UpSet plot of each of the four CUT&Tag hPTM-specific DiffBind consensus peak sets, the set of constitutive peaks, and the ATAC-seq DiffBind consensus peak set. Numbers above each bar represent the number of peaks. Numbers in parentheses denote the percentage of CUT&Tag peaks overlapping with the ATAC-seq peak set. Constitutive peaks are those that show the highest overlap with ATAC peaks. (**d**) Correlation matrices of normalised CUT&Tag scores in the hPTM-specific DiffBind consensus peak sets across the development of *O. fusiformis*, for H3K4me3, H3K27me3, H3K4me1, and H3K27ac. The colour scale denotes the Pearson correlation coefficient. (**e**) Principal component analysis of normalised CUT&Tag scores in the hPTM-specific DiffBind consensus peak sets, averaged by developmental stage, for H3K4me3, H3K27me3, H3K4me1, and H3K27ac. (**f**) Percentage of CUT&Tag consensus peaks present in the 12 chromosomes of the *O. fusiformis* assembly (89.19% of the assembly, 10.81% in scaffolds). Peaks from all hPTM-specific consensus peak sets are overrepresented on the 12 chromosomes (minimum: 94.59% for H3K27me3) and underrepresented in scaffolds (maximum: 5.41% for H3K27me3).

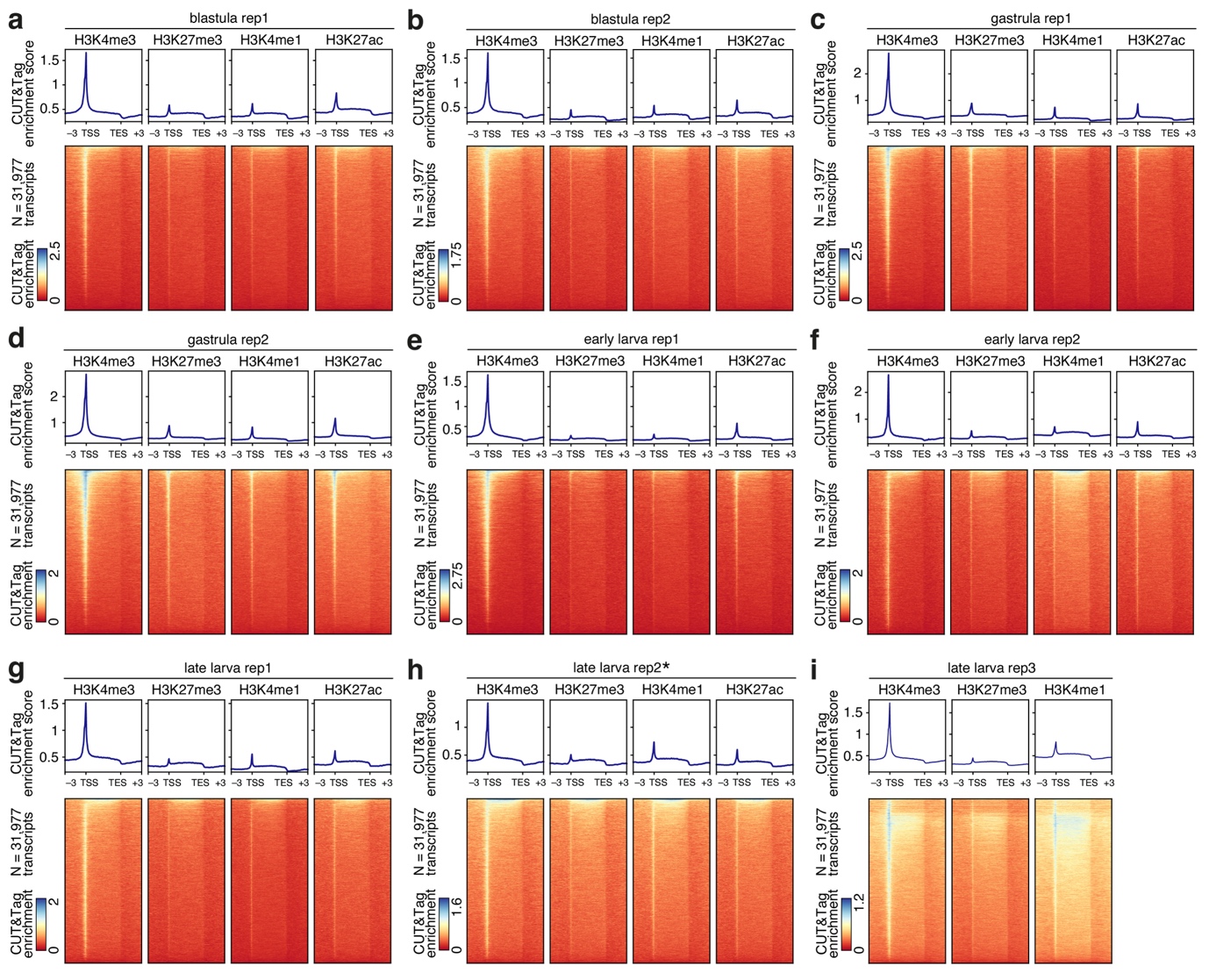

**Supplementary Figure 18 | Histone mark enrichment in *O. fusiformis* genes.** (**a**–**i**) CUT&Tag enrichment meta-gene profiles (top) and heatmaps (bottom) around *O. fusiformis* gene models for each biological replicate at the blastula (**a** and **b**), gastrula (**c** and **d**), early larva (**e** and **f**), and late larva (**g**–**i**) stages. For each replicate, enrichment is shown from left to right for H3K4me3, H3K27me3, H3K4me1, and H3K27ac. The CUT&Tag enrichment score scale is replicate-specific. Distances are in kilobases (kb). TSS: transcription start site; TES: transcription end site.

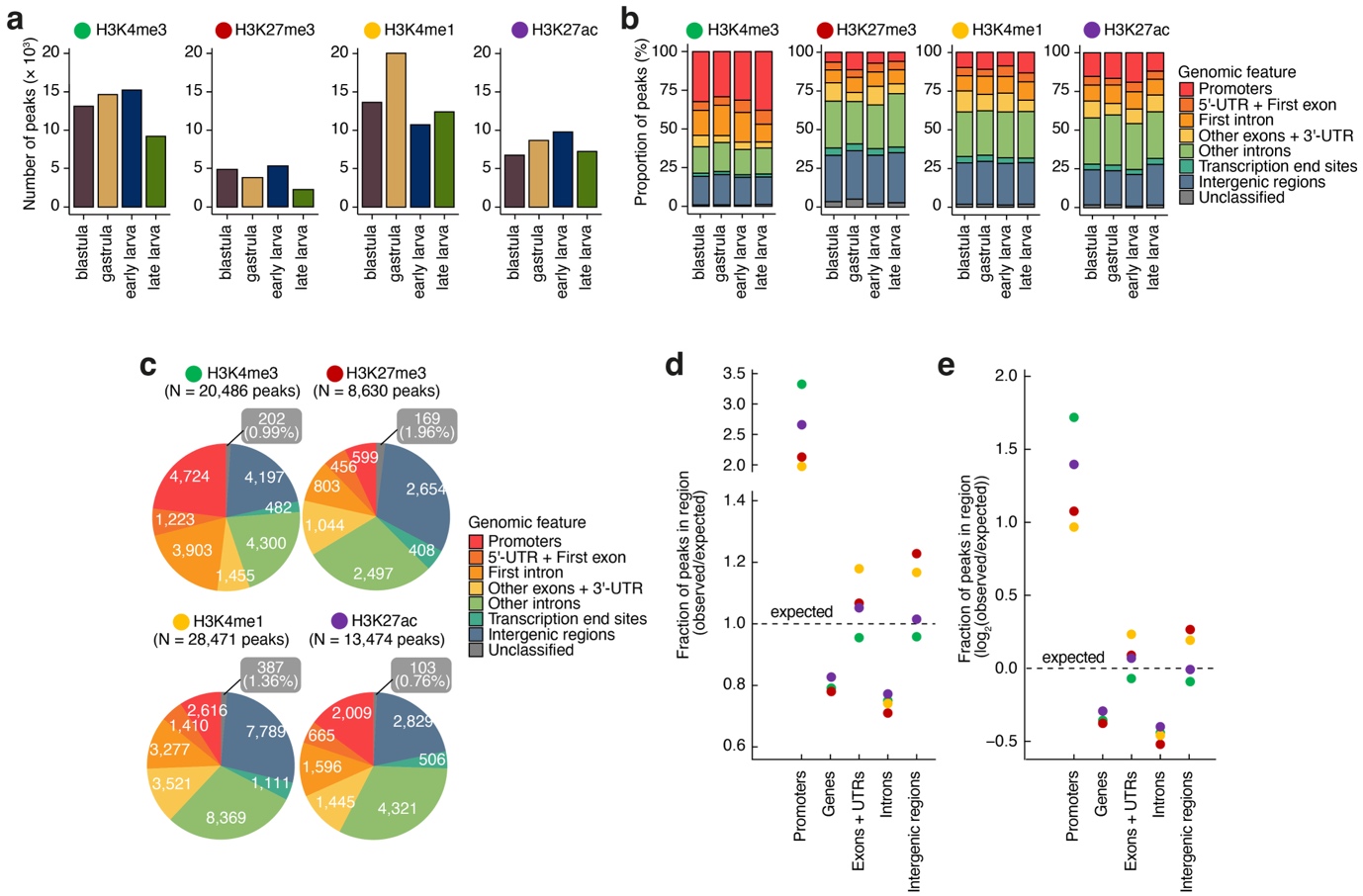

**Supplementary Figure 19 | Developmental dynamics of histone marks.** (**a**) Bar plots showing the number of peaks per developmental stage for H3K4me3 (left), H3K27me3 (centre left), H3K4me1 (centre right), and H3K27ac (right). (**b**) Bar plot showing the percentage of peaks called per developmental stage. (**c**) Genomic feature annotation of the CUT&Tag hPTM-specific consensus peak sets for H3K4me3 (left), H3K27me3 (centre left), H3K4me1 (centre right), and H3K27ac (right). Absolute numbers for each genomic feature are shown inside each section of the pie chart. For unclassified peaks, both absolute numbers and percentages (in parentheses) are shown. (**d**, **e**) Enrichment plots of peak location, showing the ratio of the observed to the expected (genome background) percentage of peaks (**d**) or the base 2 logarithm of this ratio. All marks are enriched in promoters and underrepresented in genes, both overall and in introns. Some hPTM-specific differences can be observed across exons, UTRs, and intergenic regions.

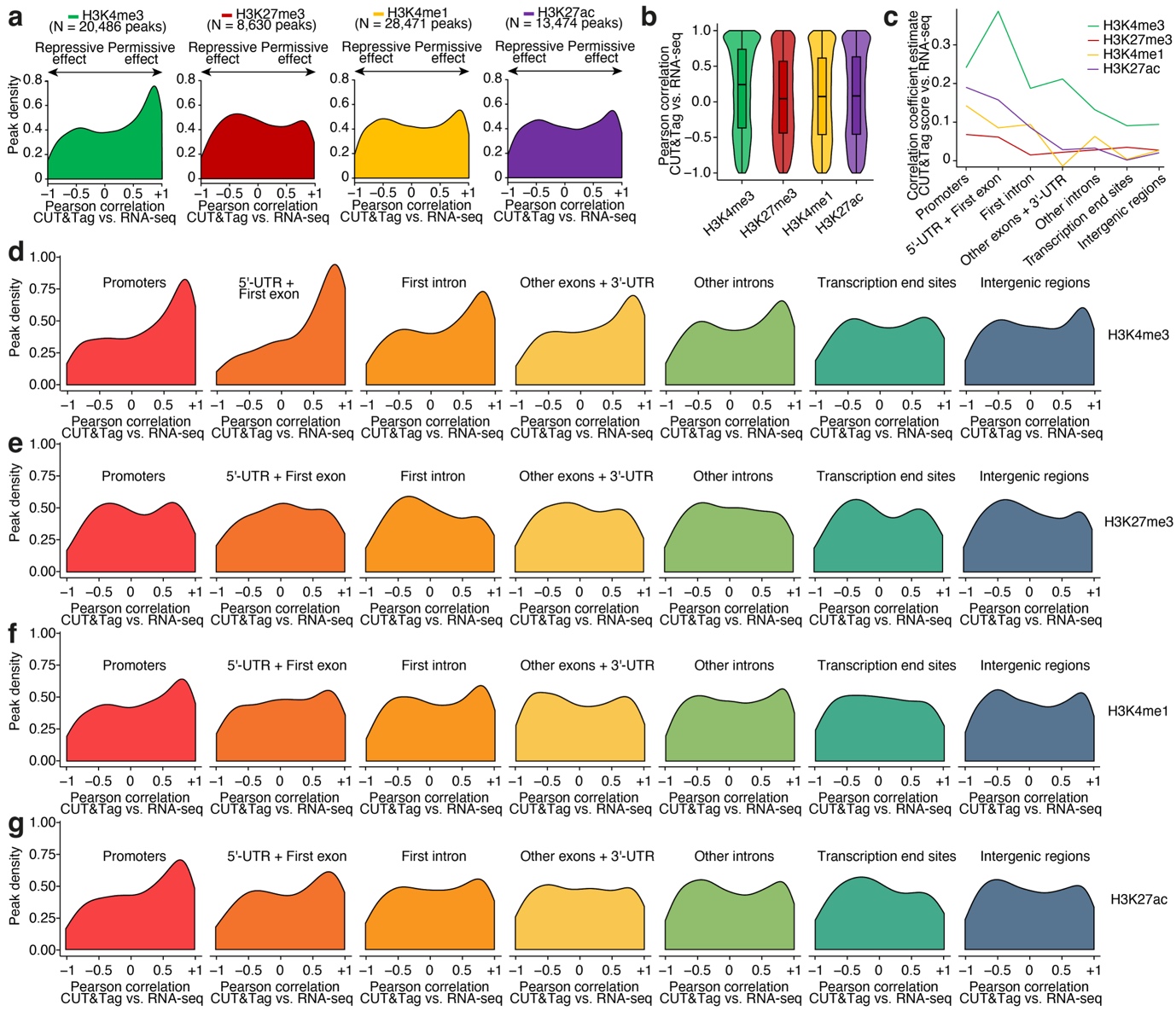

**Supplementary Figure 20 | Correlation between hPTM dynamics and gene expression dynamics.** (**a**) Density plots of the Pearson correlation coefficient between the CUT&Tag score and the RNA-seq expression level of the nearest transcript to each CUT&Tag peak, for H3K4me3 (left), H3K27me3 (centre left), H3K4me1 (centre right), and H3K27ac (right). (**b**) Violin and box plots depicting the distributions shown in (**a**). For box plots, the centre line shows the median; the box shows the interquartile range (IQR); and the whiskers extend to the first or third quartile ± 1.5 × IQR. (**c**) Simplified interaction plot of a bivariate linear model with interaction (correlation~hPTM*feature). (**d**–**g**) Density plots of the Pearson correlation coefficient between the CUT&Tag score and the RNA-seq expression level of the nearest transcript to each CUT&Tag peak, classified by genomic feature, for H3K4me3 (**d**), H3K27me3 (**e**), H3K4me1 (**f**), and H3K27ac (**g**). Promoter and 5’-UTR + First exon peaks of H3K4me3 are the most positively correlated with transcription.
